# Targeting EZH2 Increases Therapeutic Efficacy of Check-Point Blockade in Models of Prostate Cancer

**DOI:** 10.1101/730135

**Authors:** Anjali V. Sheahan, Katherine L. Morel, Deborah L. Burkhart, Sylvan C. Baca, David P. Labbé, Kevin Roehle, Max Heckler, Carla Calagua, Huihui Ye, Phillip Galbo, Sukanya Panja, Antonina Mitrofanova, Anis A. Hamid, Adam S. Kibel, Atish D. Choudhury, Mark M. Pomerantz, Matthew L. Freedman, Christopher J. Sweeney, Stephanie K. Dougan, Adam G. Sowalsky, Massimo Loda, Brian M. Olson, Leigh Ellis

**Affiliations:** Department of Oncologic Pathology, Dana-Farber Cancer Institute, Boston MA; Department of Medical Oncology, Dana-Farber Cancer Institute, Boston MA; Division of Urology, Department of Surgery, McGill University and Research Institute of the McGill University Health Centre, Montréal, Québec, Canada; Department of Cancer Immunology and Virology, Dana-Farber Cancer Institute, Boston MA; Department of Health Informatics, Rutgers School of Health Professionals, Newark NJ; Department of Medicine, Hematology-Oncology Division, Beth Israel Deaconess Medical Center, Harvard Medical School, Boston MA; Department of Pathology, University of California at Los Angeles, Los Angeles CA; Department of Cell Biology, Albert Einstein College of Medicine, Bronx NY; Rutgers Cancer Institute of New Jersey, Rutgers, The State University of New Jersey, New Brunswick NJ; Department of Urology, Brigham and Women’s Hospital, Harvard Medical School, Boston MA; Laboratory of Genitourinary Cancer Pathogenesis, Center for Cancer Research, National Cancer Institute, NIH, Bethesda MD; Department of Pathology and Laboratory Medicine, Weill Cornell Medicine, New York, NY; Departments of Hematology and Medical Oncology and Urology, Emory University School of Medicine, Atlanta GA; Department of Pathology, Brigham and Women’s Hospital, Harvard Medical School, Boston MA; The Broad Institute, Cambridge MA

**Keywords:** EZH2, Prostate Cancer, Epigenetics, Targeted Therapy, Immunotherapy, PD-L1, PD-1

## Abstract

Prostate cancers are considered immunologically ‘cold’ tumors given the very few patients who respond to checkpoint inhibitor therapy (CPI). Recently, enrichment of interferon (IFN) response genes predicts a favorable response to CPI across various disease sites. The enhancer of zeste homolog-2 (EZH2) is over-expressed in prostate cancer and is known to negatively regulate IFN response genes. Here, we demonstrate that inhibition of EZH2 catalytic activity in prostate cancer models derepresses expression of double-strand RNA (dsRNA), associated with upregulation of genes involved in antigen presentation, Th-1 chemokine signaling, and interferon (IFN) response, including PD-L1. Similarly, application of a novel EZH2 derived gene signature to human prostate sample analysis indicated an inverse correlation between tumor EZH2 activity/expression with T-cell inflamed and IFN gene signatures and PD-L1 expression. EZH2 inhibition combined with PD-1 CPI significantly enhances antitumor response that is dependent on up-regulation of tumor PD-L1 expression. Further, combination therapy significantly increases intratumoral trafficking of activated CD8+ T-cells and M1 tumor associated macrophages (TAMs) with concurrent loss of M2 TAMs. Our study identifies EZH2 as a potent inhibitor of antitumor immunity and responsiveness to CPI. This data suggests EZH2 inhibition as a novel therapeutic direction to enhance prostate cancer response to PD-1 CPI.

## Main Text

Prostate cancer (PCa) is the currently the most commonly diagnosed non-cutaneous malignancy and the second most common cause of cancer death amongst men in the United States (*1*). Unfortunately, metastatic castration-resistant prostate cancer (mCRPC) still remains incurable, despite recent advances in therapy options for these patients. While checkpoint inhibition (CPI) can generate dramatic responses in about 15-20% of patients with a number of cancer types including melanoma, kidney and bladder cancer, this occurs in approximately 5% of PCa patients. Resistance towards CPI in PCa patients is thought to be related to low tumor immunogenicity and an immunosuppressive tumor microenvironment.

The enhancer of zeste homologue 2 (EZH2) is the methyltransferase catalytic subunit of the polycomb repressive complex 2 (PRC2) that trimethylates lysine 27 of histone H3 (H3K27me3) to promote transcriptional repression (*2*). Increased expression and activity of EZH2 is an important contributor to PCa initiation and progression (*3, 4*). EZH2 can negatively regulate interferon (IFN) response genes, Th-1 type chemokines, and MHC expression in multiple tumor cell types (*5, 6*). Dysfunction of epigenetic regulation within a cancer cell including effects mediated by EZH2, DNA methytransferases (DNMT), histone deacetylases (HDAC), BET bromodomains, and lysine specific demethylase 1 (LSD1) have proven to be critical mediators of acquired tumor immune escape. Subsequent inhibition of these epigenetic mechanisms results in increased tumor immunity and successful combination with CPI in preclinical cancer models (*5, 7–14*). Importantly, recent data from a phase Ib/II clinical trial, ENC0RE-601 (NCT02437136), illustrated the power of epigenetic therapy to restore partial sensitivity in melanoma patients who had progressed on an inhibitor of PD-1 (*15*). However, targeting epigenetic mechanisms, especially those mediated by EZH2, have not been tested for their ability to induce response to CPI in PCa.

Using 3-dimensional tumor organoids derived from a genetically engineered mouse model of PCa (GEMM) that expresses oncogenic *cMYC, Ezh2* floxed alleles (*16*), and an inducible *Cre* recombinase driven by the prostate specific antigen promoter (*17*) (EMC mouse, fig. S1), we demonstrated that chemical or genetic inhibition of EZH2 catalytic activity resulted in diminished organoid growth (Fig. 1A and 1D), accompanied by significant decrease in DNA synthesis and H3K27me3 (Fig. 1B and 1E). Independent gene set enrichment analysis (GSEA) revealed a significant enrichment of type I (IFN∝) and II (IFNγ) gene signatures (Fig. 1G, fig. S1). The enrichment of IFN response genes following EZH2 inhibition had been previously demonstrated to be exclusive to *cMYC* over-expressing PCa models (*14*). However, we didn’t observe enrichment of IFN response genes exclusive to MYC over-expression in human datasets (data not shown). Interrogation of the leading-edge genes related to IFN signaling from mouse PCa organoids revealed significant increases in expression of Th-1 chemokines (*Cxcl9, Cxcl10, Cxcl11*), antigen-presentation genes (*B2m, Tap1*), and IFNγ regulated genes (*Stat1, CD274/Pd-l1*) (table S1). The enrichment of IFN response genes was further corroborated when master regulator (MR) analysis using MARINa was applied to identify transcription factors (TFs) driving this pattern of gene expression. Our top TFs from chemically inhibited organoids included Stat1, Stat2 and Irf9 (Fig. 1H, table S2). These 3 proteins heterodimerize to form transcriptional machinery that drive IFN response gene expression (*18*).

**Fig 1.**
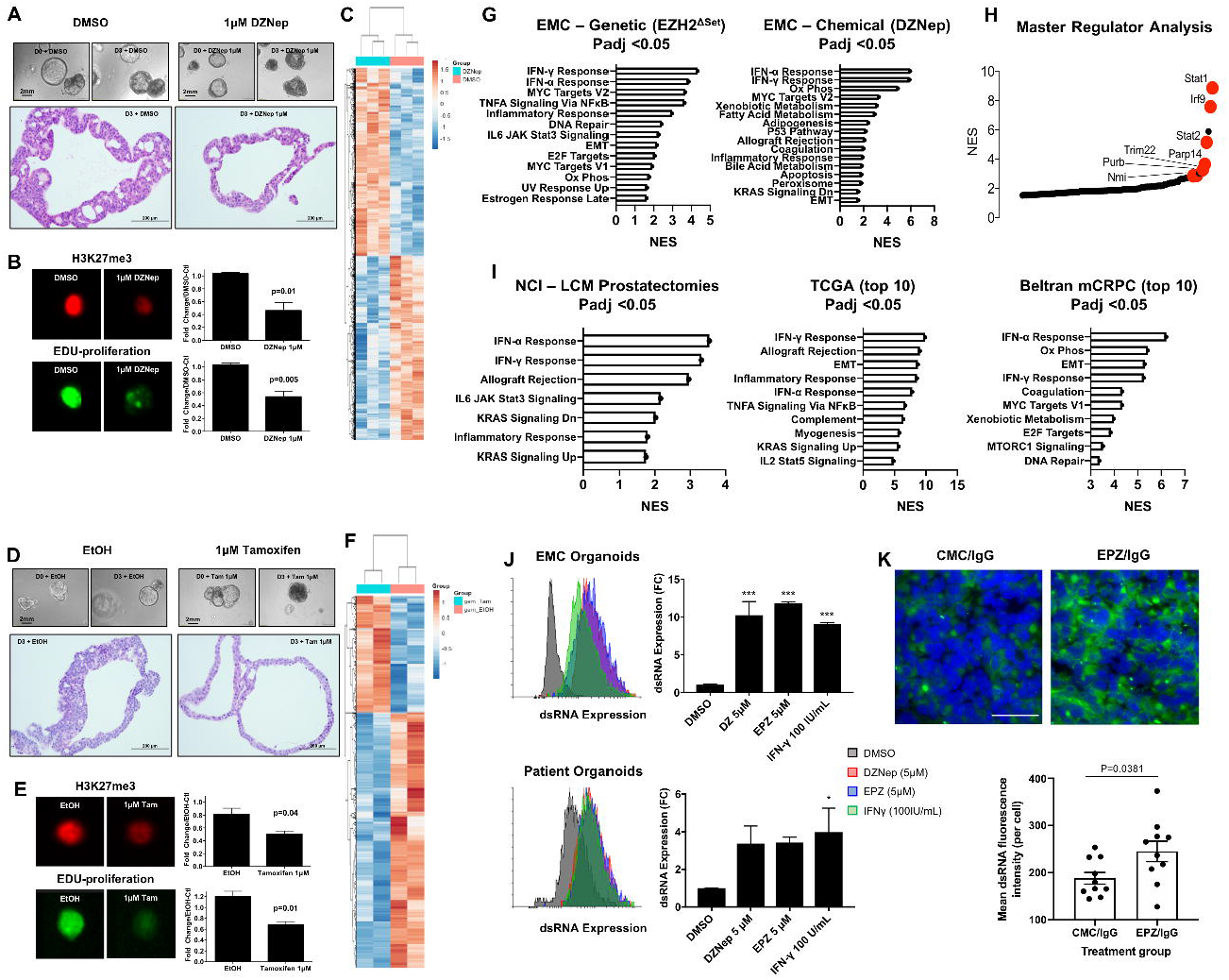
EZH2 inhibition induces viral mimicry in prostate cancer. EZH2 catalytic activity in EMC PCa mouse organoids was inhibited by **(A-C)** chemical or **(D-F)** genetic methods. Both chemical and genetic EZH2 inhibition significantly decreases H3K27me3 and DNA replication, and significantly alters gene expression. **(G)** Gene set enrichment analysis reveals enrichment of type I/II IFN gene signatures in mouse PCa organoids following EZH2 inhibition. **(H)** Master regulation analysis of RNA-seq data from 1C enriches for transcription factors that regulate type I/II IFN response genes. **(I)** Gene set enrichment analysis reveals enrichment of type I/II IFN gene signatures in human prostate cancer patients with lowest EZH2 activity. **(J)** Inhibition of EZH2 induces expression of dsRNA in mouse and human PCa organoids and **(K)** PCa tissue *in vivo*. *P<0.05, ***P<0.0001.

To determine if loss of EZH2 catalytic activity was associated with enrichment of IFN response genes in human PCa, a 29-gene EZH2 repression signature was derived using differentially expressed genes following chemical inhibition of EZH2 (Fig. 1C, table S3) and applied to independent human PCa RNAseq data sets (table S4). A similar EZH2 repression signature was previously reported (*19*) and while both signatures had no overlapping genes, they were significantly correlated with each other (fig. S2). Of importance, EZH2 activity was not altered because of changes in EZH2 expression (fig. S2). Upon quartile distribution of patients, differential gene expression between quartile 4 (lowest EZH2 activity) and quartile 1 (highest EZH2 activity) validated our *in vitro* murine results by indicating patients with lowest EZH2 activity were enriched for type I/II IFN response genes (Fig. 1I, fig. S3). In line with our *in vitro* data, low EZH2 activity in PCa patients was also associated with increased expression of Th-1 chemokines (*CXCL10, CXCL11*), antigen-presentation (*B2M, HLA-A*), and IFNγ regulated genes (*STAT1, IRF9*) (fig. S4, table S1).

One potential mechanism underlying enrichment of IFN gene response to EZH2 inhibition would be upregulation of interferons themselves, however supernatants from murine PCa *in vitro* models following EZH2 inhibition were tested and showed no induction of soluble IFNα or IFNγ (data not shown). Recently, epigenetic targeted therapies were shown to induce IFN gene response by de-repression of double-strand RNA (dsRNA) (*7, 9, 10*). This mechanism is known as viral mimicry and involves the re-expression of dormant transposable elements by treatment with epigenetic therapies (*7, 9, 10*). This instructs the cancer cell to adapt and respond as if infected by an exogenous virus and mount an innate immune defense via induction of dsRNA sensor machinery and IFN response genes (*20*). Indeed, treatment with EZH2 inhibitors significantly induced total intracellular levels of dsRNA in murine and human 3D PCa organoids (Fig. 1J), as well as in 2D human PCa cell lines (fig. S5) and murine PCa tissue *in vivo* (Fig. 1K). PCa patients with low EZH2 catalytic activity further demonstrated increased expression of dsRNA sensors, *RIG-I* and *MDA5*, and co-regulated innate immune receptors *TLR3* and *STING* (fig. S6). Also enriched in patients with low EZH2 activity were genes recently identified to be co-regulated by STAT1 and EZH2 that house endogenous retroviral sequences responsible for inducing an innate immune response (fig. S6) (*21*).

We next overlaid both mouse and human IFNα/γ differentially expressed genes from figure 1J and 1I to identify an overall 97-gene IFN gene signature (Fig. 2A and 2B). Importantly, cross-species analysis solidified the importance towards EZH2 regulation of TH1 chemokines (*CXCL10, CXCL11*), MHC class I pathway genes (*B2M, TAP1*), and IFN response genes (*STAT1/2, IRF9*, and *CD274*) in PCa (Fig. 2B). Collectively, there was an enrichment for biological and molecular gene ontology terms including *innate immune response* and *double-stranded RNA binding* and *Tap binding* (fig. S7), validating our previous findings (Fig. 1). Because low EZH2 catalytic activity was associated with the upregulation of these genes, we proceeded to interrogate the chromatin landscape in primary PCa patient samples. Surprisingly, these 97 genes did not display overall enrichment for H3K27me3 or DNA methylation indicating that repression of these genes was not directly regulated by EZH2 or DNA methyltransferase activity. Instead, we observe that these genes are associated with enrichment of H3K27ac and open chromatin regions as assessed by ATAC-seq (Fig. 2C, fig. S8). It was recently demonstrated that in high-grade gliomas driven by loss of H3K27me3 due to a H3K27M mutation that distinct areas of the genome become enriched for H3K27ac (*22*). These enriched H3K27ac sites appeared at repeat elements resulting in their increased expression which was further amplified following treatment with DNA methyltransferase and HDAC inhibitors, suggesting these locations are primed for rapid activation (*22*). Consistent with this, our 97-gene IFN signature was significantly up-regulated upon inhibition of EZH2 catalytic activity in PCa models (Fig. 2D).

**Fig 2.**
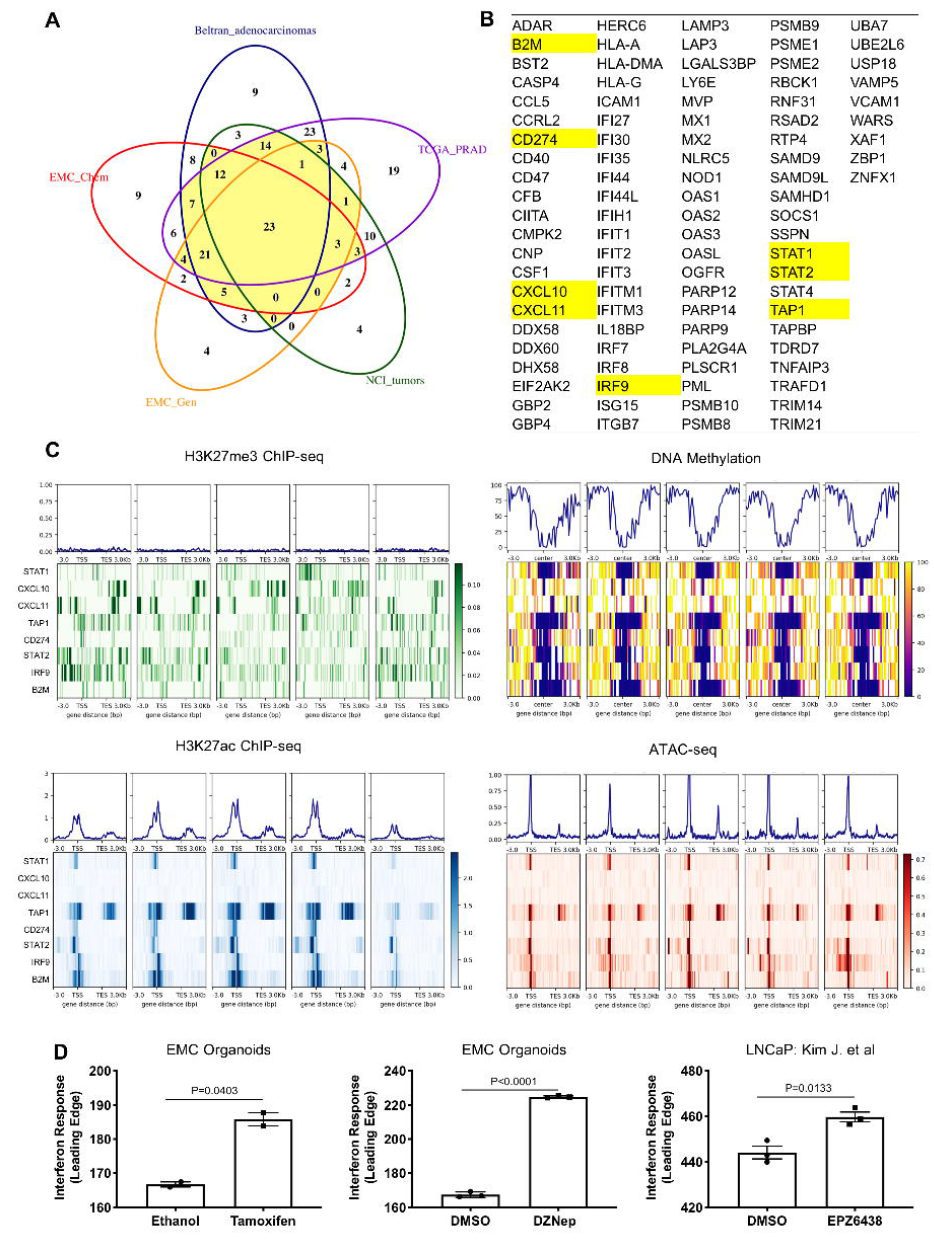
Interferon response genes are poised for activation by EZH2 inhibition. **(A)**Overlay of five independent differentially expressed IFN⍰ and IFN□ gene lists from mouse and human RNA-seq data provided a merged gene list of **(B)** 97 IFN response genes. **(C)** Selected genes (highlighted in yellow) representing IFN response (*STAT1, STAT2, IRF9*), Th1 chemokines (*CXCL10, CXCL11*), MHC class I (*B2M, TAP1*) were examined for their chromatin accessibility, DNA methylation, and indicated histone modifications in human primary PCa samples. **(D)** Mouse and human RNA-seq data was queried to demonstrate that IFN genes from (B) are upregulated in response to loss of EZH2 catalytic activity.

Within our IFN gene signature, we further analyzed the regulation of *CD274* (PD-L1) by EZH2 activity. Correlation analysis of patient PCa samples indicated that patients with lowest EZH2 activity had significant enrichment of PD-L1 gene expression (Fig. 3A). Further, human prostatectomy samples with tumor PD-L1 protein overexpression (positivity in < 5% of tumor cells by immunohistochemistry) were stained for EZH2 protein, revealing opposing EZH2/PD-L1 expression patterns in the majority of tumors (8/11 patients – 73%) were positive only for one mark (Fig. 3B-C). Treatment with two independent EZH2 inhibitors, DZNep and EPZ6438, resulted in significant mRNA and protein upregulation of PD-L1 in mouse and human PCa *in vitro* models (Fig. 3D and fig. S9). Because of the significant induction of PD-L1 expression in PCa models following treatment with EZH2 inhibitors, we sought to determine if increased PD-L1 was functionally consequential. For this, we employed the use of a mixed lymphocytic reaction assay (MLR). Two independent mouse PCa cell lines with either wild-type PD-L1 or knockout of PD-L1 were pretreated with DMSO or an EZH2 inhibitor before coincubation with murine splenocytes and evaluation of immune cytotoxicity (Fig. 3E, fig. S10). Inhibition of EZH2 activity resulted in significant loss of immune mediated cytotoxicity which was dependent on tumor cell upregulation of PD-L1 (Fig. 3F). Strikingly, immune-mediated cytotoxicity was restored in EZH2 inhibitor treated cell lines by the addition of a PD-1 antibody, and this combination effect was also dependent on tumor PD-L1 upregulation (Fig. 3F).

**Fig 3.**
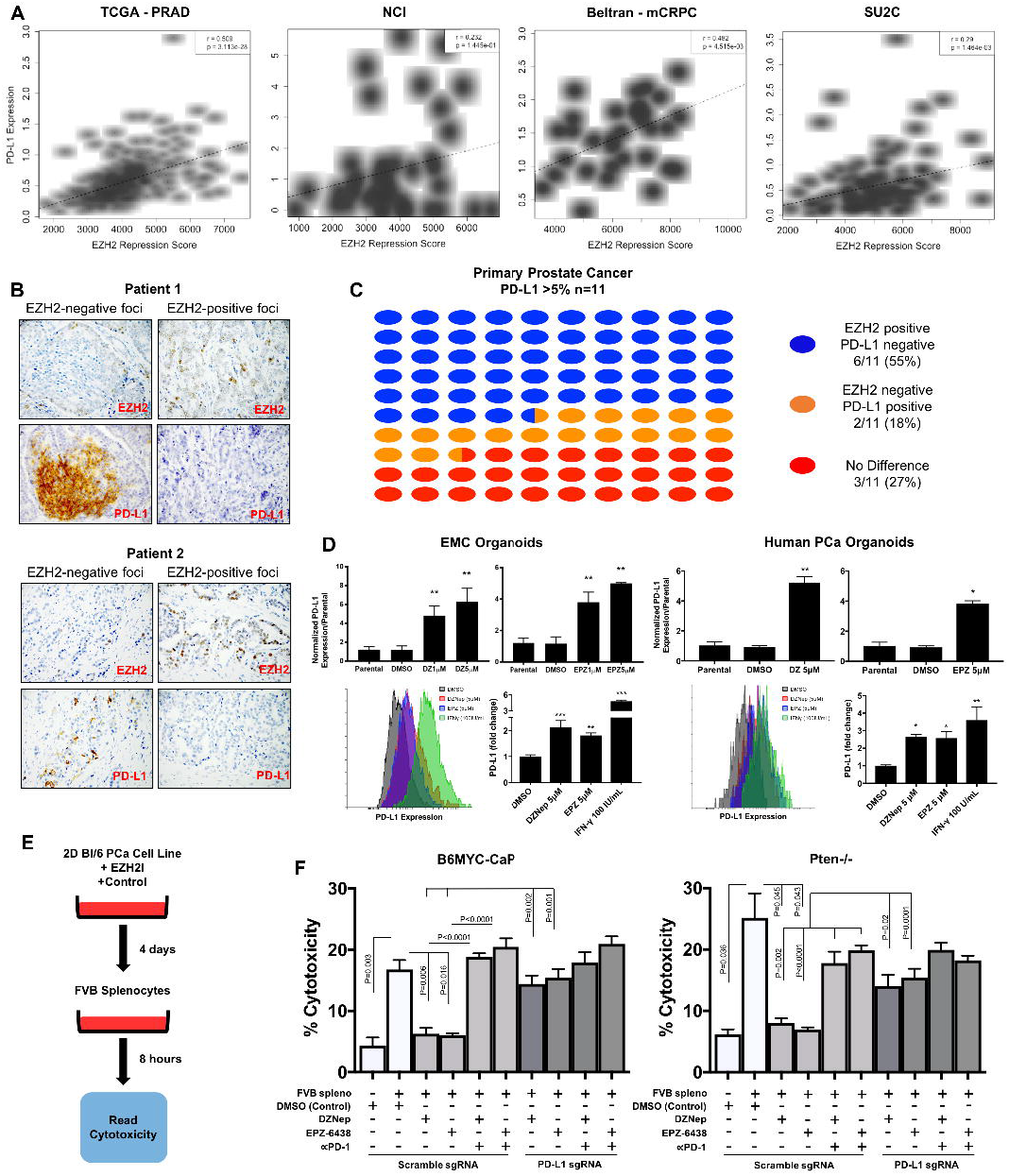
EZH2 regulates tumor PD-L1 expression and is dependent for response to PD-1 CPI. **(A)** Human PCa gene expression data was queried to demonstrate that increased PD-L1 gene up-regulation is significantly correlated with low EZH2 activity. **(B-C)** Immunohistochemical staining for EZH2 and PD-L1 using a human prostatectomy TMA revealed that a majority of patients (73%) had an inverse relationship between EZH2 and PD-L1 positive expression. **(D)** Mouse and human PCa organoids treated for 96 hours with EZH2 inhibitors significantly up-regulate PD-L1 mRNA and protein expression. **(E)** Schema of mixed lymphocytic reaction assay (MLR) protocol. **(F)** Upregulation of PD-L1 expression was functionally assessed using the MLR assay. Inhibition of cytotoxicity following EZH2 inhibition was rescued by PD-1 blockade. Inhibition of cytotoxicity following EZH2 inhibition and rescue by PD-1 blockade. This rescue is dependent on tumor PD-L1 up-regulation.

Based on the data, we proposed that EZH2 inhibition would sensitize PCa tumors *in vivo* to PD-1 CPI. Further support of this proposition was that human PCa samples with low EZH2 activity were significantly enriched for 2 gene signatures noted to predict response to checkpoint inhibition, a MImm-score (*23*) and a T-cell inflamed gene signature (*24*) (Fig. 4A). Using a Hi*MFC* PCa transgenic tissue transplant model (*25*), EZH2 inhibition (EPZ) or PD-1 CPI did not display anti-tumor activity as a monotherapy, however combination treatment produced significant therapeutic efficacy (Fig. 4B). EZH2 inhibition *in vivo* was also associated with increased tumor expression of PD-L1 (Fig. 4C, fig. S11), and reduction of PD-1 in tumor infiltrating CD8+ and not CD4+ T-cells (fig. S13). Tumor microenvironment assessment further revealed that EZH2 inhibition and combination groups showed increased accumulation of CD4+ and CD8+ T-cells (Fig. 4D, fig. S12) and M1 tumor associated macrophages (TAMs) (Fig. 4E, fig. S12). In addition, M2 TAMs were significantly decreased in EZH2 inhibited, PD-1 CPI, and combination groups (Fig. 4E, fig. S12). Other immunosuppressive infiltrating cells including myeloid derived suppressive cells (MDSCs) and regulatory T-cells (T-regs) were not significantly altered in any treatment group (fig. S13). Although both CD4+ and CD8+ T-cell trafficking was increased, only CD8+ T-cells were significantly activated in PD-1 CPI and combination groups (Fig. 4F). Using published T-cell gene signatures (*26*) we also demonstrated that human PCa with low EZH2 activity were associated with increased overall T-cell and specific CD8+ T-cell infiltration (fig. S14). In concordance with our data, it was recently shown that inhibition of polycomb repressor complex-1 in double-negative PCa models resulted in increased T-cell tumor infiltration, decreased immune suppressive cells and provided superior therapeutic benefit when combined with CPI (*27*).

**Fig 4.**
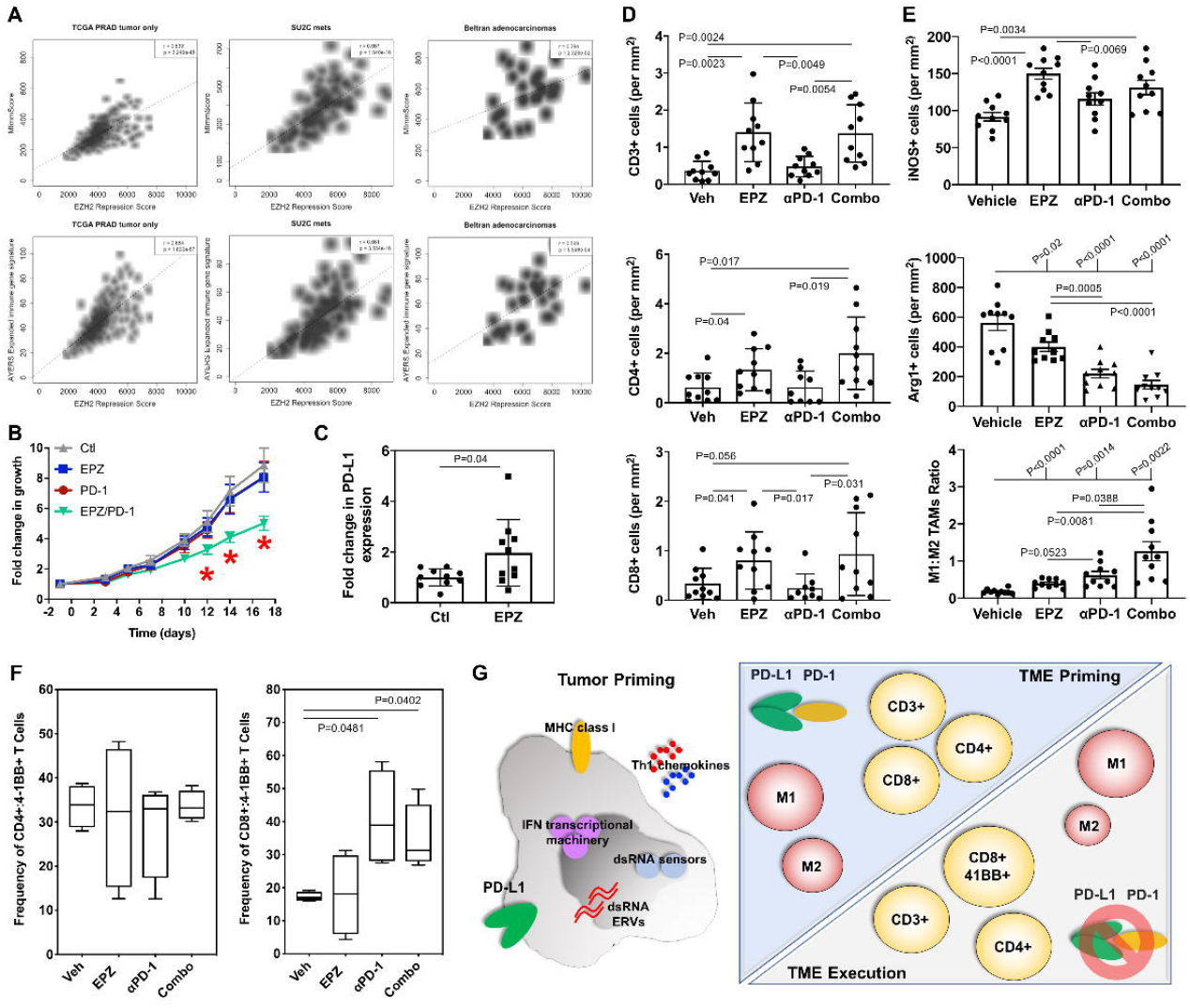
EZH2 inhibition induces PD-L1 tumor expression and sensitizes murine prostate tumors to PD-1 checkpoint inhibition. **(A)** Analysis of human RNA-seq datasets reveal immune signatures related to check-point blockade response are significantly enriched in PCa patients with low EZH2 activity. **(B)** EZH2 inhibition combines with PD-1 blockade to significantly inhibit prostate tumor progression *in vivo*. **(C)** EZH2 inhibition significantly increases PD-L1 tumor expression *in vivo* as assessed by flow cytometry. EZH2 inhibition alone or in combination with PD-1 blockade significantly increases **(D)** CD3+, CD4+, CD8+ T-cell trafficking and **(E)** increases M1 TAM with concurrent decrease in M2 TAM populations within the tumor microenvironment (TME). **(F)** PD-1 blockade alone or in combination significantly increase activated CD8+ T-cells within the TME. **(G)** Schema of overall study reveals EZH2 inhibition primes the tumor and TME by inducing viral mimicry in PCa cells (increased dsRNA) with associated up-regulation of dsRNA sensors, IFN gene transcriptional machinery, MHC class I molecules, Th1 chemokines, and tumor PD-L1 expression. This tumor response is associated with an overall increase of infiltrating CD3+, CD4+, CD8+ T-cells and M1 TAMs, and decreased M2 TAMs. The addition of PD-1 blockade to EZH2 inhibition executes the TME by further decrease of M2 TAMs and significant increase of activated CD8+ T-cells.

Identifying mechanisms driving resistance towards checkpoint immunotherapy in PCa patients remains a critical requirement. While progress has been made describing molecular events that increase response, including patients with DNA damage repair pathway defects (*28, 29*), biallelic loss of CDK12 (*30*), recycling of PD-L1 in patients lacking a common mutation in SPOP (*31*), and inhibition of IL-23 (*32*), the majority of patients treated with CPI remain unresponsive. This study provides a first look towards how epigenetic mechanisms mediated by EZH2 drive resistance towards CPI in PCa. Collectively, EZH2 inhibition in tumor cells induces dsRNA intracellular stress correlating with increased IFN response gene expression, reprogramming TAM infiltration, promoting CD8+ T-cell activation and sensitivity to PD-1 blockade *in vivo* (Fig. 4G). Together, our work provides significant insight into PCa tumor immunity, and proposes stratification of patients by EZH2 activity and generates rationale to develop combinatorial use of EZH2 inhibitors with PD-1 CPI as a novel strategy to increase PCa response to check-point immunotherapy.

## Materials and Methods

### Experimental Models

#### Mouse Models

The Institute of Animal Care and Use Committee (IACUC) at Dana-Farber Cancer Institute approved all mouse procedures. C57BL/6N and FVB mice were obtained from Charles River Labs (Strain 027 and 207, respectively). Pten^f/f^;Pb-Cre, Pb-HiMYC, Ezh2^fl/fl^, and PSA-Cre(ERT2) strains have been described previously (*17, 25, 33–36*). All models were validated by genotyping PCR analysis prior to use in subsequent studies using genomic DNA extracted from mouse ears or tails. Genotyping primers used are detailed in Table S6. The Ezh2^fl/fl^;Pb-HiMYC;PSA-Cre(ERT2)^pos^ mice (EMC) mouse strain generated in this study were mixed background consisting of FVBN and C57Bl/6.

#### 3D organoid models

All 3D organoid models were generated using previously described methodology and maintained in accordance as previously published (*37*). Clinical samples were provided for organoid generation under IRB approval (Protocol Number: 17-571 - Ellis). Human 3D organoids were generated from 2 independent patient samples - a prostatectomy and a pleural effusion sample provided by Drs. Adam Kibel and Atish Choudhury respectively (IRB Protocol Number: 01-045 – Gelb Center DFCI/HCC). Murine EMC 3D organoids were generated from the dorsolateral prostates of 8-week-old GEMMs, whereas the Pten ^-/-^ 3D organoids were generated from end-stage prostate tumors at 61 weeks-of-age. Mouse 3D organoids were validated by genotyping and recombination PCRs prior to use in subsequent studies. All primers are detailed in Table S6.

#### 2D cell lines

Pten^-/-^ and B6MYC-CaP murine cell lines have been previously described and maintained in DMEM (Gibco) supplemented with 10% fetal bovine serum (Sigma) (*25, 38*). The LNCaP cell line was obtained from ATCC and maintained in RPMI-1640 (Gibco) and supplemented with 10% fetal bovine serum (Sigma). The PD-L1 knockout models and appropriate controls [B6MYC-CaP;sgPD-L1, B6MYC-CaP;sgEmpty, Pten^-/-^;sgPD-L1, and Pten^-/-^;sgEmpty] were generated using pSPCas9(BB)-2A-Puro (PX459) V2.0 that was a gift from Feng Zhang (Addgene plasmid # 62988; http://n2t.net/addgene:62988; RRID:Addgene_62988)(*39*). PD-L1 knockout and control cell lines were generated transfecting parental cell lines with the PX459;sgPD-L1 (sgRNA sequence listed in Table S6) or empty PX459 vector using Lipofectamine 2000 DNA Transfection Reagent (11668, ThermoFisher) in accordance with manufacturer’s instructions. Cells were selected with puromycin (Pten^-/-^ 4 μg/mL; B6MYC-CaP 8 μg/mL). Following antibiotic selection, B6MYC-CaP;sgPD-L1 and Pten^-/-^;sgPD-L1 cells were treated with 20 ng/mL interferon gamma, stained for PD-L1 (558091, BD Pharminogen) and the lower expressing population was isolated by fluorescence-activated cell sorting on a BD Aria III (BD Biosciences, Dana-Farber Cancer Institute) to eliminate any residual PD-L1-positive population. PD-L1 knockout was validated by qRT-PCR following stimulation with 100 U/mL interferon gamma for 24 hours.

### Therapy Experiments

#### In Vitro Assays

For all *in vitro* therapy experiments, cells were seeded at the following concentrations: 2D cell lines were seeded at a concentration of 25,000 cells per well of a 24-well plate; 3D organoids were seeded at a concentration of 20,000 cells per 40 μL Matrigel disc (1 disc per well of a 24-well plate). In both cases, each well was treated with either 1 μM or 5μM DZNep or EPZ6438, or DMSO control, or 100U/mL interferon gamma control.

#### Mixed Lymphocytic Reaction Assay

Spleens from wildtype FVB mice were mashed through a sterile 40μm cell strainer (Corning) that had been pre-wet with sterile PBS (Gibco). Red blood cells were lysed using a commercial ACK Lysing Buffer (Gibco). The resulting splenocytes were frozen down at a concentration of 20×10^6^ cells/mL in 0.5mL aliquots. B6MYC-CaP;sgPD-L1, B6MYC-CaP;sgScramble, Pten^-/-^;sgPD-L1, and Pten^-/-^;sgScramble cell lines were cultured in standard DMEM supplemented with 10% FBS. Cultures were treated with 5μM DZNep or EPZ6438, or DMSO control for 4 days. Following EZH2 inhibitor treatment, tumor cells were washed with PBS, digested to a single cell suspension with TrypLE (Gibco), and washed with DMEM supplemented with 10% FBS. After washing by centrifugation, cells were resuspended in DMEM supplemented with 10% FBS, and re-plated into non-adherent 96-well round bottom plates. Cells were allowed to incubate with 10 μg/mL anti-mouse PD-1 antibody, or IgG control for 30 minutes at room temperature (antibodies detailed in Table S5). Following antibody incubation, splenocytes derived from FVB mice were added at a tumor cell:splenocyte ratio of 1:10. Cells were co-cultured with splenocytes for 8 hours, after which the plates were spun down and 50μL of supernatant was extracted for assessment of cytotoxicity. Cytotoxicity was measured using the CytoTox 96 Non-Radioactive Cytotoxicity Assay (Promega) according to manufactures instructions. Cytotoxicity was measured using a SpectraMax plate reader (Molecular Devices).

*In Vivo Therapy Experiment*

Pb-HiMyc derived tumor tissue (*25*) was sectioned in 2mm^2^ tumor chunks and subcutaneously implanted into syngeneic C57BL/6N mice (Charles River Laboratories). Four days following implant, mice were treated with either 250 mg/kg EPZ0011989 (Epizyme) or 0.5% CMC by oral gavage twice daily, 200μg anti-PD-1 (29F.1A12) or IgG control (2 A3) by intraperitoneal (IP) injection every 3 days started on the 5th day after initiation of EZH2i therapy, or combination (antibodies detailed in Table S5). Tumor size was measured 3 times a week by caliper measurements. Mouse weights were monitored 3 times a week. Treatment toxicities will be assessed by body weight (twice weekly), decreased food consumption, signs of dehydration, hunching, ruffled fur appearance, inactivity or non-responsive behavior. Tumor tissue from each mouse was s were further assessed by flow cytometry, histopathology and immunohistochemical procedures.

### Immunohistochemical and Immunofluorescent Staining and Quantification

#### In Vitro Samples

2D cell lines were seeded in a μ-Slide 8 Well chambered coverslip (80826, ibidi) and treated as previously described. Cells were washed with PBS (Gibco), and fixed with 4% paraformaldehyde for 15 minutes. Following another 5 minute PBS wash, cells were permeabilized by the addition of PBS containing 0.25% Triton X-100 for 15 minutes. Following 2x 5 minute washes with PBS, cells were incubated with a blocking solution [1% BSA in PBST (PBS + 0.1% Tween 20)] for 1 hour. Cells were then incubated with diluted primary antibody in blocking solution overnight at 4°C. Following 3 additional 5 minute PBS washes, cells were incubated with diluted secondary antibody in blocking solution for 1 hour at room temperature in the dark. Following 3 additional 5 minute PBS washes, coverslips were imaged using an EVOS FL Auto 2 Cell Imaging System (ThermoFisher Scientific). Antibodies used are detailed in Table S5.

#### In Vivo Samples

For immunohistochemistry, 4 μm thick sections were cut from paraffin-embedded blocks and dried onto positively charged microscope slides, deparaffinized in xylene solutions and then rehydrated in graded ethanol. Slides were boiled in 10mM sodium citrate solution (pH 6) in a microwave for 10 minutes. Immunohistochemistry staining was carried out using the ImmPRESS® HRP Anti-Mouse IgG (Peroxidase) Polymer Detection Kit (Vector Laboratories) was used according to manufacturer instructions. Tissues were incubated with primary antibodies (diluted in PBS containing 1.25% horse serum) in a humidified chamber at 4°C overnight. For protein visualization, DAB Peroxidase (HRP) Substrate Kit (Vector Laboratories) was Slides were subsequently washed in tap water, counterstained with hematoxylin and cover-slipped. For immunofluorescence, 4 μm thick sections were cut from frozen OCT blocks and allowed to dry onto positively charged slides for 30 minutes. Tissue sections were fixed in 2% paraformaldehyde (in PBS) for 20 minutes at room temperature and permeabilized in 0.1% Triton X-100 (in PBS) for 10 minutes, washed in PBST, then blocked for 1 hour at room temperature with 5% goat serum + 0.1% Tween-20 in PBS. Sections were incubated with primary antibody (diluted in PBS containing 1% goat serum) in a humidified chamber at 4°C overnight, washed in PBST and cover-slipped with VECTASHIELD® Antifade Mounting Medium with DAPI (Vector Laboratories). Antibodies used are detailed in Table S5. For analysis, 20 representative images from each tumor were taken using an EVOS FL Auto 2 Cell Imaging System. Staining intensity was scored using analysis pipelines generating in Image J software (*40*) (IHC staining) or CellProfiler software (*41*) (IF staining).

#### Clinical Samples

The human prostatectomy tissue used have been previously described (*42*), and was assessed as described by Calagua *et al*. Briefly, immunohistochemical staining was carried out on a Dako Link 48 autostainer. Sections were incubated with primary antibody for 1 hour, followed by amplification using Envision FLEX rabbit or mouse linkers, and visualization using the Envision Flex High-sensitivity visualization system (Dako). Tumor PD-L1 positivity was defined by moderate to strong membranous staining, and cytoplasmic staining was not considered. Scoring was performed semiquantitatively as follows: 0 (negative or < 1%), 1 (1%–4%), 2 (5%–24%), 3 (25%–49%), and 4 (≥ 50%). Antibodies are detailed in Table S5.

### Flow Cytometry

#### In vitro analysis

The Click-iT EdU Alexa Fluor 488 Flow Cytometry Assay kit (ThermoFisher) was used to measure DNA synthesis according to manufacturer’s instructions. Cultures were treated for 72 hours. Organoid discs were dislodged by pipetting, then digested to a single cell suspension by treatment with TrypLE (Gibco), which was in turn deactivated by resuspension in DMEM (Gibco) supplemented with 10% FBS (Sigma). Cells were washed with PBS (Gibco) by centrifugation at 500g at 4°C and fixed with 4% paraformaldehyde for 15 minutes. Following another PBS wash, cells were permeabilized by the addition of ice-cold methanol to a final concentration of 90% methanol. This suspension was incubated for 30 minutes on ice. Following 2 washes with FACS buffer (PBS supplemented with 10% FBS), cells were resuspended in primary antibody prepared in FACS buffer (antibodies detailed in Table S5). These cell suspensions were incubated overnight in the dark at 4°C or for 1 hour at room temperature. Cells were washed two additional times in FACS buffer. H3K27me3 and Edu was analyzed using an Amnis ImageStream Mark II (Luminex) and dsRNA and PD-L1 with a BD LSRFortessa (BD Biosciences).

#### In Vivo Tumor Analysis

Tumors were mechanically dissociated and filtered into single-cell suspensions in PBS on ice. Tumors were analyzed as follows. Cells were washed with PBS (Gibco) by centrifugation at 500g at 4°C and fixed with 4% paraformaldehyde for 15 minutes. Following another PBS wash, cells were permeabilized by the addition of ice-cold methanol to a final concentration of 90% methanol. This suspension was incubated for 30 minutes on ice. Following 2 washes with FACS buffer (PBS supplemented with 5% FBS), cells were resuspended in primary antibody prepared in FACS buffer. These cell suspensions were incubated overnight in the dark at 4°C or for 1 hour at room temperature. Cells were washed two additional times in FACS buffer and analyzed as on a BD LSRFortessa (BD Biosciences, Dana-Farber Cancer Institute), separated into “CD45-“ and “CD45+” events. Antibodies used are detailed in Table S5.

#### In Vivo Tumor Immune Profiling

Tumor cell suspensions were stained using two different antibody panels: lymphocytes or myeloid, using appropriate IgG and full minus one (FMO) controls, followed by analysis on an LSRII flow cytometer (BD Biosciences). Antibodies for the various immune panels are as follows: lymphocyte panel (Ghost Dye™ Red 780, anti-human CD8 (dump channel), anti-mouse CD3, anti-mouse CD4, anti-mouse CD8, anti-mouse CD45, antimouse PD-1); myeloid panel (Ghost Dye^™^ Red 780, anti-human CD8, anti-mouse CD11b, anti-mouse CD45, anti-mouse Ly6C, anti-mouse Ly6G, anti-mouse I-A/I-E). After surface staining, fixation, and permeabilization (BD Cytofix and BD Cytoperm), cells were stained for the following intracellular markers: lymphocyte panel (Foxp3, Ki67, or the appropriate IgG controls). Following staining, cells were analyzed on an LSRII flow cytometer (BD Biosciences). Cells were gated based on singlets, size/nucleation, Ghost Dye^™^ Red 780 negative events, and dump negative events (“Live events”). Cells were then separated into “CD45-“ and “CD45+” events, and immune populations were defined as follows: CD3+CD4+ T cells, CD3+CD8+ T cells, CD3+CD4+Foxp3 + Treg, Granulocytic MDSC (CD11b+ MHCII-Ly6C^lo^ Ly6G+), Monocytic MDSC (CD11b+ MHCII-Ly6G-Ly6C^hi^). Antibodies used are detailed in Table S5.

### Quantitative Real Time PCR

Quantitative PCRs were performed in accordance with MIQE guidelines (*43*). RNA was harvested using a standard TRIzol® protocol according to manufacturer’s instructions. cDNA was synthesized using the qScript cDNA SuperMix (Quantabio) according to manufacturers’ instructions. The SsoAdvanced Universal SYBR Green Supermix (BioRad) was used for PCRs using the cycling conditions recommended in the manufacturers’ instructions. Primers used are detailed in Table S6.

### Statistical Methods

Graph preparation and statistical analyses of *in vitro* and *in vivo* experiments was performed with the GraphPad Prism software. Statistical significance for assays was assessed using a Welch’s corrected un-paired t-test unless otherwise stated. Specific for *in vivo* tumor growth curves (fig. 4B), a multiple t-test was used to assess therapy response. An observation with a p-value of <0.05 was considered statistically significant.

### Sequencing Analysis

#### RNA Sequencing Data Generation

EMC organoids were seeded at a concentration of 20,000 cells per 40 μL Matrigel disc (1 disc per well of a 24-well plate) and treated with either 5μM DZNep, DMSO control, 1μM Tam, or Ethanol control for three days. RNA was harvested from samples using Trizol (ThermoFisher Scientific) according to manufacturer’s instructions. Samples were sequenced at the Molecular Biology Core Facilities at the Dana-Farber Cancer Institute as follows. RNA libraries were prepared with the TruSeq Stranded mRNA sample preparation kits (Illumina) from 500ng of purified total RNA according to the manufacturer’s protocol. The resultant RNA and ChIP dsDNA libraries were quantified by Qubit fluorometer, Agilent TapeStation 2200, and qRT-PCR using the Kapa Biosystems library quantification kit according to manufacturer’s protocols. Uniquely indexed libraries were pooled in equimolar ratios and sequenced on a single NextSeq 500 Sequencing Platform (Illumina) run with single-end 75 base pair reads. Sequencing reads were aligned to the UCSC mm9 reference genome assembly and gene counts were quantified using STAR (v2.5.1b) (*44*), and normalized read counts (RPKM) were calculated using Cufflinks (v2.2.1) (*45*).

#### Additional Datasets Used

Microarray data for LNCaP cell lines treated with EZH2 inhibitor has been previously described (*46*). Raw and normalized expression data for 550 TCGA prostate cancer samples was obtained from the National Cancer Institute Genomic Data Commons Data Portal. 102 samples were excluded based on pathological criteria provided by Dr Svitlana Tyekucheva and Massimo Loda, and the remaining 448 samples (40 normal samples and 408 tumor samples) were included in subsequent analyses. NCI data was provided by Dr. Adam Sowalsky. The Beltran collection of human prostate adenocarcinomas has been described previously (*47*) and were obtained from DbGaP (Study Accession: phs000909). Normalized counts from the Stand Up 2 Cancer dataset was obtained from cBioPortal (*48*).

#### Software/Packages Used

Differential gene expression (DE) analysis, sample-to-sample distance calculations and principal component analysis were conducted using the “DESeq2” package in R. Raw RNA-seq count data was processed to remove genes lacking expression in more than 80% of samples. Low count genes - with less than 10 total reads - were also filtered out. Following variance stabilizing transformation, a Euclidean sample distance matrix and principal component plots were generated to compare global gene expression profiles between samples. Differentially expressed gene (DEG) lists were then generated. Further interpretation of gene expression data was enabled using Gene Set Enrichment Analysis (GSEA). A ranked list was generated from the DEG output by multiplying the −log10 of the adjusted p-value by the sign of the log2FoldChange. The ranked list was then used as an input to the GSEAPreranked tool to generate enrichment scores using the Hallmark, Curated and Oncogenic Signatures gene sets in the Molecular Signatures Database. Heatmaps and unsupervised hierarchical cluster analysis, using Euclidean distance measurements, were performed using the “pheatmaps” package in R. The ‘corr.test’ and ‘smoothScatter’ functions were used for Pearson correlation analysis and to generate scatter plots. The ‘VennDiagram’ package was used to compare gene lists and generate Venn diagrams. Master regulator analysis was performed using MARINa (*49*). Protein association network generation and Gene Ontology analyses were performed using STRING v11 (*50*).

#### EZH2 Repression Score

DE analysis and GSEA was first performed using RNA-Seq data obtained from EMC 3D organoids treated with DZNep (n=3) and EMC 3D organoids treated with the DMSO vehicle (n=3). DZNep vs DMSO RNA-Seq data was used to generate a 29 gene signature, which contained the most differentially expressed genes with human homologs. Weights were again defined as the –log10 of the adjusted p-value multiplied by the sign of the log2FoldChange. The EZH2 Repression Score was generated for each tumor sample by multiplying the log-transformed count data for each of the 29 human orthologous genes by its established weighting and summing these 29 values for each sample.

#### Molecular Signatures

The HALLMARK_INTEFERON_GAMMA_RESPONSE and HALLMARK_INTERFERON_ALPHA_RESPONSE molecular genesets were obtained from the Molecular Signatures Database (MSigDB v6.2). A complete, and refined polycomb repression signature has also been previously described (*19*). Molecular signatures used to define the immune microenvironment included the Ayers *et al*. preliminary expanded immune signature (*24*), and the MImmScore (*23*).

#### Hallmark Interferon Leading Edge Gene List

DE analysis, and GSEA were performed on the 5 datasets indicated. A Hallmark Interferon Leading Edge Gene list was obtained for each dataset by taking the union of the leading edge genes identified in the GSEA reports for the HALLMARK_INTERFERON_GAMMA_RESPONSE and HALLMARK_INTERFERON_ALPHA_RESPONSE gene sets. An overall list of Hallmark Interferon leading edge genes was by genes that appeared in at least 3 out of 5 datasets.

#### ChIP Sequencing and ATAC Sequencing

Fresh-frozen radical prostatectomy specimens from patients with localized prostate cancer were obtained from the Dana-Farber Cancer Institute Gelb Center biobank under Dana-Farber Cancer Institute/Harvard Cancer Center IRB-approved protocols (Protocol Numbers: 01-045, 09-171). Hematoxylin and eosin (H & E) stained slides from each case were reviewed by a genitourinary pathologist. Areas estimated to be enriched >70% for prostate tumor tissue were isolated for analysis. ChIP-seq was performed using the protocol previously described (*51*) with antibodies to H3K27Ac (C15410196, Diagenode) and H3K27me3 (9733S, Cell Signaling Technology). Libraries were sequenced using 75 base pair reads on the Illumina platform. The ATAC-seq assay was performed at Active Motif using fresh-frozen Gelb Center RP tumor and normal epithelium specimens. The tissue was manually disassociated, isolated nuclei were quantified using a hemocytometer, and 100,000 nuclei were tagmented as previously described (*52*), with some modifications (*53*) using the enzyme and buffer provided in the Nextera Library Prep Kit (Illumina). Tagmented DNA was then purified using the MinElute PCR purification kit (Qiagen), amplified with 10 cycles of PCR, and purified using Agencourt AMPure SPRI beads (Beckman Coulter). All samples were processed through the computational pipeline developed at the DFCI Center for Functional Cancer Epigenetics (CFCE) using primarily open source programs. Sequence tags were aligned with Burrows-Wheeler Aligner (BWA) to build hg19 of the human genome, and uniquely mapped, non-redundant reads were retained (*54*). These reads were used to generate binding sites with Model-Based Analysis of ChIP-seq 2 (MACS v2.1.1.20160309), with a q-value (FDR) threshold of 0.01 (*55*). The ChIP-Seq and ATAC-seq data will be reported separately (Pomerantz *et al*., submitted). Bisulfite sequencing data from localized prostate tumors were reported previously (*56*) and processed and as previously described. deepTools (*57*) was used to create heatmaps for epigenomic data visualization.

## Acknowledgements

This study was supported by Dana-Farber Cancer Institute (DFCI) Faculty Start-Up Funds (L.E), and a Prostate Cancer Foundation Young Investigator Award (L.E.). B.M.O was supported by Emory University Faculty Start-Up funds. D.P.L. is a Lewis Katz - Young Investigator of the Prostate Cancer Foundation and is the recipient of a Scholarship for the Next Generation of Scientists from the Cancer Research Society, and is also a Research Scholar, Junior 1 of the Fonds de la recherche du Québec-Santé (FRQ-S). This research project was supported in part by the Emory University School of Medicine Flow Cytometry Core. We would like to thank Epizyme Pharmaceuticals for supplying EPZ0011989. The results shown here are in whole or part based upon data generated by the TCGA Research Network: https://www.cancer.gov/tcga.

