## Supplement Figures & Tables for "Targeting EZH2 Increases Therapeutic Efficacy of Check-Point Blockade in Models of Prostate Cancer"

#### Supplement Figures and Tables

**Figure S1**

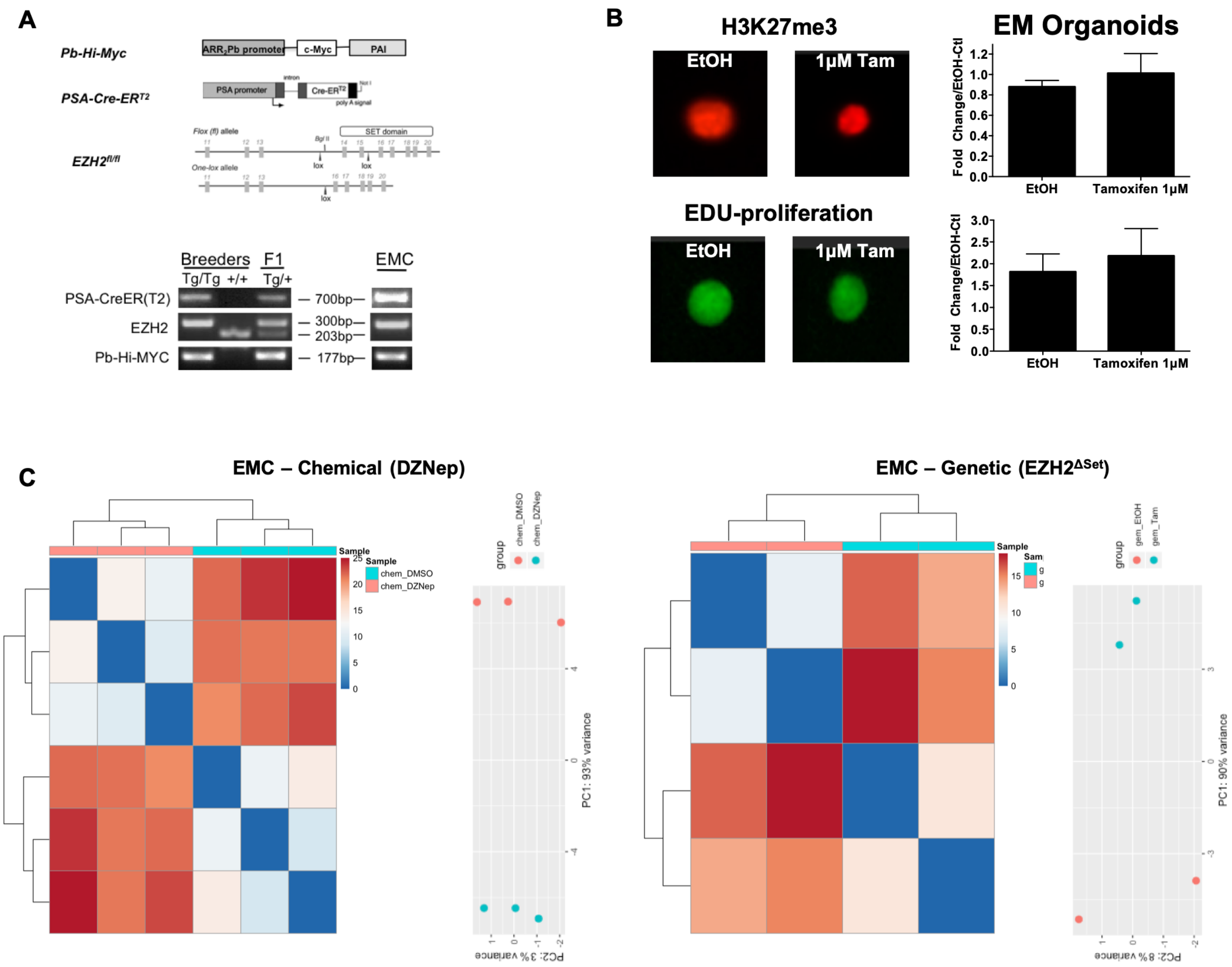


**Fig. S1. (A)** Schema and genotyping PCR example for the creation of EM and EMC genetically engineered mice. **(B)** Three-dimensional PCa organoids generated from EM mice (without PSACre$^{ERT2}$) alleles. When treated with tamoxifen, demonstrates no loss of H3K27me3 or EDU staining, indicating specificity of tamoxifen-PSACre$^{ERT2}$ mediated deletion of the Ezh2 set domain. **(C)** Principle component analysis (PCA) following chemical and genetic inhibition of Ezh2 catalytic function results in significant changes in gene expression.

#### Figure S2

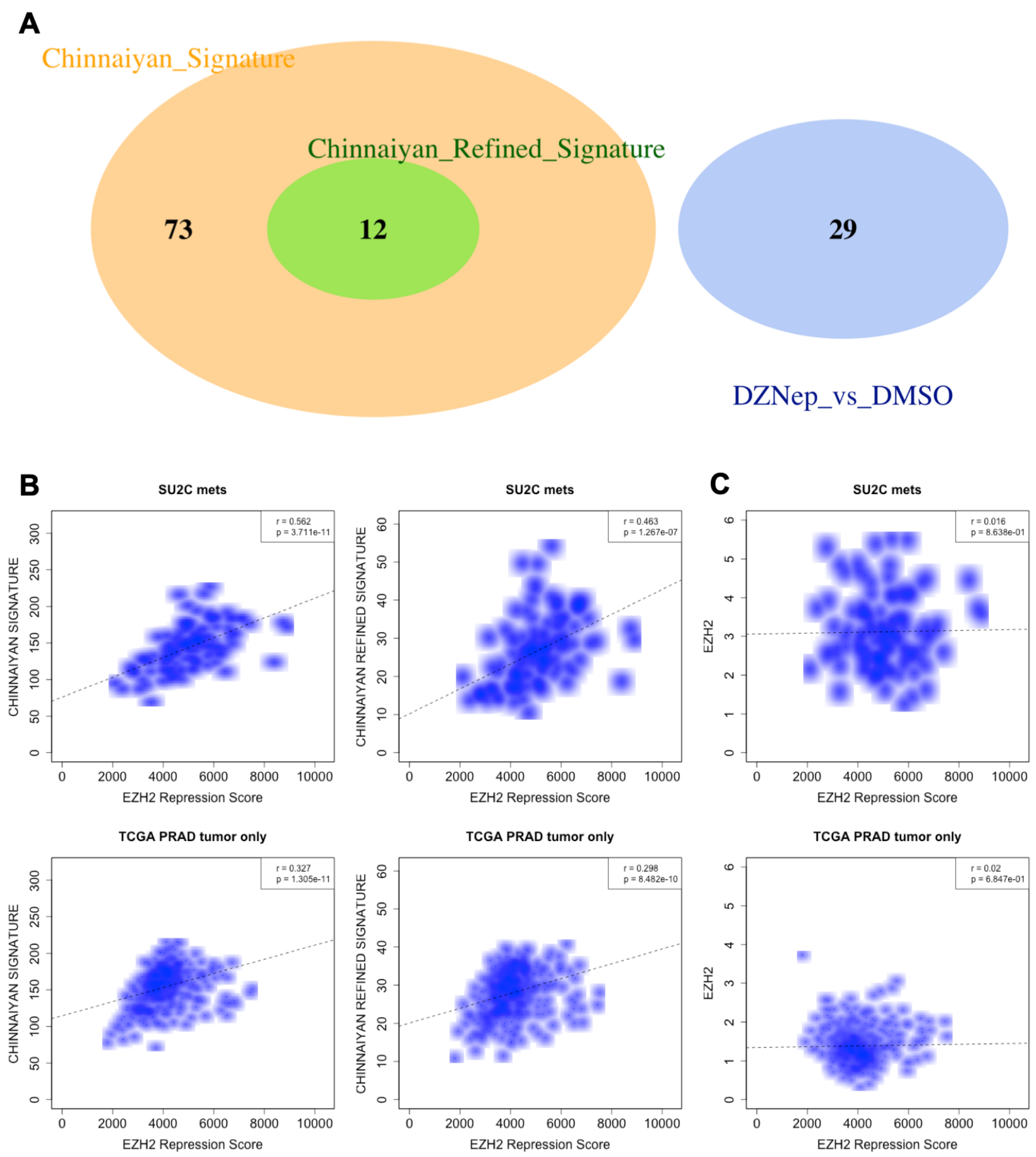


**Fig. S2. (A)** A 29-gene signature derived from Fig. 1C demonstrates complete independence from a previously published polycomb repression signature. **(B)** Our 29 gene signature demonstrates significant correlation with a previously published polycomb repression signature in 2 independent human PCa gene expression datasets. **(C)** EZH2 activity is not determined by EZH2 mRNA expression.

**Figure S3**

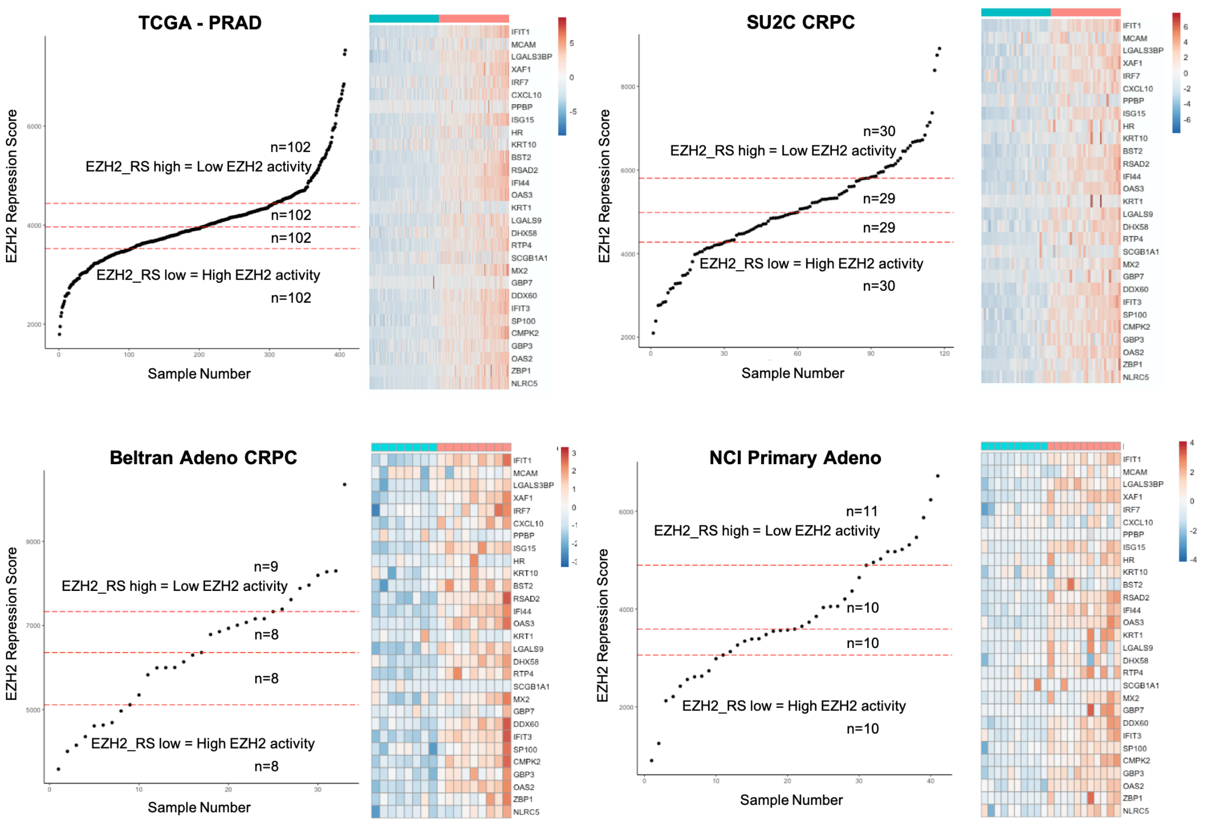


**Fig. S3.** A 29-gene signature derived from Fig 1C was used to generate signature scores for each patient within four independent human prostate cancer RNA-seq datasets. Patients were ranked highest score to lowest score and subject to quartile separation. First (blue) and fourth (red) quartiles were analyzed by supervised clustering to demonstrate expression differences within patients with most lowest EZH2 activity and most highest EZH2 activity.

**Figure S4**

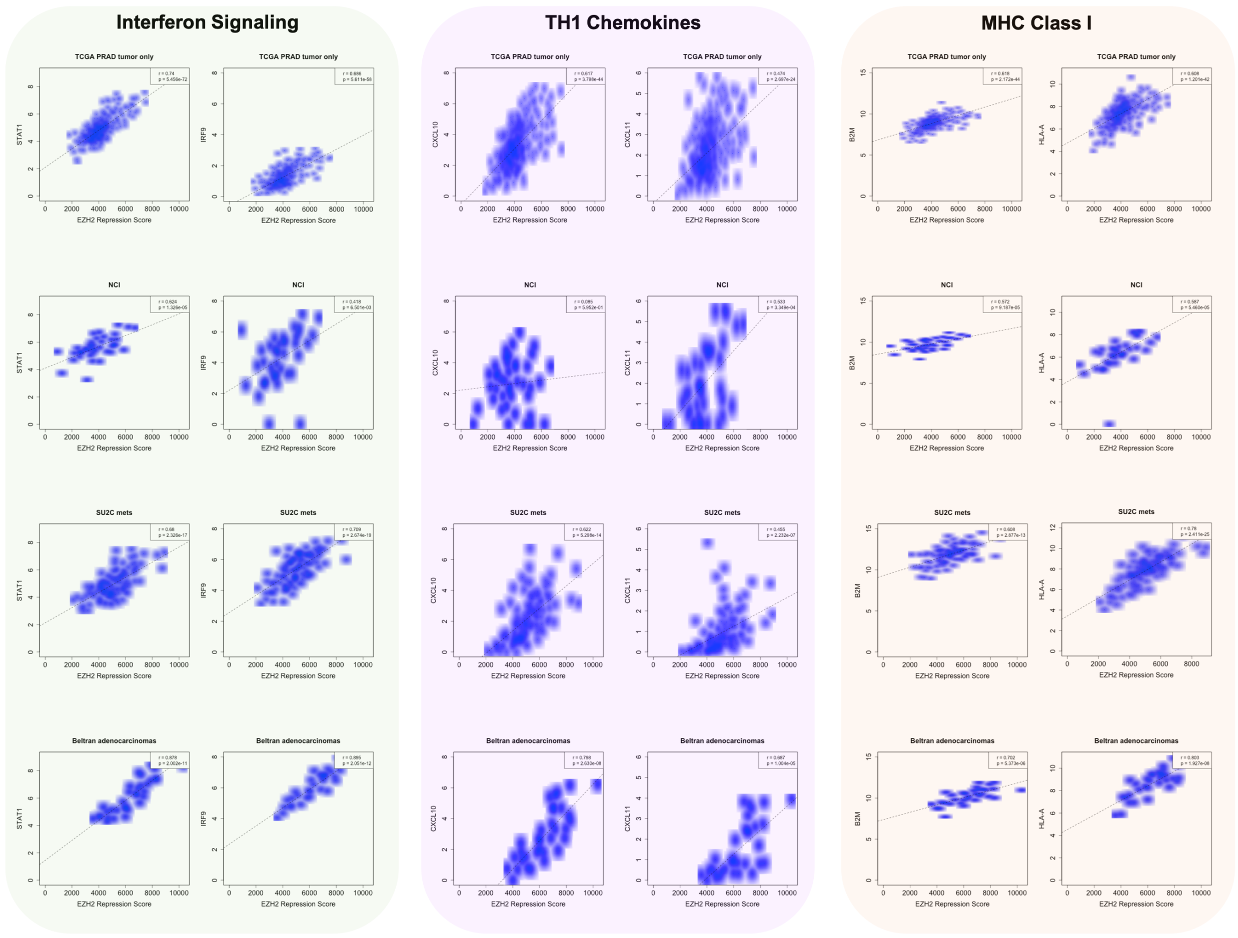


**Fig. S4.** Genes representing IFN signaling (*STAT1, IRF9*), Th1 chemokines (*CXCL10, CXCL11*), and MHC Class I molecules (*B2M, HLA-A*) were shown to be enriched in PCa patients with low EZH2 activity.

**Figure S5**

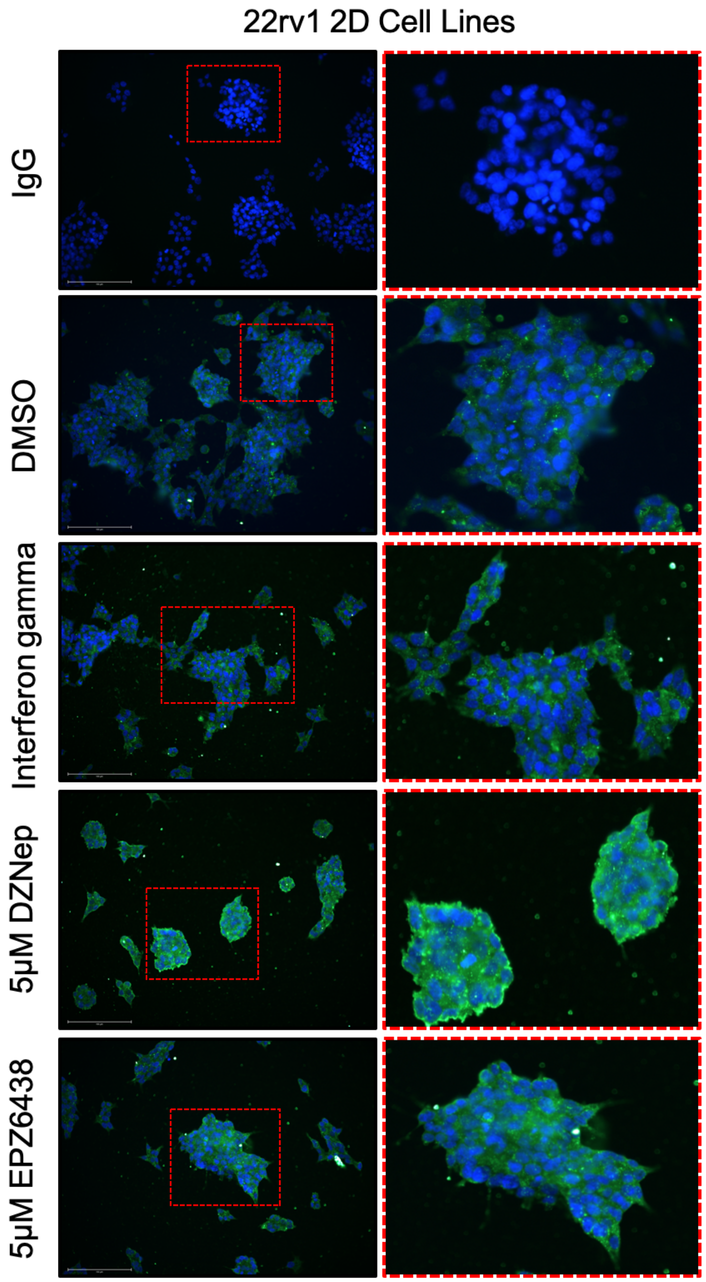


**Fig. S5.** Treatment of 22Rv1 human 2D cell lines with the demonstrated conditions for 96 hours show that EZH2 inhibition increases expression of dsRNA (green = dsRNA, blue = nuclei).

**Figure S6**

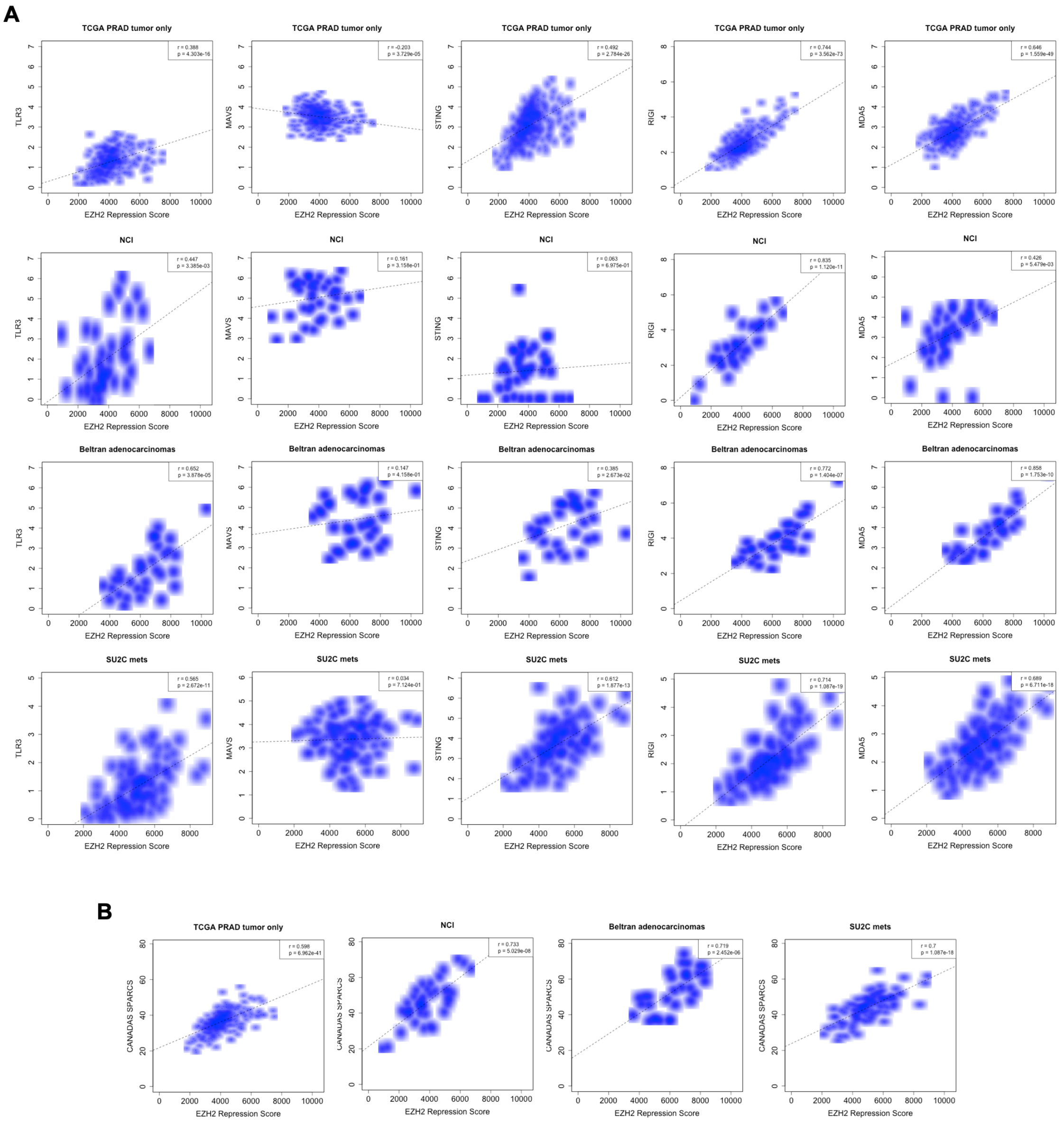


**Fig. S6.** (A) Genes representing intracellular sensors of dsRNA (*TLR3, MAVs, STING, RIG-I, MDA5*) were shown to be enriched in PCa patients with low EZH2 activity. (B) Genes from Canadas et al. (2018) described as 'SPARCs' regulated by STAT1 and EZH2 that house endogenous retroviral sequences important for inducing an innate immune response, were shown to be enriched in PCa patients with low EZH2 activity.

**Figure S7**

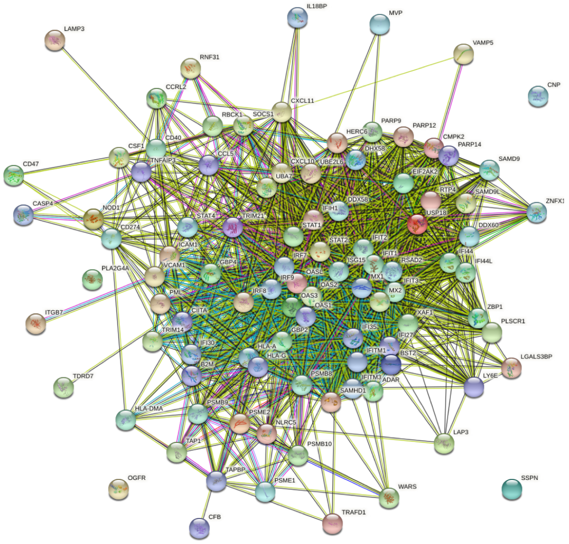


**Network Stats**

| | | | |
|---|---|---|---|
| number of nodes: | 97 | expected number of edges: | 87 |
| number of edges: | 1475 | PPI enrichment p-value: | < 1.0e-16 |
| average node degree: | 30.4 | *your network has significantly more interactions than expected (what does that mean?)* | |
| avg. local clustering coefficient: | 0.709 | | |

**Functional enrichments in your network**

**Biological Process (GO)**

| *GO-term* | *description* | *count in network* | *false discovery rate* |
|---|---|---|---|
| GO:0045087 | innate immune response | 52 of 676 | 2.84e-45 |
| GO:0006952 | defense response | 61 of 1234 | 1.28e-44 |
| GO:0002376 | immune system process | 73 of 2370 | 3.58e-43 |
| GO:0006955 | immune response | 63 of 1560 | 1.05e-41 |
| GO:0060337 | type I interferon signaling pathway | 28 of 65 | 5.63e-41 |

*(more ...)*

**Molecular Function (GO)**

| *GO-term* | *description* | *count in network* | *false discovery rate* |
|---|---|---|---|
| GO:0003725 | double-stranded RNA binding | 9 of 70 | 6.07e-08 |
| GO:0042802 | identical protein binding | 25 of 1754 | 0.00019 |
| GO:0001730 | 2'-5'-oligoadenylate synthetase activity | 3 of 3 | 0.00025 |
| GO:0042605 | peptide antigen binding | 4 of 22 | 0.00063 |
| GO:0046977 | TAP binding | 3 of 7 | 0.00083 |

*(more ...)*

**Fig. S7.** String analysis of the generated 97 type I/II IFN gene list reveals significant enrichment of biological processes including *innate immune response*, *defense response*, and *type I interferon signaling pathway*. Moreover, molecular function terms including *double-stranded RNA binding*, *peptide antigen binding* were also significantly enriched.

**Figure S8**

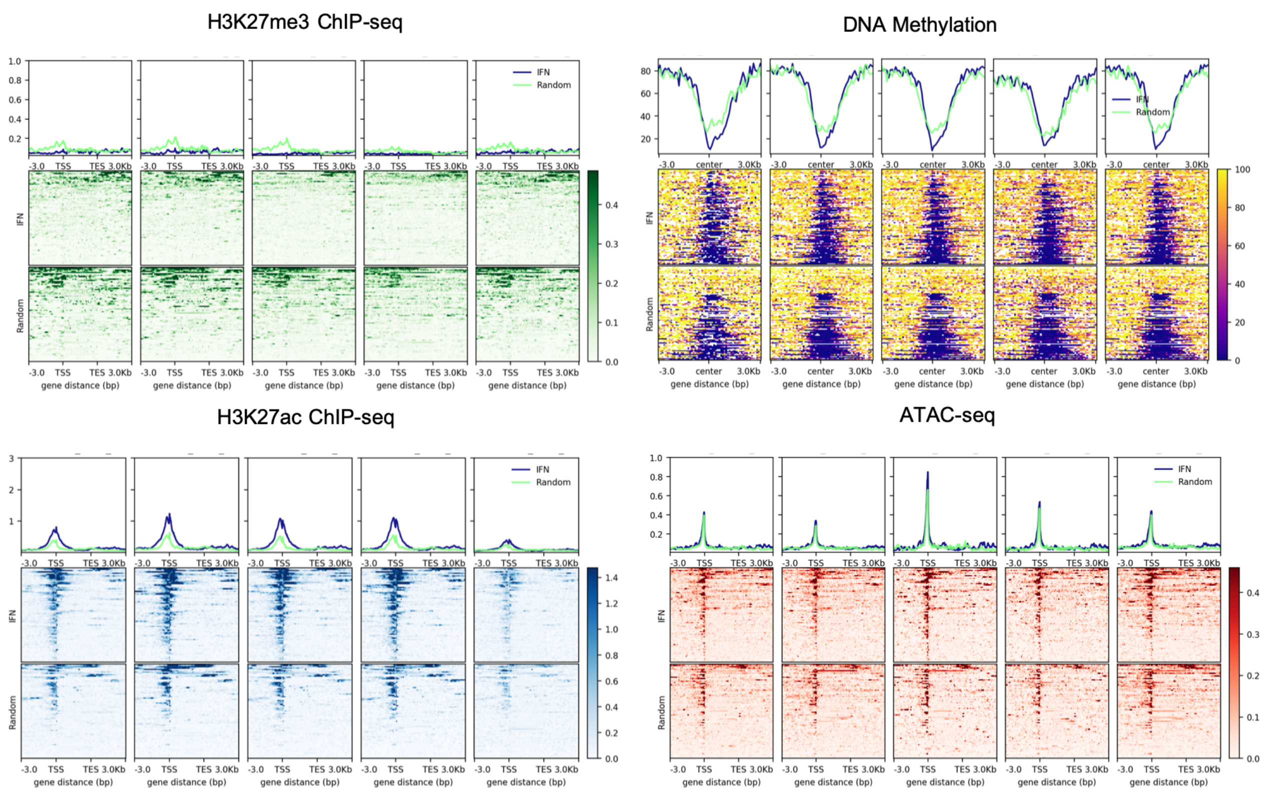


**Fig. S8.** Heatmaps of normalized tag densities for H3K27me3, H3K27ac, DNA methylation, and ATAC-sequencing within 5 independent prostatectomy patient samples.

Figure S9

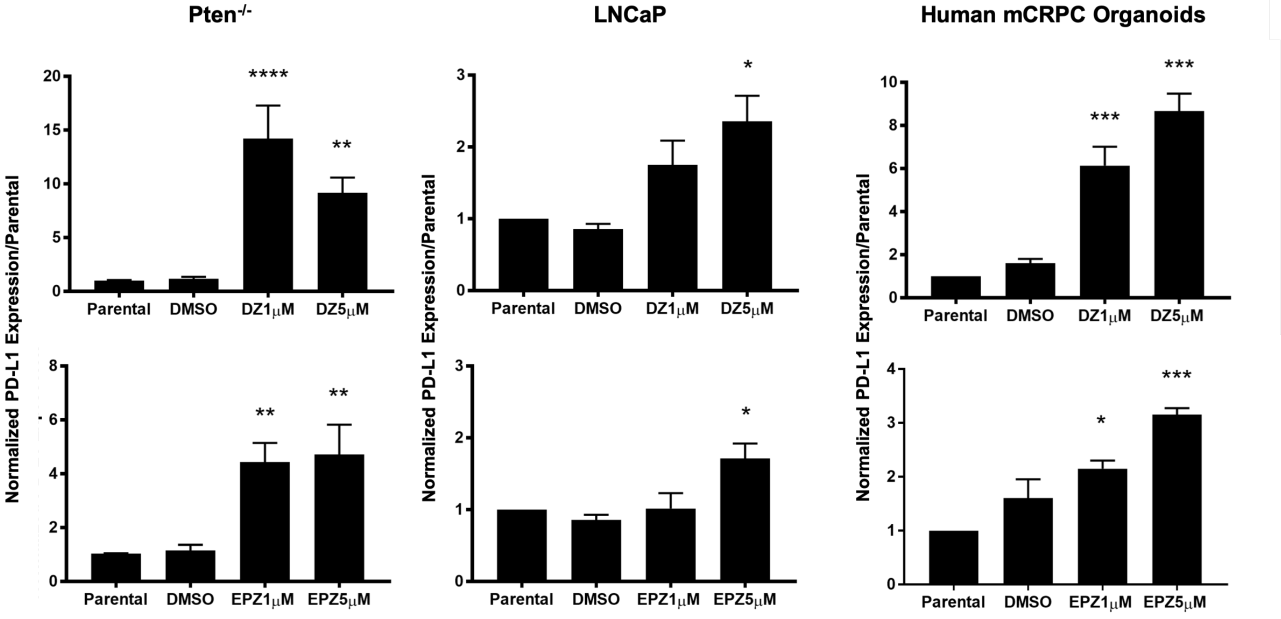


**Fig. S9.** Mouse and human prostate cancer organoids (Pten-/- and human mCRPC organoids), and human LNCaP 2D cell lines treated with indicated EZH2 inhibitors for 96 hours demonstrate upregulation of PD-L1 mRNA.

**Figure S10**

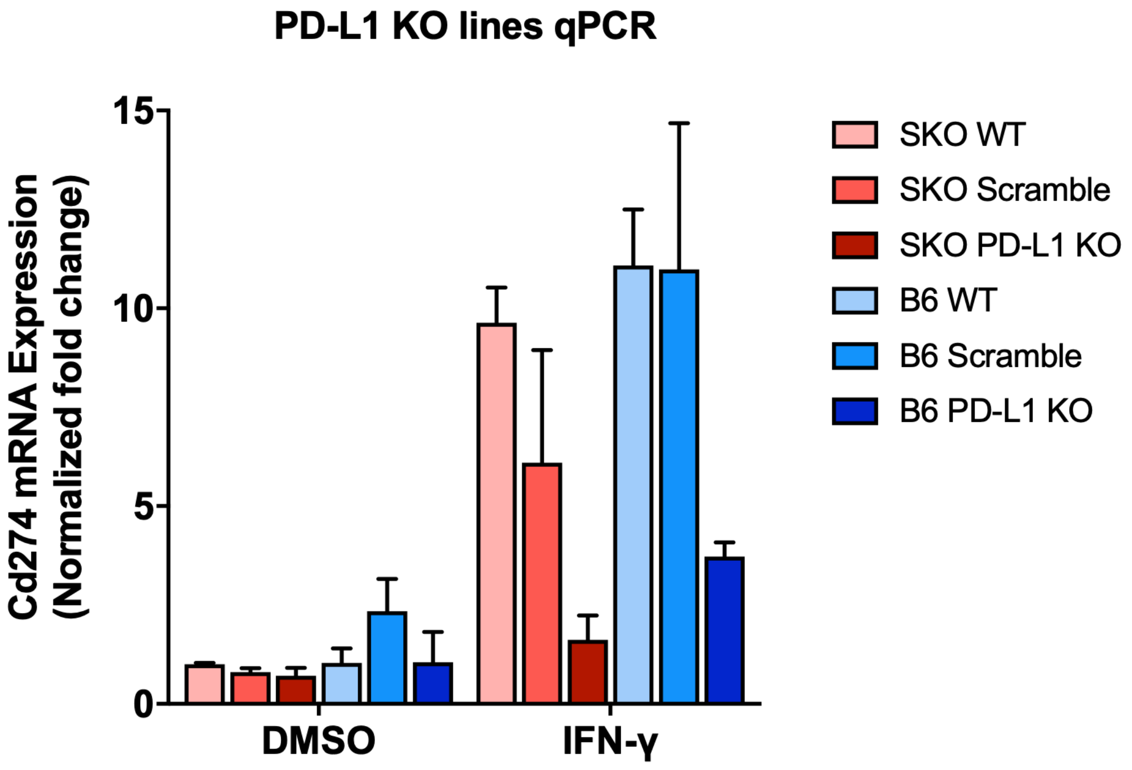


**Fig. S10.** B6MYC-CaP and Pten-/- 2D cell lines that express Cas9 were stably infected with gRNA towards *Pd-l1* (*Cd274*). Treatment with IFNɣ validates the inhibition of *Pd-l1* expression in KO cell lines.

#### Figure S11

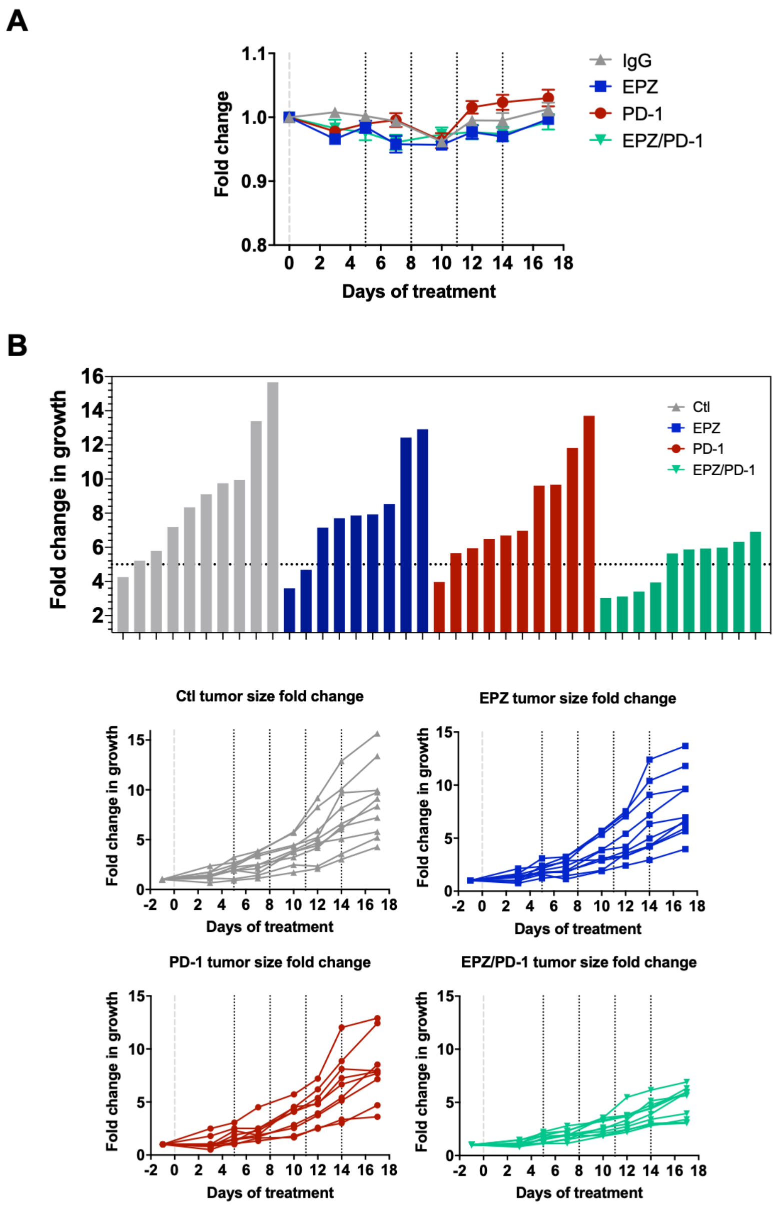


**Fig. S11.** (A) Normalized weights of mice indicate that no significant weight loss (ie: toxicity) was observed following therapy with indicated treatment cohorts. (B) Alternate tumor measurements of individual tumors by waterfall or spider plots validate significant anti-tumor activity of EZH2 inhibition combined with PD-1 blockade.

**Figure S12**

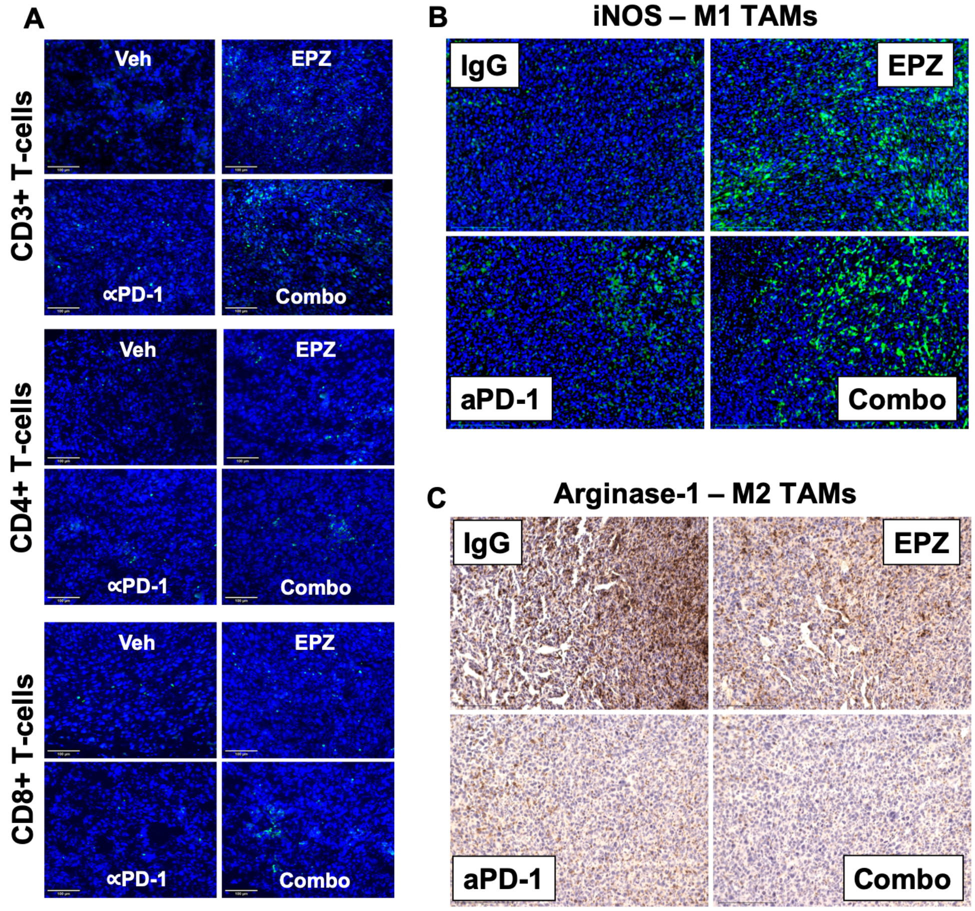


**Fig. S12.** Representative *in vivo* tumor analysis of immune infiltrate staining of CD3+, CD4+, CD8+ T-cells and M1 and M2 TAMs in B6-HiMYC PCa tumors. Scale bar = 100µm (T-cells), and 200µm (TAMs).

**Figure S13**

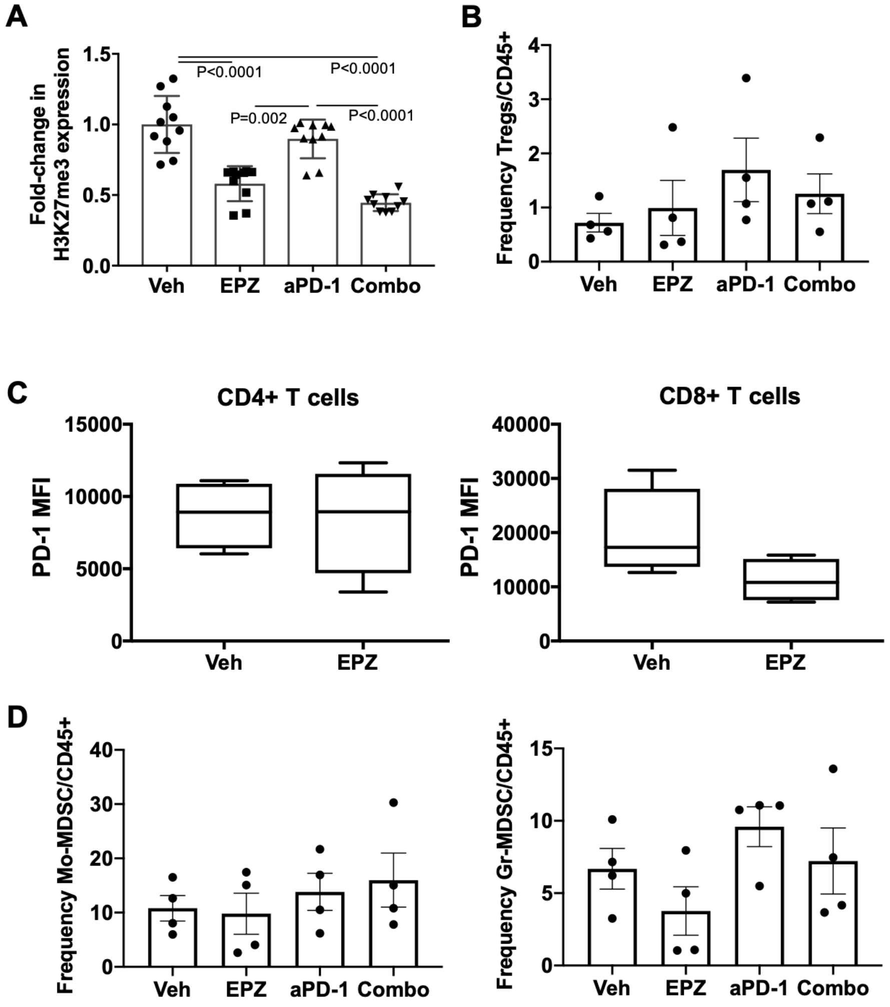


**Fig. S13.** (A) Representative *in vivo* tumor analysis indicate that EZH2 inhibition and combination significantly reduce tumor H3K27me3 expression. (B) Frequency of Foxp3+ T-reg cells was determined by flow cytometry. No significant change was observed following treatment. (C) PD-1 protein expression on CD4+ and CD8+ T-cells was analyzed by flow cytometry. Only CD8+ T-cells were observed to express lower PD-1 protein following EZH2 inhibition. (D) Frequency of Mo-MDSC and Gr-MDSC cells was determined by flow cytometry. No significant change was observed following treatment.

### Figure S14

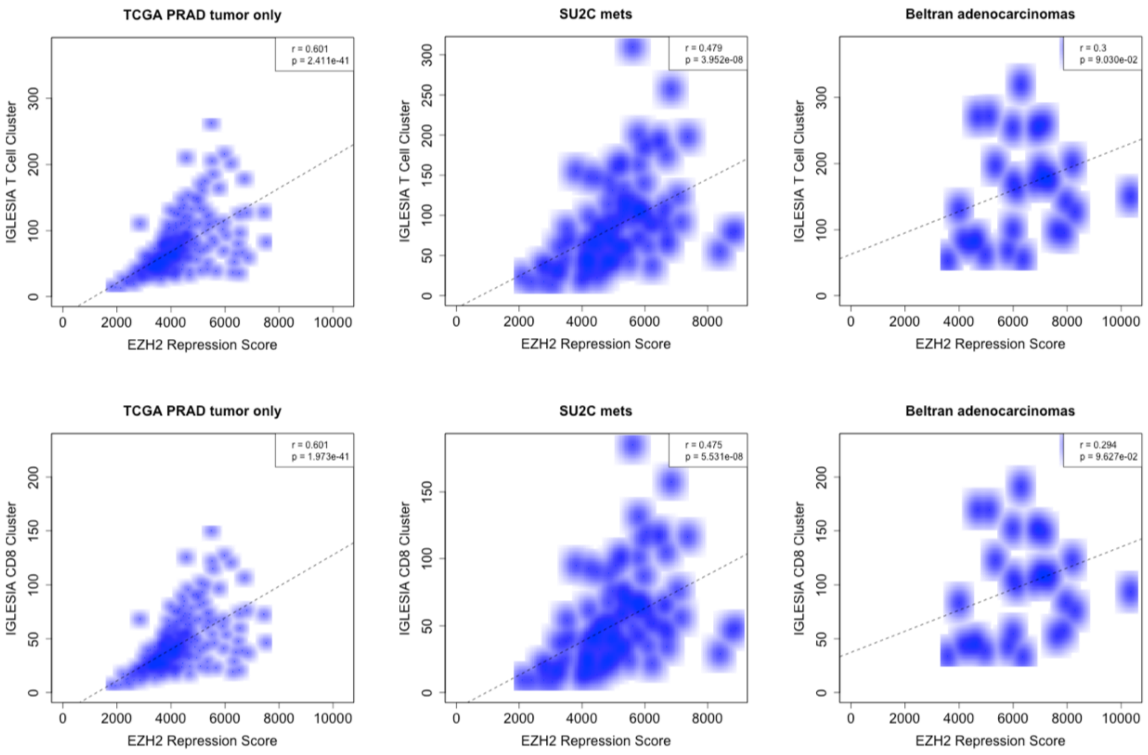


**Fig. S14.** Correlation analysis for T-cell and CD8+ T-cell verse EZH2 activity score in PCa patient samples from TCGA PRAD, mCRPC SU2C Dream Team, and mCRPC-Ad from Beltran et al. Gene signature used for T-cell identification was from Iglesia et al.

### Table S1. Significantly differentially expressed genes (EZH2 Signature High vs Signature Low, p<0.05)

| EMC organoids - DZNep vs DMSO | | | | EMC organoids - Tam vs Eth | | TCGA - PRAD (Top 3992 genes) | | | | Trento/Cornell/Broad | NCI | |
|---|---|---|---|---|---|---|---|---|---|---|---|---|
| IFIT1 | SLC14A1 | GCNT4 | LTBR | KRT13 | CLUH | IFI44 | WISP1 | EIF2A | FBXO17 | OAS3 | OAS3 | HRC |
| OASL2 | MPHOSPH8 | BACH2 | 9030624G23RIK | SLURP1 | WBSCR16 | IFIT3 | IGLV3-9 | TNXA | SCHIP1 | IFI44L | XAF1 | CTLA4 |
| MCAM | HTR1D | DUSP14 | ETV4 | DSG3 | KIRREL | IFIT1 | RP11-374F3.5 | STX2 | LINC00607 | MX1 | ISG15 | GPR65 |
| STAT1 | AI462493 | AK010878 | RAB23 | PPBP | MCM3 | MX1 | TLR8 | IER3 | ACTR3B | IFIT1 | LGALS3 | ACP5 |
| GM14446 | GSTP1 | STON2 | ABHD2 | KRT16 | SRP19 | IFI44L | GCNT4 | ICA1 | RRAD | IFI44 | UTS2 | HNRNPAB |
| KRT13 | RPS20 | NDUFAF5 | IBA57 | IL33 | PAPD5 | RSAD2 | IGLV3-12 | HPCA | MLC1 | OAS2 | GPR115 | CPAMD8 |
| LGALS3BP | PCMTD1 | TRAPPC12 | SIRT6 | CLCA3 | BET1 | OAS1 | AP001610.5 | DIO3OS | CLSTN1 | IFI6 | PTGS2 | OBSCN |
| XAF1 | CYB5R3 | STRBP | NAA20 | AQP3 | TRIM13 | EPSTI1 | MR1 | CHST6 | RP11-452C13.1 | IFI27 | ETV4 | CYB5A |
| IRF7 | ENPP2 | DZANK1 | RTKN2 | UBE2D3 | CDC42EP5 | ISG15 | IGFBP4 | DUOX2 | RUNX1 | CHRNB2 | SLPI | SOCS1 |
| SLFN2 | FUBP3 | GM4841 | KRAS | PPP1CB | BC030336 | OASL | TMEM45A | IFNWP19 | CFAP221 | STAT1 | MBOAT1 | TMEFF2 |
| IRGM1 | PLXNA3 | WIPF3 | ZBTB12 | KRT4 | TXNDC17 | IFITM1 | TRAV5 | PPM1M | RP1-153P14.7 | RSAD2 | CMPK2 | NLRP1 |
| CXCL10 | GRPEL2 | PANX1 | SEMA6D | SPTSSB | MANBA | IFIT2 | RBP1 | TAS1R3 | HNRNPL | SPTSSB | IFI44L | TBX5 |
| PPBP | YIF1A | TCERG1 | SOAT1 | RPTN | SLC25A27 | IFI6 | CXCL17 | HOXD9 | CTBP1-AS2 | IFIH1 | IFIT1B | GLIS3 |
| SAMD9L | CEBPD | PPP1R26 | ABCA1 | KRT6A | YBX1 | XAF1 | PRR5L | CACNA2D3 | ENPEP | CST1 | NSA2 | LY96 |
| IRF9 | IER3 | HERC3 | POLR2G | LGALS3BP | PTEN | CTD-2521M24.9 | FAM131B | LRFN5 | C5orf66-AS1 | RUNDC3A | SLFN12L | MAPKBP1 |
| ISG15 | ATP2C2 | ZFP207 | SASS6 | LYPLA1 | HCCS | BST2 | IL27RA | RP1-28O10.1 | MCF2L-AS1 | OAS1 | IFI6 | VAV3 |
| UGT2B34 | RPS24 | 0610007P14RIK | CP | SERPINB11 | FAM76B | SAMD9 | SLC29A1 | RAPGEFL1 | CABYR | EPHA7 | LOC441869 | YWHAE |
| HR | RABGGTB | ZBTB9 | INSR | GSTO1 | VAMP5 | ZBP1 | AC246787.4 | CLEC4E | NKX3-1 | CMPK2 | DDX56 | MAPK7 |
| KRT10 | MED16 | CAPN2 | STRADA | AGR2 | PKD2L1 | SAMD9L | PLCL2 | AIRE | LINC00908 | DDX60 | MYO1F | C11ORF66 |
| GSDMC2 | PHYHIP | CBR1 | PTPN6 | CYP2F2 | DTX3 | CMPK2 | AC246787.3 | SLC1A7 | RP11-761I4.4 | XKR7 | ABHD2 | CCNB2 |
| IRGM2 | TRIM26 | HEATR3 | DBP | ANXA8 | GALNT6 | BATF2 | RP11-730A19.9 | ASGR2 | CEACAM5 | UNC79 | EGOT | DPT |
| NUPR1 | EAF1 | PHF11C | DIDO1 | LTBP2 | NSA2 | PSMB9 | TNFAIP3 | ADCY4 | ASTL | ISG15 | MX1 | FBLN2 |
| PARP14 | ITGB4 | SDR42E1 | BLOC1S4 | KLF4 | TRIM24 | PSMB8-AS1 | VDR | CDC42EP2 | TGFB1I1 | TF | OAS1 | GPR180 |
| OAS1A | FAM195B | CEP63 | IGIP | COL4A1 | PUS1 | OAS3 | SYNE3 | MYBL1 | ABHD10 | ZNF710 | CPXCR1 | RIBC1 |
| CLU | MYO19 | MRPL33 | DDIT3 | SH3BGRL | GLRX3 | NLRC5 | CARD6 | CGB7 | SLC2A9 | APOL1 | ECM1 | NKX3-2 |
| BST2 | CLASRP | TAF12 | LARS | SELT | EIF1A | HLA-F | IGHV3-63 | WASF3 | MARVELD1 | SCG3 | STK32C | C14ORF79 |
| RSAD2 | BCAS3 | CRNN | HTATIP2 | DMKN | R3HDM1 | MX2 | RUNX2 | NUMBL | AC000089.3 | CXCL11 | HSD17B14 | ZNF433 |
| CXCL5 | ISOC2B | HINT2 | SPRR2H | PTBP3 | UPK1B | CARD11 | IGKV1OR2-6 | SOX15 | SGMS1 | BSN | PDGFD | ATXN7L2 |
| STAT2 | PLEC | S100A16 | SLC4A7 | SLC7A11 | KRT20 | CD247 | LINC00943 | NRP2 | EID3 | HERC6 | ZSWIM3 | GPATCH1 |
| IFI44 | LACTB | GINS1 | UTP23 | GPNMB | ASAH1 | GPSM3 | THY1 | DTD2 | RP11-75L1.1 | CXCL10 | ASAH1 | OSCAR |
| CXCL3 | YIF1B | BBS12 | DCAF12 | TRP53INP1 | YIPF4 | SP140 | PSME1 | ACKR1 | PTH1R | RTP4 | C5ORF56 | SULT2B1 |
| UPK3A | CYB5 | RNF44 | NOXO1 | CRNN | ARPC2 | CD6 | SLC16A4 | GPR115 | NT5DC1 | JPH3 | IL20RB | SVIL |
| KRT14 | MPPED2 | KLF3 | 2210408F21RIK | ACER3 | MFSD7C | APOL1 | CTD-2547L24.3 | TRIM17 | DGKZ | CP | LOC284440 | ZNF793 |
| APOL9A | MAN2C1 | A4GALT | PLCG1 | FUT9 | ARFRP1 | CASP1 | GAS6 | DNAJC5 | CNOT6 | HDAC9 | RASL12 | ABCG5 |
| GSDMC3 | BUB1 | GGN | TAB2 | MARCKS | ABO | PCED1B-AS1 | C1orf106 | FXYD6 | ICE2 | SPRN | SLC26A4 | DUSP16 |
| TAP1 | NDUFAF3 | ITM2B | TTC13 | VMA21 | RMND5A | LGALS9 | CR1 | DSG3 | RPS18 | DNAH11 | STARD13 | HIST1H2BO |
| OAS3 | ATRN | 5330426P16RIK | MOB3B | TMED7 | TMEM154 | CD48 | ELMO1 | CX3CR1 | AC005330.2 | VTN | MYO7B | PRH2 |
| IFIH1 | CST3 | MYBBP1A | 43164 | LCN2 | GM1821 | TRAC | CTD-2506J14.1 | SLC39A2 | RP11-431M7.3 | BST2 | OGN | SLC9A8 |
| IGTP | LIX1L | MUT | PTPRF | TMED5 | GCNT2 | CORO1A | SERPINF1 | ROBO2 | AF001548.5 | IFIT3 | KCNJ13 | SULT1A3 |
| KRT1 | PSPC1 | D15ERTD621E | RNMTL1 | COPS2 | S100A16 | TMC8 | STRA6 | THRAP3 | RPS8 | PEX5L | BNIP3L | TMSB10 |
| PIGR | PLXNA2 | ARHGEF39 | CARKD | ATF3 | CLSTN1 | RTP4 | GPBAR1 | RP11-342D11.3 | PITHD1 | C1orf168 | IFITM1 | ZNF28 |
| USP18 | PHC2 | SAMD1 | FAM13A | MBNL1 | YIPF6 | RAC2 | IGLV4-60 | RASL11B | RP11-10K16.1 | EDEM3 | TFIP11 | ITGA1 |
| OAS1G | YIPF1 | CLIP2 | PEX11G | LIN7C | NFIX | PSTPIP1 | ZEB2-AS1 | A2M | PXDC1 | TPTE | CHST14 | PRICKLE3 |
| KRT16 | CUL7 | 3110062M04RIK | AP3S1 | NEURL3 | FAM111A | CD37 | LINC00926 | RP11-483E17.1 | KCNJ8 | GP1BB | KANK2 | LOC151009 |
| LGALS9 | TTLL3 | CCDC141 | ATP6V1G2 | PTGES3 | TINAGL1 | HLA-E | LAG3 | RP11-223C24.1 | CLCN1 | ALOX12B | FCHSD2 | FAM129A |
| B2M | GEMIN4 | MAPK3 | VPS33A | MZT1 | TWF1 | CD3E | FCGR2A | RPL7AP11 | SPIN4 | CTNNBL1 | DLEC1 | ADAM11 |
| SPON2 | NANOS1 | BAG6 | CAPN13 | SGPP1 | CAPZA2 | GBP1 | CR1L | FRZB | FAM102A | TDO2 | PGBD4 | CHI3L1 |
| TRIM12C | LCMT2 | NAIP6 | MSANTD2 | DYNLT3 | TMEM65 | CD52 | CD68 | FAM155B | TMEM139 | ASGR1 | CMAH | SPATS2 |
| DHX58 | GNG10 | 5031414D18RIK | RNASEH2C | PCNP | SLMAP | PTPN7 | RP11-426C22.5 | ASPN | C12orf60 | ACTL6B | PON1 | LOC645431 |
| RTP4 | COL9A3 | TGFBR3 | GALNT6 | IDI1 | GSPT1 | SNX20 | CMKLR1 | PPP2R2D | ITGA3 | IFI35 | FRMD3 | NPM1 |
| SCGB1A1 | AXIN2 | PLEKHG3 | CALD1 | 43715 | PDCD4 | TRBC2 | SULF2 | FTOP1 | DUOXA1 | UNC5A | FLT3 | SH3RF3 |
| MX2 | SPNS2 | GNPTAB | HIST1H2AI | ENPP2 | ZFP518A | ARHGAP9 | GABRP | RP11-401P9.5 | MBP | KDELC2 | ADORA2B | LACTB |
| TGM2 | WDR62 | KDELC1 | HIC2 | KRT14 | GSTA4 | TAP2 | KIFC3 | GSN | RP3-523K23.2 | LHX4 | CERCAM | C14ORF93 |
| VNN1 | KCNF1 | PTCD1 | FPGT | TMEM176B | CDK18 | LTB | PLEKHH2 | MFSD2A | CSTF1 | LDHB | BAALC | CLDN5 |
| IFIT2 | CRYL1 | SYNGR2 | GM11110 | LRMP | ARHGEF1 | TRBV28 | LOXL1 | PLCB4 | TRIM6 | PRTFDC1 | MRPL17 | HLA-F |
| PTP4A3 | EZH2 | E2F4 | MALSU1 | LY6G6C | NAB1 | MAP4K1 | GPR82 | C15orf53 | TAGLN | AC129492.6 | GIMAP6 | LOC392196 |
| PTMS | PSMD11 | NOP16 | USP28 | TROVE2 | ZNRF1 | TRAF3IP3 | RP11-876N24.1 | LINC00519 | SPTLC3 | CCL19 | FCGRT | LYG1 |
| PRR9 | S100A10 | SYT8 | HDAC3 | FAM3B | CALM2 | BIN2 | NRROS | RSPH3 | OLFML3 | LGALS3BP | TRPC4 | CCNK |
| CBS | RDH11 | SMTNL2 | EDF1 | CAV2 | MSL2 | TRIM22 | NAALADL1 | CARD9 | WDR86 | SFN | LOC643387 | EFHD1 |
| GBP7 | TWF2 | CLDN1 | COX7B | FAR1 | GRASP | DDX60 | CNR1 | LINC01197 | RP11-666A8.9 | SCRN1 | AMICA1 | CCL21 |
| SERPINB6B | PEX6 | MFAP1A | TMPRSS11BNL | PSCA | ERH | STAT1 | DZIP1L | TMEM200B | CSNK1E | GRIK3 | ADAMTS10 | TSSK4 |
| TIMP3 | FAM210B | PDGFD | NDC80 | PRKAR2B | ARPP19 | UBE2L6 | IGHV3-52 | LGR6 | RERG | TBC1D14 | WNT2 | HTRA4 |
| DDX60 | STYK1 | TMEM173 | TMEM259 | TGM1 | S100A11 | SASH3 | RP11-169D4.2 | TTYH2 | MYO15A | DHX58 | ZNF473 | ASPG |
| SERPINB9 | MFSD1 | ABCA2 | ASXL1 | GABRP | CACUL1 | IL12RB1 | AC011899.9 | VNN3 | IQCJ-SCHIP1 | LAMP3 | DENND2C | KIAA1539 |
| DTX3L | UNC13D | IDE | PROSC | LIF | CYB561 | HLA-DOB | PDE1B | STX1B | MAGEA8 | BCAN | FBXL15 | MURC |
| APOL9B | TGFB1 | RRP8 | LYN | NR2F2 | SREBF1 | JAK3 | IGHV3-71 | RP4-555D20.2 | CTD-2192J16.15 | CPLX2 | SP1 | TNKS2 |
| TNNI2 | PLEKHH3 | ZFP512 | SLC9A2 | TMEM123 | TMPO | CD2 | IGHV3-73 | CALHM1 | NBR1 | PGPEP1 | EPHB2 | ACO1 |
| OGFR | RPLP2 | EIF3H | HIRIP3 | DR1 | NES | CD3D | SIGLEC14 | RNASE1 | CTD-2270N23.1 | AP3B2 | GNB3 | ATP2A1 |
| IL20RB | MRI1 | HOXD11 | SIPA1L3 | CYP4F39 | FIRRE | PARP15 | TMEM176B | WDR49 | RPL39L | DISP2 | KIAA0406 | EBF1 |
| PHF11D | SLITRK6 | AP1M1 | SCAMP2 | ACSM3 | NFKBIA | POU2AF1 | MAP3K8 | ZYX | CNTN1 | MATK | CHPT1 | HSPA14 |
| GSS | 2810006K23RIK | KRT75 | PDZRN4 | SLFN2 | FMNL3 | CCR7 | HMHA1 | CACHD1 | CCL14 | XAF1 | GLI1 | POLN |
| GPT | ZFP292 | TRABD | UHRF1 | NKX3-1 | PSMB10 | CD53 | POU2F2 | CTA-384D8.36 | PKD2L1 | CRABP2 | ANKDD1A | BTK |
| 1600020E01RIK | MIA | SLC27A3 | SPPL2A | CSF3 | FAM83G | PARVG | IGKJ5 | ANKRD33B | SEC23IP | CBX6 | ARHGAP8 | PIK3C2G |
| CXCL1 | SOX7 | ZNRD1AS | NKTR | COL7A1 | ADD3 | ARHGAP15 | CX3CL1 | AJUBA | UGDH-AS1 | KCNH6 | KCNA3 | KCNJ3 |
| IFIT3 | MTA1 | RGS9BP | ATHL1 | PANK3 | IFIT1 | TBX21 | RARRES2 | RP11-359E10.1 | BCAT1 | PTGDS | FGD3 | PARP1 |
| PARP12 | METTL16 | PIK3R1 | E130307A14RIK | CNOT6L | ABCE1 | GBP1P1 | YPEL4 | DPEP3 | PSD4 | TMEM132B | VASH1 | FBXL12 |
| SP100 | TRA2A | SLC38A10 | CD320 | CD164 | HID1 | SIT1 | RP11-288L9.1 | CAV2 | TRIM38 | CYB5B | MTMR10 | LRRFIP2 |
| DNASE2A | PROM2 | HSD3B7 | VAMP2 | NR4A1 | ORMDL1 | WAS | AC019117.2 | ANTXR1 | ACSS3 | BATF2 | LDHAL6B | ZSWIM6 |
| H2-T23 | DUS4L | RPL13 | B4GALT5 | ABRACL | UBXN7 | HCLS1 | C1orf54 | CLEC1A | MATN3 | C1orf21 | MFI2 | AVPI1 |
| UBE2L6 | EPHB3 | ARSB | LAP3 | CITED2 | CDCP1 | SELPLG | ADAM12 | IGFBP7-AS1 | TM6SF2 | UCK2 | IFI44 | PSMB8 |
| LAMB1 | TJP3 | ART3 | PPP1R18 | STK3 | MPP5 | ETV7 | RCN3 | LINC00877 | DUOX1 | PTPRT | PLA1A | OR1L3 |
| NKX3-1 | MNDA | TRIM34B | RHBDF1 | YTHDF3 | KATNBL1 | LSP1 | PBX4 | MIR205 | RP11-286H15.1 | AACS | GLOD5 | SHMT2 |
| H2-K1 | P2RX4 | GPRC5D | ATG4C | SPON2 | HNRNPR | CD74 | IGKV1D-39 | LPCAT4 | RP11-20O24.4 | CCNG2 | TRMT5 | TRAF4 |
| CES2E | THADA | ID4 | MRPL40 | S100A9 | UBXN2A | ITGAL | TMEM119 | HOMER2 | TMEM51-AS1 | USP18 | GAB3 | LOC728640 |
| MX1 | OGFRL1 | SEC24D | SFT2D2 | OASL2 | MGAT3 | SUSD3 | TRDV1 | EIF4BP6 | RBMX | PEBP4 | RP9 | KIRREL2 |
| CHID1 | UPK3BL | SSBP4 | TMEM220 | CDKN1B | MRPL32 | RASAL3 | IGFBP7 | EEF1A1 | RP3-467K16.7 | RAP1GAP2 | TOR1AIP2 | NDUFB11 |
| CMPK2 | ULK2 | SLC6A9 | GM4890 | RPE | CHPT1 | SLAMF1 | IGHV1OR15-2 | GPRASP1 | LAT | KCNJ6 | SIL1 | GIMAP4 |
| ATP6V1C2 | ZFP503 | TTC9 | VRK3 | 2700089E24RIK | PDLIM5 | CD79A | IGKV3OR2-268 | RP11-640L9.1 | RPL23 | CLDN10 | PCOLCE | KRR1 |
| KRT17 | CDCA8 | BIRC2 | RBFOX2 | MAL2 | ARSK | CD27 | PCOLCE | DRP2 | RASL11A | BEGAIN | DAAM2 | ANP32B |
| FRMD4A | CAR2 | TEX264 | GGA1 | IRF7 | GRAMD1A | IFI27 | LINC01358 | TSSC2 | RP11-566K19.6 | NF2 | DIDO1 | TBC1D9B |
| LGALS7 | TMEM160 | TEFM | CMIP | BST2 | CAPRIN1 | IL16 | PTGDR | MRPL42 | RND2 | EXTL3 | IL29 | TNFSF9 |
| GBP3 | POLR1E | AU018091 | SAP30BP | SMEK2 | GNB2L1 | CCDC109B | GPX8 | GIPC2 | RP11-541M12.6 | SPATS2L | LIN7A | C14ORF33 |
| B4GALNT2 | LTBP2 | TTC22 | SLC2A8 | GJA1 | MMS19 | APBB1IP | MACC1 | APOC1 | PRMT5 | STAT2 | LOC145783 | C22ORF24 |
| TMEM176A | RING1 | PHAX | CDC14B | IRF9 | 2410002F23RIK | CD96 | BTBD19 | MEFV | RAET1E | C8orf44 | PGS1 | CDCA7 |
| KCTD14 | LMNB1 | ZCCHC2 | KARS | GDF15 | SNAP29 | INPP5D | LINC00487 | IL23A | PLEKHH1 | LRTM2 | PLGLA | PTN |
| DDX58 | UBE2R2 | PDIA5 | MAU2 | XAF1 | SHB | ZAP70 | GPR25 | C16orf54 | AC109642.1 | HSD17B6 | RRN3 | HADH |
| GAS6 | SAV1 | GPI1 | PRPF39 | TRAM1 | SMARCA5 | S1PR4 | GPR31 | HHEX | PRRC2A | CRP | SARDH | PNCK |
| PARP9 | SMPDL3B | ZFP511 | LSM11 | JUN | ZMPSTE24 | CD5 | C9orf47 | TRBV6-4 | MAT1A | PARP12 | BTF3L4 | SERPINB13 |

#### Table S1. Significantly differentially expressed genes (EZH2 Signature High vs Signature Low, p<0.05) continued.

| | | | | | | | | | | | | |
|---|---|---|---|---|---|---|---|---|---|---|---|---|
| ANKK1 | DNAJA3 | ATP6V1B2 | GM20300 | GT(ROSA)26SOR | HIVEP2 | LY9 | RP11-472N13.3 | HOXB-AS1 | PINK1-AS | RPN2 | CA12 | XPO7 |
| IFI35 | POLR2A | YIPF3 | PIWIL2 | ETV4 | VPS4B | PYHIN1 | AC007386.4 | GRIK5 | CTC-429P9.3 | C1orf111 | CERK | N6AMT2 |
| ECI2 | PGM3 | POLH | ZFP628 | DDX3Y | CSDE1 | GPR132 | DTHD1 | ERN2 | COL9A1 | DLGAP3 | ENDOD1 | NFATC2IP |
| ACTA2 | AMOTL1 | RPS8 | SLC35A1 | OAS1A | TAGAP1 | CARD16 | KCTD17 | RSPO1 | MEG9 | CXCL13 | LOC728723 | ABHD14B |
| PARP10 | D8ERTD738E | PRPF31 | AQP3 | NAA30 | PI15 | AIM2 | C11orf45 | TMEM192 | STARD5 | FSCN2 | SEMA6C | CFC1B |
| RNASE4 | ABCD1 | SIRT2 | HEXB | SERPINB5 | CMIP | NCF1 | RP11-589N15.2 | NRM | BRI3BP | EPPK1 | LGALS9C | ODF3L2 |
| TAP2 | ATRAID | ARRB1 | PLSCR3 | QK | UBE2J1 | STAT4 | LYZ | RNF213 | PXN | LRFN1 | PLK5P | TREX1 |
| ITGA3 | SERPINB8 | APOC1 | SETD2 | TMOD3 | CTSL | RUFY4 | CD1B | KIAA1549 | SLC19A1 | QPRT | ANO10 | PPOX |
| KANK4 | OST4 | ZBED5 | HOXB5 | PCSK6 | PDLIM2 | CXCR3 | MMP12 | KIF13B | GOLPH3L | PDIA2 | HERC3 | ZNF286A |
| IFI27L2A | NFYA | CELF4 | GM12888 | ARFIP1 | SLC25A36 | IL2RG | IGHV1-12 | PTGES3P1 | ANGPTL6 | A2ML1 | ZNF135 | PPP1R3E |
| KIT | TTYH2 | NPHP3 | PLK1 | GM17066 | NPTN | CD72 | FLI1 | SYNE1 | DENND2D | TGM4 | ATHL1 | TDP2 |
| CTSA | TNFSF4 | PRELID2 | KPRP | KRT15 | IFI203 | MS4A1 | LGALS1 | STARD9 | QDPR | KCNB1 | ADORA1 | TTC22 |
| ENTPD8 | PFKL | SPTB | TM2D3 | RAP1A | SNX13 | CST7 | ZFP36L1 | CECR5 | NDE1 | C5orf15 | CDKAL1 | WNT3 |
| ACE | FHL1 | FGF13 | GPR125 | ZBTB7C | ATP9B | IL10RA | CTC-231O11.1 | LRIG1 | TRAV18 | TEDDM1 | KIF26A | CYP3A7 |
| PLXND1 | YDJC | MSX1 | CCBL2 | UBE2G1 | CAT | CCL5 | CCL13 | C10orf82 | GFAP | SFRP1 | MATN2 | FLJ10661 |
| KRT79 | HSP90AB1 | SLC1A1 | RBM25 | GMFB | UGDH | LINC00861 | AGPAT4 | GJB3 | RPS24P8 | HOXB9 | PAPL | TMEM63B |
| OAS2 | DTWD1 | TRPM5 | CDC42EP5 | DSTN | ATF2 | CLEC2B | TRBJ2-2P | BTN1A1 | ARHGEF28 | OLFML3 | SETD3 | ZSWIM1 |
| RSRP1 | DPYSL3 | CDA | UNC13A | ECHDC1 | FAU | GBP5 | LINC01503 | TRBJ2-2 | HOGA1 | IFIT5 | TGFB1I1 | FLJ36777 |
| VNN3 | TRAP1 | PTCD2 | DAPK3 | TMX1 | TCHH | OPTN | GLT8D2 | RAB34 | CCNA1 | DCN | TNF | ECT2 |
| ELK3 | ENC1 | CEP68 | TRMT1 | SERP1 | PI4K2A | APOBEC3G | BFSP2 | CCR10 | TRAV30 | DNAH10OS | F13A1 | EGFLAM |
| CLCA3 | FASTKD1 | BC029722 | TRUB2 | CALD1 | AGPS | CYTIP | PLEKHA2 | RP11-1024P17.1 | FAM73A | ZBTB7B | ARHGAP24 | IRF9 |
| LTBP4 | STAB1 | EIF3K | CPE | SBSN | NGFRAP1 | GZMA | GGT5 | CCDC88A | ZNF275 | UGT2B10 | SMS | HIST1H3I |
| ADH7 | COL16A1 | AHI1 | 1700021F05RIK | NRARP | A530054K11RIK | OAS2 | DAZL | ACY3 | CLVS2 | MMP1 | TCN1 | NEXN |
| CTGF | CH25H | E430018J23RIK | COMT | SYNCRIP | ATP6V1B2 | ITK | MIR8071-1 | ST20-AS1 | IL1R2 | PDGFC | C16ORF86 | ZNF727 |
| CLCA2 | SLC11A2 | ZFP703 | USP35 | KPNA4 | SDHC | FGD2 | SUCNR1 | RP11-554A11.4 | IRF2 | MTMR7 | THAP8 | JUNB |
| ID3 | GPR132 | CHL1 | EVPL | RABGAP1L | RNH1 | IL15 | MMP19 | SLC39A6 | AKNAD1 | RTN2 | KCNE3 | YEATS4 |
| GPNMB | ZWILCH | 1700025G04RIK | SLC2A4 | ZFP91 | SLC4A7 | IL18 | GABRE | ARRB2 | PTGES | FAHD2B | FMNL3 | SOX21 |
| TNC | PDE4DIP | GTF2H2 | RFX5 | PAPLN | CORO2A | SLAMF7 | IL15RA | CYS1 | ADM5 | SMOC1 | ICAM1 | ZSCAN5A |
| ATF3 | CELF2 | OIT1 | 4930556M19RIK | EPHA7 | DESI2 | ARHGAP25 | RP11-615I2.2 | MSR1 | SYVN1 | LTBP1 | PPP4R1 | ORAI2 |
| MOXD1 | E2F7 | PHF11B | WNT9A | RAB1 | ZFP266 | SLAMF6 | TOMM20P2 | PRSS12 | GFRA3 | APC2 | STAMBP | SEC1 |
| I830012O16RIK | FMO2 | NEDD4 | 4930503L19RIK | POF1B | TTLL7 | UBASH3A | GSTP1 | HLA-G | RASL12 | DDX58 | ZNF133 | MID1IP1 |
| EEF2 | AIRN | PLEKHG4 | ZMAT2 | KCNN4 | MAP4K5 | FCHO1 | VIM | RGL1 | TTC16 | INPP1 | ZNF434 | PROM1 |
| 2810474O19RIK | TCEA2 | FOXK2 | VDR | 2210407C18RIK | SLC52A3 | SCML4 | IGHV3-47 | RP11-330M19.1 | HTR3A | PPARGC1A | CCDC89 | LTB4R |
| TRIM21 | PIK3AP1 | D730048I06RIK | LTV1 | NGFR | WFDC2 | HLA-DPB1 | RRN3P2 | MMP7 | RPL10 | MFSD2A | RMRP | RFPL3 |
| SERPINE2 | CHTF8 | CREB3L4 | CDKN1C | PGAP1 | SH3BGRL2 | FGD3 | PAPLN | POLR3H | RP11-91K9.1 | FPR3 | FGD5 | NRAP |
| SLC40A1 | HIBCH | LRRC40 | GPS1 | NABP1 | ARIH1 | UBA7 | PRKCB | CLU | SYNDIG1L | 43165 | SRD5A2 | ATAD2B |
| IGFBP5 | SOX21 | COMMD9 | CAPN1 | SRSF1 | LANCL1 | PARP14 | SFXN3 | ZNF24 | RP11-802E16.3 | AMPH | SNHG6 | KIAA0564 |
| ZBP1 | ECSIT | AAMDC | ANGEL1 | PCMTD1 | INHA | LCP2 | MOB3B | FCAMR | BSND | GJB2 | PAK1IP1 | ELOVL7 |
| OASL1 | HOXA9 | AKNAD1 | SIL1 | FCHO2 | MED14 | ARHGAP30 | MOXD1 | MKX | MRPS30 | NOS1AP | LAYN | EMILIN3 |
| CCPG1 | RPL31 | DHRS7B | VPS25 | ZBTB16 | FAM46A | AC006129.2 | EVC2 | AC053503.6 | HOXB-AS2 | RAB3D | PML | GRAPL |
| GAS5 | CTBP2 | MEX3D | SLC35A2 | CHML | SRSF2 | IL18BP | CTC-378H22.1 | ECM1 | IGHV3-22 | ZNF395 | HSPA1A | LMBRD1 |
| HERC6 | PRR11 | ZSCAN22 | TSPAN9 | KITL | 5730508B09RIK | USP30-AS1 | KLRC1 | TOMM20 | ATP13A2 | CEACAM6 | PPP3CA | ASPRV1 |
| GLB1 | SRSF9 | FAM172A | SPCS3 | TMEM33 | RABAC1 | FAM26F | S100A16 | KRT4 | ITGB1BP2 | DOCK5 | AMAC1 | DAAM1 |
| AK1 | SUZ12 | CTNS | BTRC | CD14 | SH3YL1 | CD4 | IL22RA2 | CHST13 | STK39 | B3GALNT1 | C15ORF26 | ACOT4 |
| NIPAL2 | 2810417H13RIK | RAB9 | DPM3 | CCNI | VAV3 | RP11-288L9.4 | MSLN | RGS16 | SHF | HERC5 | HLA-DRA | C1ORF87 |
| PHLDB1 | IAH1 | CEP72 | RPS15 | CMPK1 | SAMM50 | IL21R | IGHV1-3 | PEAK1 | NPR2 | SMTNL1 | KCND1 | KIAA1755 |
| NLRC5 | ARL3 | WDR1 | EHBP1L1 | NAA50 | EHF | CD19 | HSPA6 | CD207 | AQP1 | MAGEA3 | KCNQ4 | UMODL1 |
| PABPC1 | IFRD1 | BCR | NEO1 | TMEM30A | CLCN3 | KIAA0125 | EIF4BP7 | NLRP7 | RP11-20D14.6 | OSR1 | PDZD4 | ZNF395 |
| MMP7 | DNAJC27 | XRCC6 | EARS2 | ELL2 | PLEKHA3 | LTA | MUC16 | IGKV2-29 | RP11-342I1.2 | CNKSR3 | TRIM43 | CNKSR3 |
| TTLL7 | CXCL16 | CHCHD4 | DHRS13 | ABI1 | GM11974 | ZC3H12D | RP11-23P13.6 | ADAMTS9-AS2 | DEAF1 | ERICH1 | GLUD1 | CADM2 |
| CES2B | IFNE | APCDD1 | FEM1C | WIF1 | IGIP | CD7 | IFNG | ZNF488 | RABGAP1L | SLC4A4 | CABYR | HS3ST5 |
| SULT1C2 | HDGF | CKB | CCDC149 | CHP1 | E2F5 | FCRL3 | SSTR3 | MAP3K14 | SDC3 | NRSN1 | KIAA1274 | TTN |
| RGS9 | EIF4B | NHLRC2 | PRRC1 | RAP1B | ATP11B | IFITM3 | TRAV26-2 | FOXI2 | RP11-498C9.15 | GPC5 | RERGL | APOF |
| DAP | CHKA | ITPR1 | MBLAC2 | FAM199X | DHX58 | LAPTM5 | RP11-640L9.2 | RP11-713C5.1 | RBMS2 | IDH2 | SBF2 | RPL23 |
| TNXB | MRPL17 | CNOT3 | CD55 | TUG1 | MAGT1 | IRF4 | LINC01094 | AC104699.1 | DNM1P46 | GBP3 | C1ORF200 | GJD4 |
| LY6D | EFS | SSR2 | CDC25A | SMIM15 | ZKSCAN8 | CIITA | RHOH | CNNM3 | RP11-158I9.5 | HES2 | DDX6 | RPS10P7 |
| HSF2BP | PCBP2 | MMP11 | TMEM66 | SUOX | AMMECR1 | SAMHD1 | HAAO | ARMC6 | RP1-142L7.9 | PSMD7 | GPR132 | FAM69C |
| SORCS2 | LTB4R2 | RPS17 | THRA | MBLAC2 | CAR5B | CD3G | HLA-J | LGALS7B | TUBA1A | RNASE1 | PIK3C2B | SLA |
| AW112010 | GSK3A | PTTG1IP | IL33 | MTPN | SPRY4 | PLSCR1 | AP003774.1 | DNAJB4 | CHL1 | LEF1 | TGM2 | SV2A |
| VSNL1 | MSRB1 | TRAK2 | CD109 | UPK3BL | NTMT1 | CD40LG | GATA3 | SLC52A1 | MYRIP | BPIFA2 | ALCAM | TDGF3 |
| ADAR | NSUN3 | MECOM | PDZD4 | HOXA9 | MXD1 | SELL | GIPR | RP11-325L12.6 | SNAP25 | EIF2S2 | CLEC18B | TAS2R38 |
| GM14137 | CBR3 | 1110057K04RIK | NDUFA3 | SP3 | GPRC5B | CCR5 | IL10 | PYCR1 | CFD | CCL28 | TRPV1 | FAM196B |
| GSTO1 | ZHX1 | ACOT9 | ZMAT1 | KRT5 | KIF5B | EVI2B | ANXA2 | RSL1D1 | RP4-555D20.1 | GAL | C5 | WDR20 |
| OCIAD2 | ERCC5 | SLC44A2 | POLA2 | ZC3H12A | ZKSCAN7 | CXCL10 | TRIP6 | LINC01128 | NRXN3 | ZBED4 | CD9 | ARMC8 |
| ACSS2 | CXCL2 | MUM1L1 | SMAD4 | ZFAND6 | SLC26A2 | TAGAP | EPHX3 | RP11-768B22.2 | EMC1 | PDE1A | COL5A3 | GCKR |
| CIB2 | EPRS | NABP1 | DTYMK | B630005N14RIK | GART | ICAM3 | LINC00954 | DPPA4 | RP11-506B6.6 | POTEI | ELTD1 | HMGA2 |
| WNT4 | PSMG2 | SMC5 | GM14322 | SLC35A3 | GCN1L1 | SLAMF8 | HLA-DRB5 | PKIG | PIGN | SIGLEC1 | FYTTD1 | SPI1 |
| ALDH1A1 | RPL37 | BCL10 | 4932438A13RIK | TMX3 | 4930427A07RIK | NCF1C | EPB41L2 | PLP2 | LINC00968 | UNC13A | LOC286002 | USPL1 |
| FOS | SYNGR1 | ORAI3 | DOK3 | SEC62 | MCU | PIK3CD | B3GNT8 | DCTN4 | CXCR2 | PRR4 | SH3PXD2B | DOCK3 |
| ASB13 | DARS2 | DHX29 | JMJD6 | PRELP | NFIA | GBP2 | TM6SF1 | CHST1 | MAML2 | DMD | TMEM86A | HFE2 |
| EDN1 | FAM73B | TMSB10 | ABHD1 | HMGN5 | DBNDD2 | FERMT3 | ARPC2 | RBBP4 | ADAMTSL3 | KCNQ2 | JAK3 | BIK |
| TMEM176B | RASD1 | MYL6 | HIST1H3F | CLU | FGFR1OP2 | GBP4 | TMEM154 | MYBPC2 | WIPF3 | SERPINI1 | MAST3 | HERC6 |
| CEP250 | RAB3B | CSF2 | LFNG | CHM | IER2 | SIRPG | NONO | DOK5 | FASN | PRRC2B | MTHFS | PIAS3 |
| LY6A | MRPL36 | TMEM253 | FSD1L | POLR3G | MED13 | GPR171 | IGLV8-61 | RPL5P34 | PPAP2A | RFX3 | C17ORF86 | GLYCTK |
| PSCA | WDR61 | PSAP | 43344 | RAP2C | SPNS1 | SPI1 | TRAV25 | COL14A1 | IGDCC4 | MTHFR | GPC4 | C20ORF134 |
| IFI202B | TMIGD1 | UMPS | CCNE1 | FOS | ADCY3 | TAP1 | GJC2 | DMKN | ADCY6 | PRRC2A | C9ORF9 | ITGA9 |
| MROH4 | CCDC66 | CSRNP1 | MKS1 | TOB1 | MAP2 | DOK2 | HCG4P7 | RP11-13K12.1 | RNF207 | PARP14 | GMCL1L | DARS2 |
| DUSP18 | DOCK6 | BCCIP | PHKB | HP | HLTF | BCL11B | GRAP2 | IL1B | PLLP | GAS1 | REEP5 | IL7R |
| PLTP | 2410006H16RIK | GTF2IRD2 | ZFP280B | RTP4 | GALNT18 | LINC00892 | RP11-89K11.1 | RP11-293M10.6 | TMEM92 | AGPAT5 | C9ORF144B | DCAF5 |
| KRT5 | TSEN2 | GAL3ST1 | CLIC4 | PPP3R1 | TMSB10 | ZNF831 | AL035610.2 | FGF14 | TULP2 | VSTM2A | TMEM50B | CCDC102B |
| LY6E | CREBRF | GPX8 | ZMAT3 | LACTB2 | PDHA1 | KLHDC7B | VNN1 | RP11-16K12.1 | RP11-305D15.8 | SNTB1 | RNF220 | CDSN |
| WFDC12 | NRIP3 | TMEM97 | DMKN | RAB10 | PISD-PS2 | RP11-284N8.3 | LHFP | COMP | RP11-116O18.1 | UBE2L6 | SAMD9L | CYP4F8 |
| GPRC5B | SUDS3 | PRKD2 | XPO1 | TRIB1 | PHF20L1 | FAIM3 | IGHJ2 | FSTL3 | LPHN3 | RRM2B | BST2 | DLK2 |
| PNPO | DDRGK1 | ECHS1 | PIK3CG | LYSMD3 | SNAI2 | MYO1F | RP11-686D22.8 | ESR1 | CETP | GNRHR | FBXW10 | FKBPL |
| PIAS3 | PLCB3 | MAPK8IP1 | GRAMD3 | LTBP4 | CYP27A1 | LST1 | AC022182.3 | LINC01055 | TPT1 | RTF1 | FHL5 | CAPN9 |
| MAPRE2 | ZFP579 | RAI1 | MANBA | CBFB | TXLNA | TLR10 | IGLV1-36 | HHIP-AS1 | NPEPPS | VPS37A | KRT20 | HAPLN3 |
| DCN | FGD1 | B3GALT2 | SNED1 | ITPRIPL2 | TATDN2 | TYMP | STK17A | TPBG | LINC01116 | PSD | LOC100130015 | IL20 |
| TAPBP | GLTP | ADCY3 | TPD52L1 | SOX2 | ANKRD16 | NCF4 | HLA-DQB2 | NEURL3 | ATP5A1 | CDKN1A | LYL1 | MESP1 |
| THBS1 | PKMYT1 | NCAPH | STAG1 | EPM2AIP1 | ATP2C2 | HCP5 | B3GNT3 | PLXND1 | METTL2B | AKR1C2 | P4HA1 | EPHA1 |
| KRT4 | INTS1 | OPN3 | CTSC | H2-Q4 | NLRP10 | LGALS3BP | RPL7L1 | S100B | RARB | TP53INP1 | TSC22D1 | ZNF416 |
| FOSB | BC030867 | NOLC1 | CERS6 | CLOCK | FRK | DOCK2 | ANP32E | TRPM5 | RP11-690G19.4 | ARMCX2 | ZBTB45 | LOC146880 |
| 2210013O21RIK | VAMP8 | NT5C2 | B3GNT7 | KLF10 | USP35 | TCL1A | GTSF1 | DKC1 | WNT4 | C19orf66 | EVPLL | ZNF71 |
| PMM1 | BACE2 | 9130019O22RIK | CLN6 | SLC25A24 | F3 | IGLV1-40 | CH25H | GNB3 | SNHG18 | SDS | RGS9 | SMAD5 |
| 8430408G22RIK | GLIPR1 | TRAF3IP2 | TTC30A1 | ITGA2 | NOD2 | PTPN22 | STAT2 | TMEM40 | KIAA1614 | ZNF862 | DDX42 | GLYATL3 |
| ALOX12E | LAMTOR4 | MKRN1 | EDN2 | HAS3 | TUBGCP6 | GZMK | TNFSF11 | S1PR5 | RP11-435O5.2 | MIEF1 | MC1R | LRRC10 |
| TRIM34A | IQCC | MYPOP | FURIN | CYP51 | FBXO30 | IRF9 | PLEKHO2 | FAIM2 | HOXB-AS3 | UBE4B | TPD52L2 | UBASH3B |
| PRKCG | NME3 | DENND2C | H60C | LAMB1 | PSMC6 | CTLA4 | RP11-283G6.5 | PIGM | CD59 | B3GNT3 | HVCN1 | ZBTB42 |
| UGT1A6A | 2010002M12RIK | ATXN1L | TMEM203 | PIGR | SMIM7 | PLA2G2D | TRAV1-2 | LURAP1 | DMRT2 | GLP1R | ADCY3 | PPAPDC1A |
| MVB12B | FAM212B | BMF | HAGH | KLHDC8B | LIX1L | PLAC8 | FMO2 | GMCL1 | CELF2 | ZSCAN29 | GLT8D2 | 43529 |

### Table S1. Significantly differentially expressed genes (EZH2 Signature High vs Signature Low, p<0.05) continued.

| | | | | | | | | | | | | |
|---|---|---|---|---|---|---|---|---|---|---|---|---|
| NMI | FAM188A | EFCAB4B | FMNL2 | CHMP1B | XRCC5 | KLHL6 | CERK | PYDC1 | ENOPH1 | VGF | ZNF488 | P2RX1 |
| MSLN | DDX24 | ELMO3 | BLM | TMEM176A | PATZ1 | DDX58 | HLA-C | AC068580.6 | AOX1 | PRED62 | DOK1 | SDCBP |
| CCNG1 | UBQLN2 | ROR2 | FBXO36 | HPGD | SLITRK6 | KLRB1 | ITGB6 | CCDC82 | PRICKLE2 | SRCRB4D | GIPC2 | SSPN |
| ADAM8 | ZFP93 | BIRC5 | TMEM106C | CCNC | G3BP2 | HCST | SLC25A42 | LINC01082 | HAR1B | FBXO32 | RAMP3 | RASIP1 |
| ZC2HC1C | CBLN3 | SORBS2 | 9630033F20RIK | USP18 | EIF2S3Y | ITGB2-AS1 | IRF7 | SCARB2 | STEAP4 | PPM1F | LANCL3 | VDAC2 |
| PTPRS | N6AMT2 | KIF2C | PRSS22 | HSPG2 | MAPK11 | HLA-DOA | TSPAN4 | RP11-445F6.2 | DCAF7 | NLGN4X | GSTM4 | C17ORF107 |
| JAG2 | COPS7A | DNLZ | CSDE1 | FAM45A | PRPF40B | BLK | SLC6A6 | TWIST2 | ASB2 | C1orf167 | PIGR | LOC100128023 |
| OMP | SNHG4 | ROMO1 | UBE2N | CPNE3 | CCDC109B | TRAT1 | ADAMTS10 | PLAC9 | OSM | CRABP1 | LIPE | LOC220429 |
| ACSBG1 | VPS41 | RAD51AP2 | HMGN3 | SLC20A1 | CATSPERD | PTPRC | STOM | LINC00299 | RP11-736N17.4 | CDR2 | PTH1R | NPAS4 |
| PSMB9 | RXRA | SCO2 | RNF4 | LIMS1 | GM12603 | TNFAIP8L2 | DNM3OS | CDH23 | CLCNKB | ABCA2 | CDK18 | VILL |
| SEPP1 | 1110007C09RIK | KDELR3 | BPGM | RNF11 | POLR1B | FCRLA | 43160 | C9orf135 | DDI2 | IGF1R | SLC6A3 | SLC25A33 |
| L1CAM | PACS1 | BRE | INTS3 | BMI1 | GPC2 | LILRB1 | FDCSP | GSDMD | CCDC42 | MRC2 | AMZ2 | ZNF618 |
| ID1 | ZFAND5 | USP31 | GM17727 | THOC2 | NT5DC3 | LY75 | DTX3L | ALDH1L2 | DPH5 | APOB | C19ORF63 | C12ORF52 |
| MAPK11 | ZFP874A | RAB3D | PRKAR2A | RHPN2 | SLC39A8 | VAV1 | MYEOV | AICDA | NTN1 | GPR20 | ERO1LB | CARD14 |
| APOF | POLR3E | CDC16 | CTPS2 | LY6A | JADE2 | RASSF5 | USP18 | CATSPER1 | EML6 | APOC1 | IGF1R | ZNF737 |
| EIF5 | RAB38 | NUAK1 | MRPL42 | RAD23B | DEPDC1A | PNOC | EMR2 | THSD7A | RP11-146N23.1 | OTUD7A | LOC100271722 | DLC1 |
| IIGP1 | CCND2 | SEC62 | EVX2 | ZDHHC20 | NDUFA2 | PLEK | AC096579.13 | KDM1A | RPSAP54 | PTCHD1 | MOBKL2C | MSRB2 |
| EXOC4 | SLC35C2 | TSEN34 | PSMG3 | ZFAND2A | PIM3 | SH2D2A | MRC2 | EHD2 | TRIM29 | CBLN2 | C6ORF154 | FLVCR2 |
| PYGL | A330074K22RIK | MFSD7B | UFC1 | MACC1 | YAF2 | LINC00426 | AKR1C1 | LINC00854 | DRD1 | C1S | ELK3 | PDE3B |
| LANCL1 | WFDC15B | NUP160 | SPECC1L | PTMA | EDARADD | IKZF3 | TGFBI | TCP1 | GSTZ1 | KNDC1 | TRIM32 | ENTPD3 |
| REEP5 | ARHGEF2 | GSTT3 | GRTP1 | SYAP1 | SLC31A1 | RP11-1094M14.8 | RP11-812E19.9 | LAMB3 | AC073115.7 | TACSTD2 | CFHR5 | TUBA3D |
| COL4A1 | GTPBP6 | MITF | SRR | RLIM | TAZ | CCL19 | GPR141 | ACTA2-AS1 | RP1-40G4P.1 | EPHA3 | KCNK13 | MFGE8 |
| FOXM1 | ALDH16A1 | LOC106740 | LIMK1 | TSPAN12 | 43723 | CXCR4 | IGHV3OR16-6 | OMG | CACNA1D | LARP1B | MRVI1 | RNF157 |
| CRLF2 | TIMELESS | MPI | TPP1 | ARHGAP5 | RHBDD2 | NUGGC | IGHV3OR15-7 | HCAR2 | BAIAP2-AS1 | MT3 | S1PR4 | H1FNT |
| CAPG | NLN | SERINC1 | HADHA | TRIM59 | | RCSD1 | CD101 | DPYD | TSG101 | POTEJ | SIX5 | ALAS2 |
| SPINK5 | TPM4 | SMYD5 | 2610203C20RIK | CPEB4 | | MICB | S100P | PRDM1 | THTPA | PZP | SLC47A2 | LRRC8E |
| TMEM8 | GATA3 | PCTP | MRFAP1 | PCMTD2 | | RP5-1028K7.2 | INSL3 | RP11-556E13.1 | CTC-327F10.4 | AMER3 | HCG27 | RCOR2 |
| PLEKHG2 | JMJD8 | JDP2 | GPALPP1 | THBS1 | | IGKV1-27 | ANXA6 | IGBP1 | MGC16275 | ADORA3 | OMD | RBKS |
| TOR3A | FAU | MYO1B | CASD1 | DRAM2 | | FYB | ADAMTS6 | RP11-973H7.4 | KLRC4-KLRK1 | BRI3BP | PRR7 | LRRC31 |
| IL18 | UBP1 | IRF2BPL | GHR | XK | | TRBV20-1 | DPYSL2 | LINC01353 | KRT17 | 43163 | ZNF707 | |
| IFITM1 | CDC42SE1 | HSPA4 | PDZK1 | ICAM1 | | APOBEC3D | AC093063.2 | HTRA3 | KLF2 | ITIH4 | ZBTB3 | |
| ARG1 | CROCC | TMPRSS11G | LHFP | RAB20 | | BTK | CP | ADORA3 | NTF4 | IFITM10 | HCN3 | |
| GBP2 | HOXA3 | PAPLN | LRIG3 | RAB18 | | CD79B | RP4-742C19.12 | ELK3 | OR52K3P | IRF9 | SFRP5 | |
| MRPL41 | TMSB4X | FOXQ1 | MRPS7 | SERINC1 | | CRTAM | ARSI | IGLV2-28 | IER5L | PSCA | FPR3 | |
| SOCS3 | TMTC4 | PML | MAL2 | ZBTB33 | | LINC01215 | CNTN2 | IGF2BP2 | PLEKHG2 | RUFY4 | IL12RB2 | |
| PRPS1L3 | HYOU1 | SMCR8 | CCDC34 | SCAI | | HLA-DQA1 | DDO | REPS2 | LRRC16B | SPATA13 | HLA-H | |
| MREG | TMEM108 | RAD51 | HOPX | PURB | | IGHM | MS4A4E | RP11-817I4.1 | ZNF697 | ZNF324 | LMF1 | |
| EDA2R | POLE2 | 4632428N05RIK | CALM3 | UTY | | NKG7 | DUSP2 | TRIM71 | FAM35A | SLC6A15 | LOC348926 | |
| ERAP1 | FRAS1 | SIPA1L2 | MARK2 | BMPR2 | | GMFG | TRBV14 | TMEM200C | KLRF1 | RB1CC1 | LOC55908 | |
| EEF1B2 | PSMB8 | MRPL12 | ACOT8 | RPS6KA3 | | LRMP | TOR4A | IGKV2D-30 | KRT18P28 | FAM21B | WFIKKN1 | |
| CLDN2 | CPPED1 | 5930430L01RIK | NUDT19 | CLCA1 | | TNFSF13B | PRRX1 | RP11-832A4.7 | NIPAL4 | ETV7 | METTL7B | |
| FBXO9 | HAUS4 | ARAP1 | SNX12 | FMR1 | | TNFRSF13B | DFNA5 | PTGES3P2 | POU3F1 | RASSF6 | FOXS1 | |
| CLCA4 | IMPA1 | PATL1 | ZFP874B | CHMP2B | | IL2RB | IGLV3-19 | PRELP | EIF2D | FAM120A | CDK5R1 | |
| SPEF1 | NLGN2 | FAM102B | PGLYRP3 | IRGM1 | | IGHG3 | ANKRD44 | COX16 | NTRK1 | SORBS2 | SNORA71C | |
| CYP4V3 | OS9 | TMED5 | PNRC2 | ZXDB | | AIF1 | PINLYP | VSIG4 | LRRN2 | TNNT2 | TLR5 | |
| ZDHHC20 | EBP | CLCC1 | SLC25A30 | HSDL2 | | IGLV1-44 | ACSF2 | CTD-2377D24.6 | COL5A3 | ZNF37A | NR3C2 | |
| COX7A2L | MED11 | RANBP9 | CCDC14 | MBNL3 | | CYBA | TVP23A | OLR1 | SCARF1 | SLC3A2 | PLEKHG1 | |
| SYNM | SLC1A4 | HSH2D | CCNB2 | STRN3 | | NAPSB | MFAP4 | RP11-295M3.4 | WDR89 | TACO1 | ART5 | |
| ADCK3 | LAPTM4B | NEK7 | RHBDD3 | SKINT3 | | TNFSF14 | IGHV3OR16-9 | GAB2 | RP11-95M15.1 | ABCA12 | TXLNA | |
| KRT23 | MITD1 | FLNA | IFT172 | ZFP207 | | NCR3 | HCG4B | RP11-67C2.2 | RP11-736N17.10 | RYR3 | AKNAD1 | |
| H2-Q4 | HYPK | POLQ | PGAP3 | SLC9A4 | | TNFRSF1B | CCDC80 | FNIP2 | TXNL1 | RPRD2 | PAQR7 | |
| GPR115 | RALGAPA1 | PTPDC1 | EPS8L2 | RNF2 | | SLA2 | CYP24A1 | FRMPD4 | SLC7A3 | OASL | GNPAT | |
| FAM118A | ARHGEF3 | CC2D2A | CENPJ | KRAS | | SPN | RP11-686D22.4 | SLC12A4 | UPK1A | CD163L1 | GDF11 | |
| BMP2 | TCF3 | NME4 | PGK1 | LGALSL | | WDFY4 | IL2 | FCHSD1 | SOGA1 | CTSK | SAP30 | |
| PLET1 | TMEM69 | RBMS3 | EDC4 | YWHAZ | | CYTH4 | AC013264.2 | A2ML1 | RNASE7 | POMZP3 | HOXC13 | |
| MGP | PSRC1 | MTCL1 | PGM2L1 | ABLIM1 | | FCRL1 | SH2B3 | LONRF2 | CCT3 | LAMC2 | IKZF1 | |
| SERPINB2 | FABP6 | 2410015M20RIK | FASTKD2 | GOLPH3 | | FCRL2 | UAP1 | MAPK6 | HTRA1 | ESPN | LRRN2 | |
| SLC16A13 | CSAD | ALG5 | 1700023F06RIK | NEBL | | TRBV5-1 | DNM1 | DANCR | CTD-3193K9.3 | CXCL9 | NPEPL1 | |
| CHAMP1 | SYVN1 | 1810026B05RIK | LGALSL | MIER1 | | EVI2A | BMP3 | PLA2G3 | DHPS | GPX8 | PLEKHG4 | |
| LPHN1 | OXSM | MEGF6 | KLC2 | IDS | | EBI3 | MIR155HG | CMTM7 | AC044907.1 | GATM | RPS2 | |
| TRP53I11 | CDKN1A | CSNK1G3 | SNRNP27 | ORC4 | | WIPF1 | TNFAIP8L3 | C2orf40 | LINC00323 | SEMA3D | CEP110 | |
| 1110038B12RIK | D17WSU104E | RHBG | SP8 | TRA2B | | LAX1 | ADAP2 | RP11-426C22.4 | ADRB3 | TECTB | COL18A1 | |
| MMP20 | PPL | ZFP438 | PNISR | ALDH3B2 | | SLFN12L | ARHGAP23 | RPL10A | LINC00689 | ZNF446 | PLEKHM2 | |
| SYTL2 | MEMO1 | MIER2 | BFAR | GLTP | | IFI16 | EGFL6 | HEPH | SPEG | UBE3A | PAK2 | |
| PRNP | 5730559C18RIK | HS3ST1 | GM5122 | CHL1 | | HERC6 | STAB1 | PRAM1 | MPO | NOVA2 | CD163L1 | |
| SURF1 | DNTTIP2 | SLC3A2 | AI854703 | ACBD5 | | DAPP1 | PPIC | STEAP3 | RASSF2 | IL13RA2 | ACACB | |
| EIF2B2 | FAM132B | DNM1 | MAP3K9 | BNIP3 | | TRBV19 | CDX1 | ADAMTSL2 | SLC25A21 | KAZN | PTPRB | |
| PDXK | RPL13A | CIPC | GLCE | WBP5 | | NCF1B | CTD-2325B11.1 | JAZF1 | KDM4B | PDIA4 | NFKBIE | |
| EME2 | 1700008J07RIK | ENHO | NAA60 | TMEM68 | | TRIM69 | LTBP2 | PGF | CSGALNACT1 | KIAA1147 | CLSTN2 | |
| CD97 | SHPK | SETBP1 | GART | NXT2 | | APOBEC3H | RP11-395L14.13 | PFKFB3 | LINC00544 | NT5E | LOC283663 | |
| DDX39B | PLP2 | CSNK2B | HID1 | CCDC71L | | P2RY10 | RP11-523O18.5 | A2MP1 | RP11-535A5.1 | INTU | RBM7 | |
| GIPC2 | SNHG3 | NDUFB3 | TRIM11 | EGR1 | | SH3BP1 | FCHSD2 | ZMYND15 | CTD-2026C7.1 | CD160 | CIITA | |
| IL3RA | GRID2 | GTF2I | FBXL15 | NEDD9 | | C1QA | SLFN12 | ZNF610 | ATP1B1 | CIC | LTB | |
| RRM2 | SCLY | PGAP1 | NASP | SYT16 | | FAM78A | AC005083.1 | RP11-212I21.5 | TTC7A | SRD5A1 | PGK1 | |
| NOTUM | LTBP3 | SLC18B1 | BCL9 | ABLIM3 | | HLA-DRB1 | ISL2 | CADM3-AS1 | RP11-206M11.7 | ERBB4 | HEBP2 | |
| CTSD | DDT | MAPRE3 | SNAPC3 | 43710 | | FCRL5 | CTD-3035K23.7 | IGHV3-75 | GLYR1 | C1orf51 | DPH3 | |
| FGFR3 | IQGAP2 | MRAS | GM17762 | JUND | | SP140L | LAIR2 | CCDC33 | CYB5R2 | SLC7A2 | FAM90A1 | |
| E2F8 | 9130401M01RIK | 9930014A18RIK | ASCC3 | NCCRP1 | | ITGB2 | DOK1 | PRODH | VKORC1L1 | ADIRF | GSG1L | |
| CGNL1 | EIF3F | MRRF | EXOC2 | CHORDC1 | | HAPLN3 | PHLDB1 | HABP4 | RP11-60L3.1 | CACNA1E | FAM20A | |
| 4930452B06RIK | KLHL9 | PTGER4 | KDELC2 | TNFRSF12A | | HCK | OXER1 | CH17-360D5.3 | HLA-S | HRH3 | SLAMF1 | |
| IGSF9 | MRPS16 | NFKBIA | RANBP2 | ZRANB2 | | GPR55 | ODF3B | SLC24A4 | PLA2G12A | SPIDR | GATM | |
| EIF2AK2 | TPRGL | RPL23 | NPNT | HES1 | | NCKAP1L | FSCN1 | SLC14A1 | BMPR1B-AS1 | B3GALT1 | NOXO1 | |
| GLTSCR2 | AGRN | PPIP5K2 | CCDC127 | HELZ2 | | TRAV13-1 | RP11-402J7.2 | GPR34 | HID1 | KDM6B | RAI1 | |
| CHML | NUP205 | RACGAP1 | MAP7D1 | RPS28 | | PARP9 | SCGB1A1 | ARL14 | ZADH2 | RDH16 | LOC115110 | |
| GDA | AUTS2 | MIF | METTL10 | RB1 | | GAB3 | PLS3 | ERAP1 | CNTRL | AK4 | SLCO3A1 | |
| FUT2 | SAMHD1 | COMMD1 | MLLT10 | CSF1 | | IL7R | MS4A4A | ZFY-AS1 | NGFR | LAP3 | DDX60 | |
| ALAS1 | RARG | RGS7 | HAX1 | CREBZF | | FLT3LG | MGAT3 | RHEBL1 | SLITRK6 | PTCHD2 | GPR109A | |
| RPS19 | GAL3ST4 | SPTSSA | EVA1B | ITPR1 | | IGHG4 | CYP21A2 | RP4-647C14.2 | RPL14 | ORM1 | KCNK7 | |
| THUMPD3 | FAM118B | GPAA1 | BLOC1S2 | SC5D | | IGHV4-39 | CNN2 | RPL22 | RP11-448G15.3 | WDR17 | CD4 | |
| UBA7 | TRPV4 | IL22RA1 | MRPS17 | HIF1A | | BTN3A1 | LUM | TMEM18 | RP11-141M3.6 | GSTP1 | LOC100128842 | |
| IMPDH2 | KLC1 | LDLRAP1 | APIP | CREB1 | | MEI1 | RP11-834C11.7 | PGAM1P5 | SDS | FBRSL1 | DKK2 | |
| FANCA | ZFP668 | MFSD12 | RRAS | 2810417H13RIK | | THEMIS2 | IGHV1-67 | GPR133 | SPTA1 | FAM189A1 | IL8 | |
| SGSM1 | PITPNC1 | AI661453 | TMEM74B | PSIP1 | | TMEM156 | RP11-117D22.2 | RP11-16E12.2 | CAMKK2 | CEACAM4 | LOC729991-MEF2B | |
| TRIM12A | RNF169 | TNPO2 | DHX8 | SNORD47 | | PSMB8 | FAM26E | METTL7B | CANX | MAML1 | NT5E | |
| ST3GAL4 | ZFP692 | RAPGEF3 | HPGD | RASSF10 | | FXYD5 | IGHV3OR16-8 | PRSS45 | CLEC1B | TTYH2 | PTPRQ | |
| SERPINA3H | FIBP | CEP170 | ATG5 | UAP1L1 | | AMICA1 | ZEB2 | BYSL | RP11-1260E13.4 | AMDHD1 | TMEM180 | |
| INTS4 | MCPH1 | PTPRJ | H2-T24 | MYEF2 | | C11orf21 | TRAV36DV7 | GYPE | ORC5 | NAMPTL | ZNFX1 | |
| PLS3 | CDH3 | GPR137B | DSG4 | SACM1L | | IGLV1-47 | GOLPH3 | RGN | KIAA0556 | ZP3 | CCNI2 | |
| AKAP12 | RPL22 | SERP1 | SYNE1 | KDM7A | | IGKV3-20 | MRAS | PRDM16 | LDOC1L | CDO1 | PODN | |
| SLC12A2 | MGAT4B | TVP23B | TMPRSS4 | CCNG1 | | TRBV25-1 | ELP2 | LL22NC03-24A12.9 | CHST4 | C19orf71 | RAB3IL1 | |
| ODC1 | ERO1L | NAIP1 | ZFP35 | ZFP946 | | LAIR1 | COL5A1 | PF4V1 | TCAM1P | TSC1 | C1ORF54 | |

### Table S1. Significantly differentially expressed genes (EZH2 Signature High vs Signature Low, p<0.05) continued.

| | | | | | | | | | | |
|---|---|---|---|---|---|---|---|---|---|---|
| COL4A5 | BNIP3 | TMEM194B | KDM3B | TPD52 | IFI35 | SERPINE2 | GJB5 | CCT4 | IDO1 | NCRNA00164 |
| CYP27A1 | CYP2B10 | EXOSC1 | GTF3C2 | EYA2 | TNFRSF9 | HLA-K | COPZ2 | KIAA1244 | TGS1 | EXOC3L |
| NEXN | TTC27 | SLPI | KANSL3 | IFITM3 | PLCB2 | CORO1C | CASP4 | PAIP2B | MTMR9 | IFITM3 |
| SMIM4 | ZFP719 | SLC26A11 | BICD2 | AKAP12 | CD226 | ADAMTS7P4 | RPS5 | RP11-226L15.5 | DLGAP2 | C1ORF203 |
| TMEM209 | PKP1 | STX8 | TANC2 | DUSP4 | IGHV3-21 | CTSB | PURG | CAMK1D | TNFRSF10B | COMP |
| ANKRD1 | PLSCR2 | NSL1 | AGAP3 | DEK | CD180 | VWA5A | MIR4539 | PHLDA3 | VMAC | ENO1 |
| C330018D20RIK | ARRDC1 | TGFB3 | KCTD18 | KRT19 | GPR114 | POSTN | LYPD5 | CPT1C | DSTYK | DMXL2 |
| TRP53INP1 | TANGO6 | UBLCP1 | LMO4 | RUNX2 | BCL2A1 | TRAV13-2 | AP000432.1 | UBQLNL | MT-CO3 | ZNF557 |
| SEMA3F | FAM101B | YWHAG | MBOAT7 | S100A3 | GPR65 | KCNJ5 | SCAMP1 | PPIP5K1 | WRN | CLEC4E |
| MYO6 | PDCD2L | PRAMEF8 | UBE2H | TGM5 | IL4I1 | LMCD1 | TBC1D16 | ENPP7P11 | KCNH7 | C14ORF126 |
| TMIE | CHAD | CDK2 | ZFP652 | RSBN1 | FAM159A | SYK | NCAM1 | NUDT19 | CRB1 | NAA40 |
| OMA1 | SOGA1 | RPS27L | CBX2 | RP2H | EXOC3L4 | AEBP1 | SIRPD | CIAO1 | 43348 | ARMC4 |
| ATP11B | ARL5A | ACOT2 | TIA1 | KANK1 | STK10 | C5AR2 | TRAV24 | RGMA | DNAH8 | KIAA1908 |
| 2210407C18RIK | COMMD6 | BCL7A | CNDP2 | GCNT3 | SPIB | AP001046.5 | BCL2 | MIR143HG | DUSP15 | GAPDHS |
| DST | PSAPL1 | RAB17 | H2AFY | VBP1 | LGALS2 | RP11-170L3.7 | YRDC | CTSZ | FAM19A2 | GPRIN1 |
| NOL8 | MAPK12 | ZFP516 | UBA52 | PPFIBP2 | FCER2 | RP11-219E7.1 | RP3-453C12.14 | MPZL2 | CSPP1 | NCRNA00188 |
| CLDN7 | GNAL | SMC6 | DGAT2 | KRT78 | CXCL9 | CCL21 | WFDC1 | RP1-29C18.9 | NUPR1L | PODNL1 |
| TRIM25 | TMEM106B | SAP30 | GLT8D2 | DSC3 | IGLV6-57 | NFAM1 | DENND1A | TNC | SELENBP1 | RASD2 |
| TEX26 | CCDC57 | PLIN2 | ANAPC13 | YME1L1 | LPAR5 | DLGAP1-AS5 | COQ7 | RPL7P23 | METTL2B | CH25H |
| NR4A1 | GM19557 | MRPL51 | FOXJ2 | METAP2 | IGLV1-51 | EPHA2 | AGBL2 | RP11-466H18.1 | GOLT1B | CYP2D7P1 |
| GPATCH4 | EGLN3 | HAP1 | PRR15L | MSMO1 | CTSS | TIMP2 | EPHA8 | PHF19 | RNF144B | BTN2A3 |
| WTIP | SIGIRR | CMTR2 | 1810037I17RIK | TFG | SP100 | SRPX2 | DLEU7 | NMNAT1 | SLC44A1 | LOC341056 |
| MPDZ | FNDC1 | CTDSPL | COL18A1 | TMTC3 | TMEM71 | CNTNAP1 | PTPRQ | FMO6P | BPIFB2 | AGTPBP1 |
| KRT8 | SNAI2 | APPL2 | ALDOC | CBLL1 | TESPA1 | APOL2 | TRAV38-1 | CCR3 | PCMTD1 | MAL |
| GUSB | HADHB | GABPB1 | CDC25B | EID1 | IGKV3-15 | RGL4 | IGF2 | TFDP1 | UBXN2B | RNF212 |
| IFI27 | EIF4E3 | DHDDS | TTC7B | ST3GAL1 | TNFRSF13C | DSE | FZD7 | YWHAQ | NEIL2 | C9ORF93 |
| FAM115C | RPS13 | ABCF3 | METAP1 | LY6C1 | PRF1 | GFI1 | LRRC15 | UTP14A | MMP25 | CLK2 |
| UAP1L1 | PDZRN3 | STK39 | NUP210 | MIR703 | C1orf162 | WNT6 | C1QTNF1 | PTGR2 | RET | COL6A2 |
| CLEC2D | WFDC3 | CELF5 | DPY30 | SQRDL | CCL22 | GPNMB | KCNJ1 | TBC1D2B | RASGEF1A | DBI |
| RPS28 | ACAT1 | FAM129C | FAXC | SNX10 | CD8A | LINC00857 | TUBB6 | FXYD1 | MTG1 | SNHG5 |
| AOC2 | PDP2 | RALGDS | PIGM | ABHD17B | XCL2 | IGKV5-2 | CEBPE | SELM | NDST1 | ZNF331 |
| ID2 | FN3KRP | TMEM147 | LYPD6B | LAMP2 | ICOS | CYP21A1P | C15orf41 | TFE3 | BICC1 | GPATCH2 |
| 5430419D17RIK | ALG12 | ARHGEF17 | LRP5 | CSNK1A1 | MILR1 | LRRC8C | COL6A2 | CTD-2252P21.1 | NSG2 | RFTN2 |
| RAD9B | COL8A1 | CLDN3 | C77080 | KCNK6 | NLRP1 | TRBV7-6 | TNFAIP6 | CCDC85C | SCGN | ATP6V1E2 |
| SULT1A1 | CDH17 | BCL2 | TTK | PSME2 | CNR2 | HLA-F-AS1 | DCAF16 | IL17REL | DPYSL4 | C16ORF70 |
| ZFP414 | GOLPH3L | MRPL20 | WAPAL | TM9SF3 | IDO1 | FAP | FUT3 | RP11-598F7.3 | RPN1 | C6ORF146 |
| PADI3 | RNF181 | RPL38 | SECTM1B | FAM25C | ARL11 | RP4-800J21.3 | CORO2B | STAMBP | LDOC1 | PARP9 |
| XRCC5 | NAB2 | FZD1 | CLNS1A | ABCB7 | HAVCR2 | 43344 | FHL3 | EIF3J | BEX5 | PRKACB |
| RASL11A | GNAI2 | 4933408B17RIK | HS6ST2 | UBE2D2A | IGHV3-15 | FEZ1 | EEF1A1P11 | PPAPDC3 | C14orf164 | TTC27 |
| ADH1 | IGBP1 | ZYX | PIGL | CHKA | LINC00996 | CTB-133G6.1 | PLBD1-AS1 | SBSPON | C22orf26 | C9ORF95 |
| PIK3R3 | NSA2 | E2F3 | NR2F6 | UBQLN1 | ALOX5AP | VENTX | SLC1A3 | TLR4 | ZFHX4 | GOLGA1 |
| CASP4 | RFX2 | 2310033P09RIK | NDUFV1 | DNAJA2 | CXCR6 | OTOF | SERPINB3 | FAM222B | GRAMD4 | HINT1 |
| DTD1 | DIABLO | KDM3A | ZCRB1 | SYNE2 | SAMSN1 | HK3 | ZNF616 | SEMA3A | KLHDC10 | ST5 |
| SMIM8 | IL18BP | BIN1 | STK16 | RNFT1 | CD86 | CD8B | RP3-492J12.2 | RP4-728D4.2 | MAMLD1 | ITK |
| PNP | SAP18 | TIGIT | SH3TC2 | SRSF10 | RIPK3 | IL1RN | ANO9 | MRGPRF-AS1 | HOXA13 | OAT |
| MYBL2 | ANXA8 | NKX2-9 | ZFP39 | GNG12 | FCRL6 | HAS2 | CORO7 | BHLHE40 | LSM6 | ARMC10 |
| 4632434I11RIK | 43354 | DPCD | CSNK1G2 | ZBTB41 | MIAT | ZCCHC18 | LINC00672 | SLC8A1 | CCL8 | SPNS1 |
| SLC19A2 | SPTSSB | SLC2A6 | SNRK | FUNDC1 | TNFSF8 | VTCN1 | CDHR1 | HSD11B1 | RARG | ZNF714 |
| ABCB1B | DCP2 | PPARGC1A | MECR | GABPA | TLR6 | TMEM213 | SLC40A1 | C1orf43 | C5orf24 | GREB1 |
| MBOAT2 | RBCK1 | RAB19 | KRT20 | DDX3X | HLA-DMA | ENTHD1 | FBXO25 | PTRF | COL3A1 | VIL1 |
| 9030617O03RIK | LMNA | KLF13 | SLC7A11 | 1190002N15RIK | LY86 | RTP5 | RNASE2 | GH1 | ALG1L | C6ORF118 |
| CSF2RA | UFD1L | ZFP746 | PCF11 | UBQLN2 | SIGLEC1 | OLFML1 | CYP4Z1 | RP11-120K18.3 | POTEE | EZR |
| MBP | MSL3L2 | KIFC1 | 6530402F18RIK | PERP | IGKV1OR2-108 | NHS | LINC01013 | AGPAT3 | GCFC2 | HES1 |
| SGMS2 | SH3TC1 | RPL27A | BTC | SOWAHC | RP1-47M23.3 | PRKCDBP | ZNF154 | AC103563.5 | REN | MYCBP |
| ZFP395 | RPS9 | NT5DC3 | ARHGEF26 | KPNA3 | NRIR | CYP4Z2P | AC007278.3 | NPM1P27 | HEATR5B | RCN3 |
| CMTR1 | KDSR | PDZD2 | THUMPD1 | MBNL2 | CTC-378H22.2 | TAPBP | TRAV35 | RP11-427M20.1 | NACC2 | NCRNA00219 |
| MAP6 | PRPF40B | C330027C09RIK | ZBTB17 | NAA15 | PATL2 | RASGRP3 | RP11-798K23.5 | HRH3 | CRY1 | MLKL |
| TMEM43 | TMED4 | ANKRD49 | ALDH4A1 | PNRC2 | HVCN1 | SND1 | ZC3H7B | HCG4P5 | TMEM120B | SSC5D |
| SLC6A6 | TONSL | DCAF10 | MPHOSPH10 | A830080D01RIK | TNFRSF4 | PLEKHO1 | DPY19L3 | CTD-2240E14.4 | ABL2 | LOC642852 |
| B4GALT2 | FAM193B | ALDOA | PFN1 | PDCL | CLEC4A | LINC01410 | IGF1R | RP11-705C15.2 | NAIP | FBXL21 |
| H2-K2 | SERPINB11 | 1110004F10RIK | SPC24 | ALDH3A1 | SLC17A9 | IGHV3-60 | WDR86-AS1 | RFTN2 | FAN1 | INPP5D |
| IFITM3 | STXBP2 | TRIM31 | MAGI3 | IKBKE | ITGAX | C5orf58 | IPCEF1 | NECAP1 | IFNAR1 | RSAD2 |
| POGLUT1 | TICRR | AU021092 | TMEM87A | ARF6 | IGHA1 | NT5E | MGC32805 | GNL3 | FAM53C | SEC11A |
| ADK | CNP | ZBTB24 | CEP83 | HAUS3 | IGLL5 | AP000783.1 | PDLIM7 | ADAD2 | IGF2 | FAM78B |
| RPL17 | SLC37A4 | FAM131C | CTNNAL1 | SPRY1 | CD300A | RHBDL2 | JAKMIP1 | GLYATL1P4 | C2 | RHD |
| EML1 | NCOA4 | OAZ2 | PUM1 | PMAIP1 | TRBV7-9 | CACNA1F | RHPN2 | MAN1A1 | TFF3 | ADAMDEC1 |
| TAF1D | KDM6B | SYK | CMAH | TOP1 | FCER1G | ANKRD35 | LPCAT2 | RP11-326C3.11 | CCNG1 | TMEM65 |
| VKORC1 | PRSS8 | GET4 | SMIM15 | LRIF1 | SCIMP | PPP4R1L | RAP1GAP2 | SS18L1 | IL4I1 | HTR7 |
| PRUNE2 | UBE2V2 | PTK2B | GBF1 | FUT2 | IGLV7-43 | SERTAD4-AS1 | MPI | RP11-379F4.1 | TRIM5 | RNASEK |
| TRIM30A | CDKL5 | A530046M15RIK | TM7SF3 | TRIM2 | TRBV6-1 | HEMGN | LINC00582 | IFNL1 | DENND1B | SNORA48 |
| NT5E | FUOM | LLGL1 | COMMD4 | SP4 | IL24 | FOXI1 | ST13 | RPL37 | PWWP2A | PRICKLE1 |
| DUSP1 | TCF7 | NDUFA12 | CCDC50 | ZFP943 | FBXO39 | QPRT | TMEM178B | RP11-686D22.3 | C5orf63 | TUSC5 |
| GM94 | POC1A | RPS6KA1 | 2310009A05RIK | DSG4 | SLC15A3 | TRABD2A | DNAJC5B | CLCN3 | DDIT4L | C17ORF59 |
| TPM2 | PRRC2B | L7RN6 | TBL3 | CEPT1 | CD80 | BGN | TBX19 | RP11-356J5.12 | NAGLU | ZCWPW1 |
| TGFA | HMOX1 | ING2 | D630045J12RIK | TMEM165 | BTN3A2 | LILRA4 | CACNB1 | PENK | DACH1 | FAM82A1 |
| ABHD12 | DMTN | GM16576 | JUNB | CYR61 | CTSW | HLA-DPB2 | LINC01010 | KCTD10 | CH25H | IL3 |
| AEN | CTNND2 | MOGS | BGN | PSME1 | SERPINB9 | SYDE1 | ADCY1 | ZNF521 | C17orf78 | TAS1R2 |
| DLG4 | VPS13C | ELMOD2 | SAR1B | MSI2 | TIFAB | PLXDC1 | HLA-U | CYP4F12 | B3GALT6 | ZNF425 |
| RPP25L | DIEXF | CHCHD2 | 3632451O06RIK | CXCL15 | PLCG2 | OLFML2B | MIR31HG | AF064858.11 | DPT | CX3CR1 |
| 5031425E22RIK | NIFK | MRE11A | UGGT1 | ANXA1 | CERKL | AC021188.4 | GNG11 | NOD1 | EFS | ZBTB39 |
| FAM187B | DNAJC18 | HMGB3 | GM10336 | CNOT7 | RARRES3 | MAP3K12 | CTD-2314G24.2 | PI16 | FLJ27365 | ATP10D |
| GBP9 | MTG2 | NQO1 | SMARCC2 | SDF2 | C1QC | SMARCD3 | COL4A2 | GFI1B | RNF212 | GBP3 |
| 2700099C18RIK | POLR3H | QARS | GM527 | POGK | CLEC7A | HTR7 | LINC01133 | AC005082.12 | LIPA | DLEU2 |
| BCAP29 | SCAF1 | GJB4 | TMEM123 | TCEAL8 | RP11-876N24.3 | IGHV1-58 | SELP | GPR87 | FBXO46 | MRPL53 |
| PNN | SEC14L1 | ANXA1 | RWDD2A | RELB | ITGB7 | HCAR3 | HIPK2 | EDIL3 | CRYAB | PKM2 |
| BTG1 | PKM | PCDHB17 | POLR1A | ME1 | TBC1D10C | IGLV5-37 | HOXD11 | TRMT10A | ACSM5 | FZD9 |
| CYB561A3 | CYS1 | ARHGEF5 | ARHGEF4 | MDFIC | BTN3A3 | IGHV1-69-2 | EEF1A1P9 | MICE | AC127496.1 | OAF |
| UROS | FABP5 | TARS2 | AVIL | UPK3A | IGKC | LINC00475 | ARL10 | RP11-526F3.1 | TAPBPL | PYGM |
| DSC3 | NRARP | 4930414L22RIK | CEP112 | LNP | GZMM | AC145110.1 | TCF21 | KNSTRN | MACC1 | TSHZ1 |
| SMARCA2 | LGR5 | RIC8 | DAAM1 | UBE2E1 | CSF2RB | PADI1 | CENPO | EVX2 | FGFR1 | LRCH1 |
| EEF1A1 | ZSCAN12 | CUL4A | RASSF1 | RAB27B | AOAH | APOBEC3C | MARCKSL1 | 43165 | MAST1 | LOC100130776 |
| SLC15A2 | MRPS18B | WDR46 | DKC1 | UBE2W | TNFRSF17 | NR2F2-AS1 | KCNT2 | GTF3C1 | B3GAT1 | RPLP1 |
| PORCN | SLC12A4 | ZDHHC17 | PRR12 | PLK2 | IL32 | ZNF365 | PMEPA1 | ANKDD1A | SFRP4 | MIPEP |
| MDN1 | GRHL2 | PKD2L1 | EFCAB12 | RDX | ERMN | HLA-DQA2 | LPIN1 | TNFRSF18 | ZFP62 | FAM124A |
| PAPSS1 | EZH1 | PCYT1A | D030025P21RIK | SERBP1 | SLA | CTA-384D8.34 | KCNK2 | HSPB2 | CCDC85C | C22ORF42 |
| PSME2 | MANBAL | TBCK | CLEC2E | TTC36 | LILRB4 | HS3ST3B1 | ADARB2 | LAPTM4B | DNAJB7 | PDCD4 |
| GDF15 | ATP7B | ANK1 | UGT1A6B | CXCL10 | OSBPL3 | MROH7 | RBPMS | ATHL1 | HIPK2 | CLECL1 |
| IFI203 | EVL | MCU | METTL8 | CLDN4 | IGLV2-23 | UCHL1 | RBM15B | E2F3P1 | ELP2 | MS4A15 |
| LOXL2 | GRPEL1 | ZFP317 | VPS54 | UBE3A | CCR4 | CCL20 | C1orf110 | RPL23A | GALE | UPP2 |
| SLC52A2 | DNAJC10 | A530054K11RIK | POLR3B | ROCK1 | HLA-A | CHRNA6 | KIAA0319L | TACR1 | RHD | ZNF878 |
| CENPE | SHISA5 | ANKFY1 | POLR2M | EVPL | TRBV3-1 | TMEM52B | FER1L5 | FOXC2 | ZNF106 | HMGCS2 |

#### Table S1. Significantly differentially expressed genes (EZH2 Signature High vs Signature Low, p<0.05) continued.

| | | | | | | | | | | |
|---|---|---|---|---|---|---|---|---|---|---|
| PXDN | TMEM70 | ZFP169 | PTPLAD2 | FOSB | C19orf66 | EDC3 | NUDT9 | IGLV2-33 | PIGM | LAMP3 |
| ABCC5 | PYGB | RGL1 | KRTAP3-2 | FADS3 | C1QB | PARP12 | RAB32 | DDOST | GLUL | ZNF862 |
| 3830406C13RIK | ASL | PBK | NDUFA7 | ARPC5 | IGLV2-14 | ATP6V0A4 | GFRA1 | SSPN | PRR7 | PADI2 |
| ANKRD35 | HMOX2 | S100A8 | DAGLB | TNFAIP3 | TYROBP | COL1A2 | RENBP | TRPV4 | TNKS | ZNF683 |
| CHIL1 | SLC43A2 | BOP1 | UBE2G1 | 2310002J15RIK | TFEC | RP11-119D9.1 | CTD-2054N24.2 | RP11-131H24.4 | COL16A1 | LIMS2 |
| WIF1 | DDX21 | BTD | ZFP346 | RAB11A | IGHV3-74 | AC018890.6 | IL7 | NES | RPP25 | MTTP |
| RCOR2 | F2RL1 | ITGB3BP | NAAA | TNFAIP2 | LGALS17A | ST6GALNAC2 | CTB-114C7.4 | RP11-108M9.5 | C17orf103 | PBX4 |
| SOX12 | PTRH1 | RNF217 | EGFR | POMGNT1 | B2M | FOLR2 | SLC35F4 | IGHV1-14 | IFITM1 | TXNIP |
| TTC39A | DENND4A | 4933411K20RIK | SUPT5 | AGRN | TRIM21 | RP11-356I2.4 | METTL2A | BDKRB1 | ZNF282 | PROC |
| XRCC1 | FKBP5 | NT5C | PVRL1 | FOXN1 | CXorf21 | SLFN5 | NME4 | KIF26B | SLC6A14 | TAS2R8 |
| METTL3 | MED24 | SOX9 | SIDT1 | 2310007B03RIK | HLA-B | PODN | RP11-1036E20.9 | SCMH1 | TGFBI | RPL28 |
| EGLN2 | FAM65A | KLHL8 | ERCC6 | CPSF6 | BTLA | CTHRC1 | ZNF469 | EDNRA | PROP1 | XKR9 |
| CREG1 | MBOAT1 | SLFN4 | IGFBP3 | CCNE2 | TRPV2 | GABRA3 | SDK1 | AC002480.3 | TBC1D8 | BEX4 |
| EPB4.1L1 | INTS6 | HARBI1 | LY6F | IER3 | IGLV3-1 | SSC5D | MRGPRF | RP11-58K22.4 | TOB2 | AXL |
| FXYD5 | ATG9B | KCNRG | PANK2 | PDXK | FOXP3 | RP11-473M20.9 | CHI3L1 | ABCC9 | TMEM208 | GRIK5 |
| NEIL3 | EIF4H | SYDE2 | TNFRSF10B | EXOC5 | PLBD1 | RAB3IL1 | DDX26B | SEC23B | PCM1 | HSBP1 |
| ZFP750 | DYNAP | WASF2 | KIF4 | AFAP1L1 | CD22 | TRBV4-2 | AKR1C2 | APOBEC3F | ULK1 | MPZL3 |
| DEGS2 | LEMD2 | RPUSD3 | ZSCAN21 | SH3TC2 | PPP1R18 | C16orf89 | ZNF296 | OSBP2 | DSCAM | MSLNL |
| H2-D1 | NDUFAF1 | NFE2L3 | CCDC124 | SMC4 | ARHGAP4 | IGHV1-45 | RNF41 | C19orf84 | TNNC1 | RPL32P3 |
| UBR3 | PDS5B | EYA2 | SEMA3C | ACTR3 | IGKV3-11 | MUC4 | COL15A1 | CTD-2260A17.1 | FTSJ3 | MMP13 |
| NUTF2 | ASS1 | SNW1 | CITED4 | PARP14 | CR2 | EFEMP1 | CCL15 | UBIAD1 | CLN8 | NKD1 |
| ZFP518A | PARP3 | LANCL3 | KIF1B | ACKR3 | TRAV8-2 | AC006273.5 | ARHGAP31 | HSPA9 | IPO9 | NOV |
| SAA3 | TMCO1 | NUDT6 | HAL | ST5 | RASSF4 | CXCL11 | PAICS | RP11-73M7.1 | NLRP8 | SNORA49 |
| BSCL2 | RASEF | PURA | FNDC3B | ATP13A3 | LY96 | GIMAP8 | SMARCC1 | DNAH5 | PAOX | FCRLB |
| TMEM175 | UQCR11 | TMX4 | PPP2R2B | SLC12A4 | TRAV8-3 | ZSCAN4 | RPS2P32 | CYR61 | SYN1 | MYOM1 |
| GFER | ELOVL4 | UTRN | CALCB | IL31RA | RP11-1094M14.5 | PARP3 | CLMP | MEGF8 | ATF5 | SCFD2 |
| NYAP1 | LRP8 | TTC39B | SLC30A1 | ZFP654 | GNLY | TRBV10-2 | GPRC5A | TFAM | PHACTR3 | TMEM120B |
| HNRNPR | ZFP697 | NRADD | ECE1 | DGKD | GAPT | TRBJ2-3 | RPLP0 | RP11-568J23.8 | SERPINA11 | S100A2 |
| BHLHE41 | FAM76B | CAR5B | DNAJC5 | ARHGAP29 | GIMAP2 | KRT86 | ISL1 | ARID5A | PARP9 | HSD17B12 |
| BC064078 | COQ4 | RIC3 | SYTL3 | MALL | IGKV1-17 | TMEM255A | IGKV7-3 | AC017060.1 | PROX1 | KLHL3 |
| PCYT1B | FGFR2 | SCAND1 | AGBL5 | YWHAE | CD244 | CTD-2089N3.1 | AREL1 | RP5-858L17.1 | ILDR2 | LIG1 |
| HSPG2 | 2310040G24RIK | NOS2 | COX6A1 | TSPAN7 | TRBV18 | AC067945.4 | IGKV1OR22-5 | WDR12 | MST1R | LOC388428 |
| OSR2 | HPCAL1 | 0610011F06RIK | RABL6 | BNIP3L | PLD4 | CD28 | RP11-138I18.2 | GOLGA3 | SEC13 | PRKD3 |
| CD14 | RAB13 | MRPL43 | TUBB4B | PLS1 | TRBV29-1 | RAB42 | SERPINB5 | LOXL2 | MARVELD1 | TMEM141 |
| CDK16 | GTF2IRD1 | RPL14 | EFNA2 | ATP11C | TIGIT | RPL7A | RPL13A | SIRPB3P | CAMK2D | TMEM175 |
| CORO6 | CDC45 | PIGS | ZMAT5 | TRP63 | SAMD3 | GNAI2 | GLI2 | EPHA3 | ADAMDEC1 | NFIX |
| JRK | IPO4 | REXO1 | LSG1 | TSPAN8 | GPR174 | LINC00605 | PPP6R2 | RP4-641G12.3 | NFE2L1 | CEP164 |
| DIS3L2 | CCDC120 | NCAPG | YAF2 | CD63 | CTD-2521M24.5 | POMC | RPL14P1 | SIX2 | C1orf27 | OAS2 |
| MNDAL | NCK2 | TNFSF13 | ICMT | MMGT1 | RP11-18H21.1 | TRAV39 | CPQ | FOCAD | FRMD6 | GGNBP2 |
| TGM1 | SLFN10-PS | GM12250 | CISD3 | RNPC3 | IFIT5 | ARRDC2 | IGHV3OR16-10 | ASIC1 | SERPINA1 | TIMM9 |
| NDUFA4L2 | TLN2 | MARVELD1 | H1F0 | DEPTOR | LINC00528 | PPP2R2B | TSPAN7 | RIN1 | IGSF9B | RAB19 |
| LAMB2 | HS6ST1 | RPL11 | PLSCR4 | CRBN | THEMIS | MUC1 | RP11-420L9.5 | RP11-807H22.7 | DNAH10 | TCF7L2 |
| ARHGAP22 | HOTTIP | KREMEN1 | KPNA4 | IER3IP1 | PDCD1 | CYP27C1 | LINC00937 | CCND3 | MMP2 | AQP12B |
| FBXW9 | STARD5 | KRT84 | PXMP4 | METTL7A1 | IGHV3-11 | BACH2 | GPHN | LDLRAD3 | PCK2 | DDX21 |
| TM4SF1 | MIS18BP1 | PCDHGB2 | CXCL15 | NEAT1 | IL18RAP | PDGFRA | WARS | TMC4 | CHGA | C20ORF197 |
| SAMD14 | BAX | CCRN4L | RABEP2 | PXDC1 | RNASE6 | APOE | AC061992.2 | CTC-236F12.4 | TFG | DSC3 |
| 4833423E24RIK | ADAMTSL4 | RERE | MRPS33 | ADAR | TBC1D27 | CCR9 | RP11-817J15.2 | TMEM69 | MAP1B | ACTG1 |
| SAT1 | SESN1 | MRGPRE | FAM199X | TMPRSS11E | IL2RA | NYNRIN | TIAF1 | RP4-737E23.2 | SCARB1 | ADAP1 |
| SPR | AW549877 | TMEM143 | SKIDA1 | SH3GLB1 | REC8 | TRBV23-1 | LRRC10B | RHOG | FBP1 | ARHGAP22 |
| 9530053A07RIK | CDC25C | ADSL | BOC | RNF13 | C10orf128 | DERL3 | EIF3I | JSRP1 | LHFPL5 | ARL8B |
| PSME1 | UROD | PRPS1 | FARP2 | LARP4 | UBD | IGLV3-27 | FAXDC2 | RP4-610C12.4 | KDELR2 | CARD6 |
| HOMER2 | H19 | AHR | SLC29A1 | 2900056M20RIK | RP11-367G6.3 | RND3 | DDAH1 | INHBA | C12orf44 | CASP1 |
| PI15 | PTGS1 | MAP1A | ATP5E | MFGE8 | IGLC2 | NETO1 | NARS2 | DACH2 | PCDH19 | CCL20 |
| CDT1 | CYP26B1 | RBM3 | PPIB | ZFP758 | MSC | MLEC | MAZ | CTD-2020K17.4 | CSF3R | GPR82 |
| ZFP36 | ATM | SEC63 | CAND2 | KCTD12B | PTAFR | COL1A1 | RP11-13K12.5 | RP11-118K6.3 | UBQLNL | HOXD8 |
| PLCD4 | EPB4.1L2 | EIF2B3 | AKR1C12 | CAR13 | PRR33 | BTN2A1 | MREG | RP11-1029M24.1 | AKAP8 | LAS1L |
| LOC100503676 | ARHGAP29 | DOK1 | PIK3C3 | STAT1 | MS4A6A | CST6 | ABCA6 | CRNKL1 | C4orf6 | MTBP |
| DPP7 | BC005537 | AKT1 | GOT1 | MOB4 | TRAV19 | MAL | SLC25A3 | RP11-861A13.3 | SETDB1 | NID2 |
| SMAD6 | FAM98A | WRB | AURKB | LUC7L2 | IGLV2-11 | IGKV1OR22-1 | MYO6 | AC011893.3 | COX4I1 | PRKCD |
| EEF1G | MRPS12 | NDUFAF2 | PCX | IL18 | SLC7A7 | NFATC2 | SOSTDC1 | PTGDS | SNAP23 | PROP1 |
| PAPD4 | ENDOD1 | FBRS | OLFM2 | CCL20 | VPREB3 | SIRPB1 | KRT81 | BPI | SLC28A2 | TUSC2 |
| KRTDAP | VPS8 | SMOX | GM6548 | BMP6 | AC079767.4 | CCDC8 | RP5-866L20.1 | PECAM1 | SH2D1B | UBA7 |
| BID | IRF3 | REEP3 | NDUFS6 | OSR2 | GPRIN3 | SYT11 | PYCARD-AS1 | UBE2G1 | FUCA1 | VDAC1 |
| SMTN | PDGFC | TMEM141 | UBE2Q1 | FAM135A | TRAV8-6 | EMR4P | RAB3D | SLC9A4 | VCPIP1 | C16ORF61 |
| S100G | ERCC3 | SYT5 | PSMA4 | NUFIP2 | RUNX3 | FHL2 | NOX4 | IGHV1OR21-1 | DICER1 | C17ORF108 |
| HOOK2 | PACSIN2 | TRPM6 | 4732490B19RIK | FAM84A | XCL1 | RP11-404F10.2 | TENM4 | SFRP1 | LRRC8B | C6ORF70 |
| SCRIB | RFC4 | NOTCH3 | RPS6KB1 | EREG | IGLV4-69 | TMPRSS3 | BCORP1 | DGKH | PYGB | C9ORF109 |
| DCAF11 | S1PR5 | SAMD10 | FLRT3 | 3830406C13RIK | PDCD1LG2 | SLC26A9 | CEBPB | TRIM31 | CAPN2 | ZNF572 |
| NEDD9 | MESDC1 | HS1BP3 | LIN7C | COL16A1 | CPNE5 | ICAM2 | SPDEF | POLDIP3 | C9orf172 | IKBKB |
| STX2 | SNORD87 | EPN1 | SLC18A1 | SSR3 | KYNU | DHRS2 | CCL23 | B3GNT9 | CCDC144A | HOXD3 |
| FLOT1 | EPHB4 | PLA2R1 | PARP8 | NCOA4 | DOCK10 | EIF3L | TSLP | SCRG1 | SHC4 | RNASEH2C |
| RPL18A | SCIN | MAF | KCTD12 | SP8 | GPR18 | VSIG1 | RP11-180N14.1 | CTH | NDUFA9 | STK40 |
| MPEG1 | HSD17B4 | SMIM1 | WWOX | EGLN3 | CD200R1 | AD000864.6 | RP3-439F8.1 | RP11-805I24.3 | TRPV4 | C20ORF194 |
| ESPN | LAMTOR3 | PEX13 | SLC22A23 | FOXQ1 | RP11-731F5.2 | CYSLTR1 | SH3PXD2A-AS1 | GRIK2 | ZNF687 | GOSR2 |
| SIRPA | CREB3 | RNF128 | SF3B3 | ESD | SIGLEC10 | ITPRIPL1 | RP11-999E24.3 | LARP4B | COL5A2 | GPR20 |
| SERPINA9 | CDC20 | ZC3HC1 | PPP1R11 | GM15421 | LRRC25 | TCF7 | DIO2 | FAM43B | AC091801.1 | ACADL |
| ELL3 | JUP | ZC3H11A | SIK3 | TUBA4A | HNRNPA1P21 | RP11-673E1.3 | EIF4BP3 | PWRN1 | RP1 | ZNF780A |
| RHBDL2 | BBS4 | DIAP2 | 2410076I21RIK | HBEGF | GIMAP4 | GFPT2 | RPL17 | AC244250.4 | ATF6 | DNM1P35 |
| SUCO | 2610005L07RIK | RPS10 | 4921524J17RIK | WASL | CD1D | ECM2 | GRIN2A | NETO2 | SLCO4C1 | GABARAPL3 |
| HSPH1 | IFRD2 | FAM83H | TMEM223 | COL18A1 | C10orf54 | RP5-1071N3.1 | RPS6KA6 | STEAP2 | XKR6 | ZFP28 |
| FAM208B | ITGB2 | PPP3CB | 1810010D01RIK | PRSS23 | CHST11 | RP11-294C11.2 | CMTM3 | HAS2-AS1 | SLITRK3 | ODC1 |
| NEURL1B | KHDRBS1 | MRPL28 | DNAJC21 | RPL37A | NLRC3 | TCF7L1 | RP1-228H13.5 | FREM1 | ABCG1 | ZNF322B |
| LRAT | CTR9 | CHTOP | KTN1 | EIF2S2 | SPNS3 | FGF1 | COL5A2 | INHBA-AS1 | CDH2 | FAM184B |
| ATOH8 | ACACB | CENPV | IGFLR1 | BRCC3 | CTSK | CLIC3 | IFITM10 | RP11-417E7.1 | GCSAML | C20ORF11 |
| TOP2A | NEBL | TBL1X | CREB5 | GPBP1 | CLEC9A | RGS1 | ETNK2 | IL26 | ITPK1 | JPH3 |
| GNB2L1 | WDR18 | TMEM42 | RAB22A | DNAJB9 | DPEP2 | ACSL4 | PHB2 | IGLV5-48 | EPSTI1 | CNIH4 |
| ARHGEF25 | SERINC3 | TNFRSF19 | TANGO2 | HSD17B7 | TRAV17 | MET | IGHV1-69 | SLC30A9 | NRN1 | LOC727896 |
| PSAT1 | HIP1R | LEPROTL1 | TMEM41A | ST8SIA6 | IGKV4-1 | HERC5 | RP11-408P14.1 | TRBV10-1 | PXDC1 | ABI3BP |
| 4930523C07RIK | PRICKLE3 | NDUFC2 | MCM8 | SLC4A4 | IGKV1-8 | GRIA4 | XXbac-BPG248L24.1 | HGF | SCNN1A | ARPC5L |
| PLGRKT | CSPP1 | TBRG1 | CLTB | MGP | TRBV11-2 | P2RY12 | RP11-366M4.11 | STK4 | NCOR2 | CARD8 |
| MAGED2 | GPSM1 | PCDH7 | SPRR2F | LPIN2 | RP11-609D21.3 | RP11-8L8.2 | SPHK1 | CTSA | GBP1 | CNTLN |
| PSMB10 | NADSYN1 | GM16938 | FAM53B | TSKU | TRBV2 | CANT1 | CYP3A7 | COL7A1 | KCNMA1 | LHFPL5 |
| WDR55 | NES | RAB11FIP2 | KPNB1 | PLAGL1 | CSF1 | UCK2 | LINC00924 | AC027319.1 | HSPB6 | OASL |
| DANCR | PRMT7 | 5830418K08RIK | EMB | ANGPT1 | CTSC | CDKN2B | NFKBID | GLUD1 | WDR35 | SLC2A8 |
| GPR56 | RPL7L1 | AGPAT1 | FAM98C | PRKAA1 | LAMP3 | CDH11 | RP11-685G9.2 | BMP4 | PTPRN2 | TRIM13 |
| RBM33 | SELM | ARF1 | ZCCHC17 | LMO4 | MTND1P23 | AC083949.1 | P2RY2 | AF064858.8 | TOR1AIP2 | CYP21A2 |
| LIPA | PCID2 | RGS10 | PHTF1 | MEF2A | SEMA7A | MFGE8 | SLC4A8 | BCHE | PITPNA | MSC |
| UBE2M | ARL14 | VPS18 | FNIP1 | SPNS2 | CLEC17A | ACACA | CLCA4 | GSE1 | ZC3H11A | C3ORF1 |
| ADH6A | TOMM40L | NEURL3 | GM20324 | 1810022K09RIK | IGHA2 | ARHGEF40 | CTB-12A17.2 | CTD-3148I10.15 | ANGPTL6 | CXCL6 |

### Table S1. Significantly differentially expressed genes (EZH2 Signature High vs Signature Low, p<0.05) continued.

| | | | | | | | | | | |
|---|---|---|---|---|---|---|---|---|---|---|
| CCL20 | HEMK1 | SPTLC1 | SMARCAD1 | DCUN1D1 | CTD-3128G10.7 | GPA33 | SLC22A23 | MROH2A | INHBC | HNRNPA1 |
| HIPK4 | ATAT1 | ALKBH6 | TMCO4 | CD109 | TRBV5-6 | AL928761.1 | CTF1 | RP5-1172A22.1 | OTUD7B | ZNF514 |
| RPTN | FBLN2 | GM1976 | KCTD2 | CSTA1 | LCK | TNFAIP8L1 | RP11-732A19.1 | ARL9 | SNX7 | APBA3 |
| ASF1B | GALNT7 | INSC | PPP1R7 | ACTA2 | IFFO1 | RGS9 | RAP1GAP | UPF1 | NBL1 | LRRC30 |
| USP45 | RPS4X | DDX5 | AKIRIN2 | ZDHHC21 | CTD-2521M24.4 | CLIC6 | TMPRSS11D | TMPRSS13 | PRKRA | ZNF304 |
| POLN | H2-EB1 | PDF | PIGU | CALM1 | OR2I1P | SPOCD1 | C1orf109 | C16orf70 | GSDMA | ARRDC4 |
| 43348 | HEATR1 | MTMR2 | USP5 | CGGBP1 | CXCR5 | REM1 | FAM83C | RP1-102E24.10 | TECPR1 | IL12RB1 |
| LIG1 | ERN2 | LCE1B | LIN9 | ZXDA | IGHV3-30 | CAPNS2 | RP11-61F12.1 | TRIM61 | ARF4 | SFRP2 |
| MS4A10 | TMC8 | NDUFA4 | GP5 | PLEKHS1 | BTN2A2 | PMCH | RCAN2 | RP11-121E16.1 | SLC26A7 | SP9 |
| E2F2 | PABPC4 | GNRH1 | SMAGP | EIF1AX | LILRB2 | PPM1H | ZEB1 | LINC01451 | P2RX6 | TRYX3 |
| BBOX1 | GDI1 | CIZ1 | EXOSC2 | MVD | TNFRSF25 | C1orf147 | PTPRH | RP1-193H18.2 | TUBGCP4 | TUBA1B |
| SLC25A24 | ATP13A2 | ZNHIT2 | REV1 | RC3H2 | FTH1P22 | FAM179A | CXCL16 | ZBTB45 | FKBP10 | ZGPAT |
| NGEF | DOK4 | UTP20 | TCEAL8 | PRR13 | SERPING1 | SLC46A1 | S100A13 | SRSF12 | FFAR2 | CROCCL1 |
| GM20554 | AK4 | TTI1 | EXOSC6 | EIF4E | TMEM173 | RP11-686D22.7 | DACT1 | MOCS3 | POLR3D | IFIT3 |
| KDM1B | DDX50 | DIRAS2 | FAM149B | FNDC1 | CTA-384D8.35 | PHACTR1 | HS3ST3A1 | RP11-493L12.5 | CD47 | C12ORF71 |
| RPL28 | 6430573F11RIK | MSTO1 | PITX1 | RAB8B | EMP3 | LCAT | RP11-373D23.3 | ASAP1 | TROVE2 | FSCN2 |
| PDLIM7 | MSH6 | TPT1 | TCF7L2 | AXL | SH3KBP1 | NACAD | SCARA5 | CCDC149 | B4GALT4 | PCNXL3 |
| RP2H | EDEM1 | DCAF15 | SNX9 | MAPK6 | SKAP1 | ST5 | C5AR1 | RP11-469J4.3 | SOCS7 | CYBRD1 |
| ING1 | BC053749 | GALNT4 | PTCD3 | CD2AP | IGKV1-9 | HOXB5 | EIF2B4 | CTC-559E9.8 | PDE3B | HDGFRP3 |
| GUK1 | UBOX5 | SNHG6 | MAP2K2 | ADCY6 | C1S | GPR15 | IQGAP2 | ZNF3 | ANKDD1B | LECT1 |
| SLC38A7 | 2200002J24RIK | HSP90AA1 | TENM4 | BOK | S100A4 | LINC00900 | FAM168B | AC009299.5 | PLA2G15 | MECR |
| ISOC1 | KCNK6 | NDUFC1 | BTBD19 | SEMA3D | SLC16A3 | IGLV2-34 | RP11-573G6.6 | MED23 | LYPLA1 | NCRNA00115 |
| BTBD2 | AFMID | WWC2 | TSPAN33 | MFAP3L | IL12B | CDC25B | EFNB1 | FOXS1 | ZIC5 | PDIA3P |
| ICA1 | NDUFA6 | SHC1 | UEVLD | PGD | IGHV4-61 | CMTR1 | RP11-756G20.1 | CLDN16 | VWDE | RAB4A |
| NPR1 | GNB1 | ETL4 | ARL6IP6 | OLFM4 | KCNA3 | FAM212A | SUSD2 | RP11-212I21.4 | LAMB3 | ZMYND17 |
| FOCAD | UQCR10 | SENP8 | SPRR1A | RAB2A | MLKL | IGHV3-72 | PRICKLE1 | SVEP1 | CFB | ST3GAL2 |
| NOP2 | DSE | CCNG2 | 43352 | ATG9B | CCL17 | RP9P | C9orf91 | PCDH8 | TPRN | IGF2R |
| PIM1 | DGKA | CRTC2 | ZBTB18 | C3 | IGHV2-26 | IFITM2 | PSMA8 | DUSP18 | PARP3 | OR2T8 |
| ALDH6A1 | VAT1 | SREBF2 | BCAM | TTC9C | RARG | SLFN14 | CCDC69 | FGF9 | DLG2 | RPS23 |
| HIST1H2BJ | GM11545 | SLC25A12 | LZTFL1 | PGM2L1 | BIRC3 | IGLV1-50 | INTS5 | SLC30A5 | DPM1 | LOC643008 |
| HSD17B11 | KIF15 | 9230110C19RIK | RHOD | CDCA7 | GVINP1 | EPHA1-AS1 | HMSD | LRRC41 | EMC8 | SCOC |
| MCTP2 | PICALM | ZKSCAN14 | CD44 | FSCN1 | ITGAM | FPR1 | RP11-641D5.2 | SOAT2 | TNFRSF10C | ANO9 |
| GTPBP4 | RNF183 | ARMCX5 | DOCK4 | PFN2 | RP11-81H14.2 | LLNLR-470E3.1 | TGFBR2 | SLC33A1 | LIMA1 | PDE6A |
| FOXI1 | RPL39 | CSTF2T | DDX19A | 5430419D17RIK | ARHGAP22 | SENCR | C5orf46 | CASS4 | IFRG15 | INPP5B |
| ZGPAT | GM7120 | TMX1 | ZMIZ1 | ZFP52 | IGKV1-5 | C10orf55 | PALD1 | PLAT | LSM11 | ATF7 |
| MYC | C1GALT1 | FAM69A | PRIMPOL | VPS54 | TRAV12-2 | SMIM10 | RP11-494M8.4 | COL13A1 | CLEC11A | C14ORF73 |
| LIG3 | RIPK4 | CLN5 | ANKRD40 | TMEM55A | WFDC21P | IGHV3-64 | ATP10D | AMPD1 | IMPAD1 | CCNE1 |
| 0610031J06RIK | CNPY2 | TPGS2 | XPO6 | DUSP6 | IGHV3-53 | RPL4 | BHLHE40-AS1 | RPL18P13 | SLC22A4 | CD2BP2 |
| HDAC7 | RALGPS2 | ACAA2 | IRF2BP1 | MED21 | HLA-DPA1 | PRKY | PCOLCE-AS1 | GJB1 | ADAM33 | DZIP1 |
| BMP7 | ADH5 | BTBD7 | MIPEP | FAM120A | IL34 | MB21D2 | AC017002.1 | RP11-302I18.1 | TMEM132D | FAM156A |
| TNFAIP2 | ZFP397 | AGR2 | EDEM2 | MOB1B | AL928768.3 | ABCB4 | RP11-85B7.2 | RP3-486I3.4 | MDH1 | LCAT |
| WDR26 | RPS7 | RUFY1 | CDKL2 | NFE2L3 | ADAM19 | AL122127.25 | SYT12 | PLCH1 | NKX2-2 | LOC100130581 |
| SEMA4C | NPLOC4 | PARP4 | LDLR | 1700066M21RIK | IGHV1-18 | SOCS1 | CFB | GDF3 | CNGB3 | NCAM1 |
| MXD4 | OXR1 | BORA | CBX4 | ZBTB7B | ARL4C | EIF4B | RP11-326C3.7 | ZFP92 | C15orf54 | OSBPL6 |
| ASAH2 | EIF4G2 | ANKRD27 | VOPP1 | SLAIN2 | IGHV3-49 | FBLN2 | MXRA7 | ATP5G2 | LPHN1 | PLEKHH1 |
| 1810019D21RIK | EIF1 | CAV2 | BSN | CDKN1C | FASLG | CCDC3 | RP11-1070N10.3 | HRASLS2 | ZNF672 | RAB40A |
| AFAP1L1 | HVCN1 | TCTN3 | AURKA | DNM1L | TMIGD2 | RP11-405M12.4 | CAPN13 | ATG16L2 | RPAP2 | RAB43 |
| OAS1B | MTM1 | HEG1 | MKLN1 | RALA | CTD-2020K17.1 | UGT1A6 | RP11-452L6.6 | TIGD6 | INSR | RMST |
| HEXIM1 | SLC52A3 | ZFP759 | CASP3 | AP1S2 | MFNG | AIM1L | IGFBP6 | PPP1R14A | PTK2B | SMYD2 |
| IER5L | SAA1 | SYNGAP1 | RECQL | ZFP120 | OSCAR | NCR1 | RP11-9G1.3 | TM9SF3 | PDIA6 | TMIGD2 |
| ANTXR1 | HIST1H2BC | LMO1 | AHNAK | ENTPD8 | TREML2 | AL122127.2 | TRPC1 | AC009245.3 | CYP27A1 | ZNF33B |
| MMAA | TXNDC17 | GM15417 | GYS1 | LTF | RASGRP2 | TRPA1 | KCNE1 | LINC01152 | ENO2 | ZNF765 |
| EIF4G3 | HTRA1 | FAM63A | GPC1 | CUL3 | IGKV2-24 | ADAMTSL4 | CLIP2 | HDLBP | CYFIP2 | CHRNA1 |
| PTP4A1 | PPARD | SUV39H1 | YPEL1 | 1600012H06RIK | MNDA | ROBO1 | HEATR9 | NOS2 | BBIP1 | AKAP5 |
| SNHG7 | EDARADD | ZBTB38 | HIST2H2BE | ADAM10 | IGHV4-34 | TMSB4X | AGAP1 | SMARCD2 | SCIN | GSN |
| LDHA | RPL32 | LEPRE1 | PMF1 | SYPL | AC104820.2 | RP11-344B5.2 | ANO7P1 | RASGEF1A | RARRES3 | SH2D3C |
| HMGCLL1 | USP10 | RBM28 | ORAOV1 | PLAT | CD40 | CNRIP1 | ALDH6A1 | SEC22C | STEAP2 | PLEKHO1 |
| C030006K11RIK | RASA1 | FAM216A | CACNA1A | NSDHL | GZMH | TRO | GPRC6A | ICAM1 | TUBB1 | AOC2 |
| VASP | CHTF18 | BC005624 | DYNC1LI2 | H2-T23 | C1R | S100A5 | COL18A1 | FTH1 | INPP5A | UQCC |
| ADCK1 | 2810013P06RIK | D17WSU92E | STK24 | PDCD10 | IGHV3-7 | IGLC6 | DKK1 | RP11-107N15.1 | SLC45A4 | C19ORF45 |
| IFI204 | FBXO32 | ANKRD10 | ATMIN | LCORL | ALDH3B1 | KIAA0141 | CPXM1 | HIST1H2BN | RASGRP4 | ZNF471 |
| SLC23A3 | CLCN2 | COQ9 | TSPAN15 | IKZF5 | RP11-686D22.10 | MMP13 | GDPD3 | SYNC | TSPAN16 | CARD11 |
| PKP3 | CYR61 | IRAK1 | 1810034E14RIK | AP3S1 | LINC00494 | ZBED4 | CRIP2 | FGFR2 | PCDHA11 | LOC399744 |
| PRSS27 | ZFP420 | NRAS | TMEM109 | KRTDAP | CD1C | RP11-379F4.4 | TRPV3 | C1orf87 | SREBF1 | TRIM67 |
| NNT | CCDC104 | PRPS2 | TPR | MDN1 | IFI27L2 | INSRR | MXD4 | GSPT1 | TMEM135 | RNF40 |
| OSBPL10 | CCNDBP1 | BARD1 | A930004D18RIK | TMED1 | TRBV6-5 | CSDC2 | BTBD9 | TADA2B | C1QB | ARHGEF17 |
| PRMT6 | DPP8 | SCARB1 | EVI5L | KCTD14 | TSPAN32 | FGFBP1 | CDKN2B-AS1 | RP11-1149M10.2 | CFDP1 | BTBD17 |
| IFI47 | MPDU1 | ACO2 | UPK1B | C330027C09RIK | IGKV1-6 | LILRB3 | GGT6 | FKBP7 | MCPH1 | TBX21 |
| NOP10 | OSTM1 | FTX | IRAK4 | LSM12 | IGHV5-51 | KEL | RELB | MSRB3 | NINL | D4S234E |
| KRTAP1-5 | KDELR1 | ADCYAP1R1 | NCMAP | ZFP947 | RP11-693J15.5 | ADAMTS2 | RP4-706A16.3 | OLFM2 | LYSMD4 | CD48 |
| TEAD3 | DCUN1D4 | 4833418N02RIK | FAM83B | CCDC3 | ADA | SYN1 | ATXN7L3B | ITGBL1 | ARPP19 | HOXD9 |
| CSF1 | SRC | NEU3 | CBL | RHOA | GLIPR1 | S1PR2 | SFRP4 | LAP3 | LPPR3 | ZNF343 |
| PKN3 | LAD1 | DUSP11 | IGFBP7 | RNF145 | KLRD1 | RBM38 | TOM1L1 | GTPBP2 | MKNK2 | TREML3 |
| CORO2A | FA2H | FAM195A | NDUFS4 | ZFP942 | IRF8 | RP11-553L6.2 | IQCA1 | EVPLL | ZDHHC11B | ABCG2 |
| CEACAM1 | CAST | ENTHD2 | ZKSCAN8 | NUCKS1 | SLFN13 | B3GNT7 | RP11-567M16.1 | SFT2D1 | UBA7 | CCDC34 |
| FMOD | CYB5B | GSE1 | PARD6B | SHOC2 | NOD2 | TRBV21-1 | RUSC2 | ADAR | XYLT2 | GLRX5 |
| 1110008P14RIK | SLC12A6 | ALDH3A2 | IMPA2 | DPYSL3 | CAPG | PLK3 | APEX1 | FGF2 | C3 | SOX8 |
| E2F1 | FAM20A | ACSM1 | KCTD3 | UBA7 | AKNA | IGLVI-70 | LINC01260 | PNMT | GRN | C10ORF55 |
| RPS14 | RAB12 | FGFBP1 | CCDC17 | RAB14 | DRAM1 | CTC-251I16.1 | GRHPR | TNIP1 | DDX47 | CASP8 |
| STARD10 | CLN8 | POLE | USP40 | PVRL2 | IGHJ3 | IGHV6-1 | IRAK3 | RP11-534L20.4 | FAM181B | GPLD1 |
| LDHB | MAP1LC3B | CD63 | PIAS2 | FAM83E | CD274 | ELN | NFASC | AC016735.2 | TPPP | CCT4 |
| RPLP0 | ACYP1 | WDR83 | KRTCAP3 | OAS1B | APOL3 | RP11-652L8.4 | TBX2 | RP11-420L9.2 | TRPV1 | LOC100133957 |
| SMAD3 | HIST1H3C | OCIAD1 | WDR73 | HVCN1 | IGLV2-8 | RP11-472K17.3 | NTN4 | JAK2 | C1QA | NDC80 |
| WFDC8 | SLC39A4 | CENPA | DEFB6 | STAG2 | IGLV3-21 | TRBV30 | SFTPA1 | LRTOMT | FOLR3 | PHLDB2 |
| 2610528A11RIK | GRK4 | TBCD | KBTBD3 | ZFP281 | LINC01550 | CAPN11 | C9orf152 | RPS10P3 | PCSK1 | TAB3 |
| DDIT4L | SNRNP200 | PAK6 | HSF4 | GNPDA2 | TGFB1 | DUSP5 | NFATC4 | CAMSAP3 | TRMT10A | TCP11L1 |
| ROGDI | PTPLB | REPS1 | VANGL2 | CNOT6 | TRAV29DV5 | C19orf35 | TMEM5 | SEC23A | PTCHD4 | ARL6IP4 |
| CHDH | CDK5RAP3 | BBC3 | PDK3 | CUL4B | IGHGP | SYNPO | KANK2 | FAM81A | PLEKHN1 | KLHDC8A |
| EPHA2 | NOC3L | ECH1 | WBSCR22 | CAB39 | TNFAIP2 | CAMK4 | RPL7AP30 | ZBTB39 | FOXK1 | C1ORF229 |
| TOPORS | HEXA | FSCN1 | FHOD3 | IL20RB | CD69 | AC116366.6 | NLRP6 | AP000439.3 | ZBTB37 | DNMT3B |
| MRPS34 | LCN2 | SNORD12 | PTBP2 | DSP | IGHV3-33 | TPPP3 | NPM1 | MRVI1 | MANEAL | GIPC1 |
| SERPINA3I | LIPT2 | DDHD1 | ALOX5 | XPO1 | AC133644.2 | IGKV1D-42 | RP11-395G23.3 | EOMES | S100A14 | RPL34 |
| MID2 | NEK8 | CEBPZ | OTUD5 | SCARA3 | RP11-693N9.2 | RAB11A | FZD2 | RP11-366L20.2 | ANKMY2 | TTF2 |
| OSGIN1 | RNFT1 | ORC6 | LIPH | G2E3 | FCGR3A | AC103563.2 | PCOLCE2 | C2CD4D | CTXN2 | ANKRD20A3 |
| MUC2 | TRIM59 | TFE3 | PSMG1 | LONRF3 | IFIH1 | PIWIL4 | RPS4XP12 | PDE1C | TMTC4 | ASB13 |
| HOXA11OS | CYB5RL | PELI1 | GNAS | PLXDC2 | IGLV3-10 | LEFTY1 | IGLJ2 | EEF1A1P12 | PNMT | GPR4 |
| UBD | KRT15 | WNT11 | ZFP873 | RPS17 | CEACAM21 | IRF5 | CD2AP | COL22A1 | CSF1 | PIM1 |
| PARVA | CCDC79 | OAF | INO80 | SLC2A12 | CD33 | NGF | TMEM255B | CLDN18 | RIC3 | SNX27 |
| ASPA | PARL | CDK10 | MAVS | OSBPL8 | CXCR2P1 | PDGFRB | TTC3 | MARS2 | FAM71F2 | LOC100131193 |
| SLC16A6 | PHACTR2 | CPNE7 | SPATA5 | TSPAN4 | GPR68 | IGHV1-2 | LILRA2 | AC010136.2 | HOXB7 | RASGRP3 |

#### Table S1. Significantly differentially expressed genes (EZH2 Signature High vs Signature Low, p<0.05) continued.

| | | | | | | | | | | |
|---|---|---|---|---|---|---|---|---|---|---|
| UIMC1 | MAPK6 | GBP4 | PPAN | KCNE4 | APOL6 | MAML3 | ADORA2B | AFAP1L2 | STX6 | FAM162B |
| FERMT1 | GNPTG | CCDC3 | MIS18A | TTLL4 | TRAV4 | MDS2 | AHCY | TEX41 | KCNQ4 | OR9A2 |
| SENP1 | SLC7A3 | EI24 | PRMT10 | NIPAL1 | FAM129C | IGKV1D-12 | FGF18 | LA16c-390H2.4 | ATG5 | C1ORF177 |
| ARHGEF40 | CDNF | RAB34 | UBALD1 | ARF4 | MMP14 | TRPC6 | MOAP1 | SAMD11 | CTSD | NKX2-1 |
| SLC10A3 | CASQ2 | INIP | DNM3 | GM4944 | HLA-DRA | DMC1 | PMAIP1 | AC092171.4 | ZFP69B | ADAM30 |
| SEMA3D | TRMT12 | CALML3 | SHCBP1 | FAM126B | TRAV3 | ANXA8 | SLC39A9 | YIPF3 | GTF3C4 | B3GALT4 |
| PDGFB | HAVCR2 | HUNK | DOCK1 | DUSP1 | KIAA0226L | PCP4L1 | AUTS2 | PRKG1 | CHGB | RAPGEFL1 |
| EPHA7 | PHC3 | SORT1 | LAMTOR2 | TMEM30B | IGJ | PDE4A | SPARCL1 | RNU4-62P | C2orf71 | RPS12 |
| HELZ2 | ETV5 | EIF3E | NKPD1 | CTNND2 | RP11-325F22.2 | OAF | TNNI2 | MYBPC3 | SLC25A16 | USP53 |
| 1500012F01RIK | CDCA2 | NOL6 | IPP | KRT6B | IGHV3-23 | IFITM4P | FLJ27354 | AC093673.5 | FBXO17 | BOC |
| ATXN2L | ECHDC2 | SLC25A5 | PCCB | PRSS32 | TMEM176A | NLE1 | MCEE | C16orf58 | CD63 | LOC151300 |
| HBEGF | PDIA6 | SDF2L1 | SMN1 | THBS2 | CSF1R | FMO3 | RP4-794I6.4 | BAG6 | ACP5 | RNASEL |
| SLC9A5 | FAM117A | E030030I06RIK | SYT16 | TTC14 | PSMB10 | SYNDIG1 | ZBTB7B | SIGLEC12 | SDC4 | SRCIN1 |
| DCXR | ACOT1 | TMEM71 | ETS1 | BCAM | TRBV7-3 | RP11-733O18.1 | CD300LB | C20orf85 | VMO1 | HS1BP3 |
| RPS29 | DUS3L | KLK10 | GALNT1 | LGR5 | TRANK1 | BATF3 | ZNF385D | ELAVL4 | PAPPA2 | LPAL2 |
| EPN2 | NUP188 | ZGRF1 | VARS | SPAST | IGLV3-25 | FER1L4 | CCL8 | ZNF749 | YTHDF3 | ZBTB1 |
| BC004004 | GPLD1 | PARN | MRPS18C | DYNC1I2 | SLC43A3 | TGM1 | CLEC4C | TMED2 | HOXB6 | AAGAB |
| 1700037C18RIK | SDHA | 1110038F14RIK | PPIC | YY1 | FCGR2B | CRABP2 | ENOX1 | CLDN1 | RGS18 | GEMIN8 |
| ALG3 | TIMM10 | MRPL10 | FAM89B | PLAC8 | HLA-DRB6 | HOPX | C11orf16 | RP11-370F5.4 | TPI1 | TTC5 |
| PCGF3 | GM20605 | VPS26A | RAB39B | TMEM167 | NINJ2 | PRTFDC1 | ACTRT3 | PATZ1 | MTDH | WBP2 |
| SETD7 | CYTH2 | TRIB3 | 1110001J03RIK | SLC2A4 | ARNTL2 | RP11-875O11.1 | ATPAF1 | HINT1 | HLA-DQB1 | ANXA7 |
| GNPNAT1 | TCAIM | CHN2 | MED12 | ADSS | AC006369.2 | IL18R1 | RAB3B | DCTN5 | NBPF12 | HNF4G |
| LAMB3 | VEGFB | LPCAT4 | AHSA1 | ADH7 | TRBV10-3 | RP11-876N24.4 | GNPNAT1 | ACLY | RABGAP1L | OR52R1 |
| CPNE2 | SRCIN1 | NOMO1 | COA7 | SPPL2A | RP11-327F22.2 | RP11-564A8.8 | LTBP1 | TRPC4 | C11orf70 | FLT4 |
| CEP55 | RBFA | MAST3 | CD34 | GCH1 | NMI | GJA1 | RPL12P4 | AGPAT9 | ZNF648 | NEURL1B |
| BNIP2 | DDX3Y | MTPN | TEAD1 | GSR | SIRPB2 | AC009948.5 | RP11-13P5.1 | KIF19 | CDC73 | PRIC285 |
| GNPDA2 | POP1 | SRSF10 | GIT2 | ATP6V1G1 | IGHV2-5 | KLF8 | YWHAE | RPL10AP6 | ALDH1A3 | TMX4 |
| NEAT1 | ERLIN1 | AI506816 | SFXN3 | COL4A2 | TLR2 | WWC1 | AC103563.8 | SLC46A2 | ENTPD3 | LOC441046 |
| 4930427A07RIK | LRIG1 | TAF4A | PCDHB16 | POLG | MST1R | C15orf61 | CERCAM | BCAS4 | DNAH2 | FURIN |
| PALB2 | WDR12 | NME1 | ZFP629 | KPNA1 | JAKMIP2 | RP11-161M6.3 | RPS2 | CA11 | LAMB2 | GP6 |
| GPS2 | RASGEF1C | CCDC115 | FAM83G | SSB | NPL | COL6A1 | SGIP1 | GPR162 | PLS3 | S1PR3 |
| CBX6 | RPL10A | ZAK | ABRACL | TMEM171 | RP11-455F5.5 | SRRM3 | ZNF615 | SLC46A3 | SLC26A8 | SLC26A11 |
| XDH | SURF2 | HARS | NR3C2 | ATXN3 | APOBR | TLR7 | CLCA2 | PKHD1L1 | GARS | ST8SIA5 |
| FBP2 | GPC2 | SULF2 | PQLC2 | MBTD1 | IGHV3-48 | SECTM1 | IGLV3-13 | USP2 | ITPKA | KIAA0802 |
| FAS | BCS1L | SRRT | BCAR3 | GSDMD | KIF21B | C2orf54 | RP11-536O18.1 | PTHLH | RALGDS | C5ORF54 |
| FZD4 | IL18RAP | TACC2 | GPX4 | SEH1L | TRAV8-4 | FBLN1 | KRT13 | VAPB | PKD1L1 | MOXD1 |
| EIF2AK1 | NUP88 | ZFP865 | ANKRD52 | MATR3 | CD84 | BMPR1B | RAB12 | CAAP1 | HPS3 | SSTR1 |
| TGIF2 | B4GALT6 | LACC1 | B130024G19RIK | YES1 | HSPA7 | ATP8B3 | KRT42P | CTC-327F10.5 | C3orf49 | UCP3 |
| TBX3OS2 | FPGS | CCDC68 | CC2D1A | UBC | OR52N4 | RP11-542M13.3 | FOXF1 | CMTM5 | CNP | BFSP1 |
| AOX1 | EPN3 | HINFP | ESYT1 | UBE2V2 | ADAMDEC1 | UNC5B-AS1 | ADAMTS8 | IVD | HAMP | HSPB8 |
| EPHB2 | RAB21 | DDX49 | NUDT17 | SPRR2J-PS | C3 | IGHV3-62 | PRSS50 | SAMM50 | ZZZ3 | KCNK10 |
| ATP13A3 | BHLHE40 | TRIM8 | MTHFD1 | C1GALT1 | MIR8071-2 | GPX2 | TBC1D2 | UBE2R2 | WDFY1 | RCCD1 |
| CPM | ZFP827 | CDK5RAP1 | PID1 | CXCL3 | IGHV3OR16-13 | EPB41L3 | DNAH11 | FES | PKNOX1 | TMEM22 |
| RPS23 | ZHX3 | FEM1A | RAC3 | CYB5B | IGHD | ADSL | FASTKD3 | HLA-L | SYN3 | CFI |
| WDR89 | CHD2 | ARHGEF1 | TPP2 | CHAC2 | RP11-1399P15.1 | CYGB | SVIP | SNHG24 | MAGEA6 | LOC400696 |
| NISCH | CALM4 | KLHDC7A | ATP6AP1 | YIPF5 | CSTA | LRRN3 | RPL7AP66 | RPL18 | GNB3 | RPL22 |
| BARX2 | NUDT16 | SKA1 | SPICE1 | RNF138 | UNC13D | SH2D1A | PAMR1 | MAP3K14-AS1 | BCR | FXYD6 |
| EMC9 | SLC25A20 | CTTN | SDHD | KDM1B | CD70 | TTLL12 | LILRA6 | ADORA1 | DNM1 | PLEKHG5 |
| ANKRD12 | CYP3A13 | HIST1H2AK | ASAP1 | ARL15 | TRBV4-1 | PADI3 | SMIM14 | EHD4-AS1 | NIPAL2 | PRICKLE4 |
| GAPDH | TRIB1 | KIF9 | IGF2BP1 | GM13157 | GIMAP7 | CECR1 | EFHD2 | SIPA1L3 | ECT2L | TRAM2 |
| PLA2G16 | TTC14 | ATG101 | BRPF1 | SETD8 | RP11-134L10.1 | TRMT5 | BDKRB2 | AC000068.5 | PIEZO1 | ESAM |
| RPS12 | CLIC5 | AP1B1 | NHLRC4 | FOXC1 | TNIP3 | DHX58 | SLC4A11 | HOXB8 | MAGEA12 | TIGD2 |
| TMEM158 | ZFP672 | CYP4F13 | GM128 | RPS6KB1 | LINC00152 | RP11-166D19.1 | PTPRO | LASP1 | LY6K | FCHO1 |
| RBP1 | SLC4A3 | SH3PXD2B | EPPK1 | PHF6 | RP11-47L3.1 | COL21A1 | RP11-297B17.3 | SGCA | YIF1B | C19ORF39 |
| USP43 | TRIM14 | FAM25C | EOGT | RIN1 | RP11-23J18.1 | SPOCK1 | RP11-323C15.2 | POLR3E | KATNBL1 | GPR172B |
| PUS7L | IFI30 | ELMSAN1 | HIST1H2BM | DYNLRB1 | GPR84 | TEPP | MEF2C | RP5-1148A21.3 | LEPREL4 | MAP3K8 |
| NIT1 | CDC42EP1 | FAM173A | AKR1C14 | CACYBP | PIK3R6 | PRRX2 | NFKBIE | KIAA0195 | PSMA6 | ATP5G2 |
| ZKSCAN17 | FAM53C | CHCHD6 | MAP4K2 | PCGF3 | TRAV26-1 | BICD1 | TBC1D1 | C11orf87 | FAM43A | CDH22 |
| ABO | HPSE | TINF2 | HCFC2 | JAG2 | TRIM34 | APLF | KCNJ11 | TMED10 | GPR173 | CDRT1 |
| CCNA2 | NAV1 | KCNJ4 | TRAM1 | BID | GAL3ST4 | WNT1 | FNBP1L | SLC24A3 | TMEM192 | DNAH14 |
| GTF2E1 | DDX47 | RLIM | MINOS1 | TMEM64 | TRAV16 | C11orf88 | IGHV3-76 | CASD1 | TIMP2 | MDFI |
| PRX | CMTM4 | PIGA | POLR2H | PIR | CLIC2 | TRABD2B | AC092066.1 | ITIH1 | NLRP13 | PCLO |
| TSHZ2 | UBE2L3 | SYT3 | TOE1 | REPS2 | ZNF683 | RNF175 | MYEF2 | PRNP | BDH2 | RPL35A |
| TUG1 | SLC20A2 | SLC17A9 | TAF11 | WDR4 | CD1E | HOXD10 | RNY3P8 | PITPNM1 | TMPRSS11F | SHQ1 |
| PXDC1 | GLT25D1 | RPL9 | D3ERTD751E | VNN1 | MVP | CLEC4F | NEDD4L | CUL7 | MTRNR2L6 | TTLL4 |
| RPL3 | SDAD1 | CRAMP1L | INPP5J | USP14 | ABI3 | RP4-610C12.3 | IL4R | IGHV5-78 | KCTD20 | USO1 |
| RASSF7 | PIGH | KPTN | RRP1 | SLC4A11 | TRBV12-3 | C1QTNF2 | COTL1 | SNCG | C6orf10 | FAM194A |
| VSTM5 | ZFP28 | MARCKSL1 | PTGR2 | RAB25 | IER5 | GIMAP6 | CTB-193M12.4 | ACTA2 | ADCY10 | IMPAD1 |
| 1700037H04RIK | PRAP1 | LASP1 | ATP5S | APBB3 | IGLV9-49 | LINC01187 | TSKS | DNASE2 | MNDA | C13ORF29 |
| TACC3 | BRMS1 | SLC35B2 | ABHD5 | B4GALT5 | IGHV1-46 | AC023590.1 | SH3RF1 | PALLD | ZNF221 | CATSPER2 |
| RPS3 | IL11RA1 | EXO5 | 1700020I14RIK | CLIC3 | AGAP2 | NOA1 | GPRC5B | ADGB | FAM107A | NBPF7 |
| GLB1L | DMPK | ZFP324 | KIF14 | ADRB2 | RP5-1171I10.5 | RP11-689B22.2 | RP11-320N7.2 | RP11-93B14.10 | AC112205.1 | NID1 |
| VASH2 | SPATA24 | TMEM134 | SEMA4B | IMPAD1 | AC092580.4 | DOK4 | LHFPL2 | BTG3 | GLT8D2 | NR5A2 |
| ANLN | SENP2 | PXN | TMC3 | PTK7 | STAT5A | RPL5 | RPL6P27 | DPYSL3 | MAGOHB | RFX6 |
| CLDN12 | ITPK1 | ZFP933 | SLC30A9 | FUBP1 | NCF2 | GALNT8 | CD209 | COL4A1 | RAB3C | S100A14 |
| PCCA | MYL6B | DIAP3 | AZI2 | SOX7 | RP11-290F5.1 | S100A14 | ESR2 | FAR2 | AC027763.2 | ST6GALNAC2 |
| LCE6A | PIM3 | BRD4 | ZFP408 | TMEM45A | TRAV12-3 | HIC1 | MBOAT2 | SULT1E1 | NSMF | TXN2 |
| AKAP1 | NIP7 | WDR43 | KRR1 | C1D | HPSE | AC017074.2 | ACVRL1 | BCL2L2 | IL6R | MCCD1 |
| PLXNB2 | PITPNM1 | PJA1 | NMNAT1 | ARC | IGHG1 | ERGIC1 | SWAP70 | MTRR | CACNG6 | SLC28A2 |
| ZFP365 | HNRNPD | EPS15L1 | CEMIP | CNEP1R1 | IGHV3-66 | STK38 | GRM2 | RP11-61L23.2 | RNF187 | AOAH |
| ANK | PABPC1L | BRF1 | CLSPN | OIT1 | SERPINB1 | POM121L9P | CAND2 | HTR1F | LHX3 | FLJ37201 |
| LRRC3 | FUT9 | LRRC16A | COL4A3BP | BCL2 | CALHM2 | SULF1 | RP11-309L24.6 | AHCYL2 | KRT23 | LOC100132707 |
| ROBO3 | RPL19 | TMEM14C | NUMBL | H3F3A | AC147651.4 | RP11-44K6.4 | CPEB1 | AVL9 | CTSL | RGMA |
| RPS18 | 4732491K20RIK | PHF3 | PPP1R9B | EEA1 | TIMP1 | CASKIN1 | TRBV11-3 | CLIP4 | POLR2C | ZNF229 |
| SLC23A2 | PHKA2 | HPDL | NUDCD3 | KCNQ1 | CCR1 | CNPY4 | NAV2 | OGN | AASDH | ANKRD6 |
| ITGA6 | TLR3 | TTC19 | H2-M3 | CABLES1 | TRBV9 | GJB4 | PZP | RABL3 | CCDC83 | DYNC1LI1 |
| SUOX | FGF15 | D8ERTD82E | SPOP | ROCK2 | SLCO3A1 | AP000662.4 | CLEC12B | EHD1 | LGMN | FNBP1 |
| SLC6A14 | DZIP3 | NPM1 | SPATA5L1 | IPO4 | AC009950.2 | FBLN5 | SRP72 | RPL31 | GALNT7 | HOXA3 |
| ASAP2 | NDUFB6 | 2700046G09RIK | FBXO25 | B230219D22RIK | STX11 | CTD-2566J3.1 | LINC01426 | RP11-576I22.2 | FAXDC2 | TMCO7 |
| ANG | KRT6B | SELENBP1 | MAP2 | CSNK2A1 | GNGT2 | CEBPZOS | BEST4 | RPL37A | PTP4A3 | PCOLCE2 |
| TNF | CYP2F2 | CD276 | RARRES1 | TNFRSF11B | ELF4 | PIP4K2A | PRR36 | EIF3E | SLC34A3 | CD24 |
| FCGBP | METTL13 | SLC22A17 | FIGF | FGFR3 | CMAHP | CTD-2207O23.3 | EGLN3 | CRTC1 | C1QC | DAPL1 |
| YBX3 | RBL2 | FASN | MYD88 | VCAM1 | IGKV1-39 | RP5-899E9.1 | ARHGEF1 | RP11-380J14.1 | NOSTRIN | FSTL3 |
| MTA3 | TRAPPC9 | SLC5A6 | SPRY2 | DLL1 | LYL1 | GAS6-AS2 | RN7SL689P | PPM1E | PIGT | IRF6 |
| SLC25A44 | SLC16A11 | BEST2 | SETD3 | TGFB2 | FLT3 | PDPN | DAK | MRPS14 | PCBP3 | TMEM49 |
| CPSF1 | GTF3A | RDH5 | DNMT1 | ST3GAL4 | GZMB | RP11-291B21.2 | TTC38 | RPL29 | GPR64 | FAIM2 |
| MEA1 | RFX1 | ATOX1 | NCAPH2 | EPG5 | CLEC2D | TRBJ2-1 | ACACB | MIR27B | FAM27D1 | C2ORF62 |
| WDR35 | MYO5B | TCEANC2 | ZFP651 | RPS26 | CLECL1 | LINC00525 | SLC28A3 | FGF11 | DYNC1LI2 | LOC145474 |
| SLC50A1 | CCDC58 | GM2373 | MRPS9 | RPA2 | CYBB | PTRHD1 | FAN1 | KLRC4 | UCHL5 | JRKL |
| HOXA2 | MRPL34 | PDZK1IP1 | CDHR1 | ASNSD1 | SLCO2B1 | SIRPG-AS1 | DLEU7-AS1 | ERVV-1 | KDELR3 | DSCR9 |

#### Table S1. Significantly differentially expressed genes (EZH2 Signature High vs Signature Low, p<0.05) continued.

| | | | | | | | | | | |
|---|---|---|---|---|---|---|---|---|---|---|
| SMAD7 | FBLN1 | CTTNBP2 | FOXN2 | HOXD13 | MPEG1 | UNC5B | RP11-73M7.6 | FAM21A | MYOF | TGIF2 |
| CTSO | 1110051M20RIK | 1700067K01RIK | SOCS6 | EIF1 | IGHV4-28 | TRUB2 | CTC-455F18.1 | ISM1 | CBR1 | CFDP1 |
| SLC7A8 | PLEKHS1 | DMWD | PDIA4 | APOL9B | CD300LF | AC004067.5 | RP11-389C8.2 | QSOX1 | SULT1C2 | CMTM2 |
| UGT1A1 | FAM160B2 | LACE1 | NAGPA | PIAS2 | IGHV4-59 | LY6D | AIFM2 | C12orf42 | PSME2 | STARD4 |
| PGF | CUEDC2 | MCEE | ATP5G3 | UBE2B | TRAV9-2 | LILRA1 | ANXA2P2 | LLNLF-187D8.1 | GAPDH | TWSG1 |
| RHBDD1 | ZHX2 | TAF6L | P4HA1 | SCNN1A | C8orf31 | ATP6V0D2 | IGHV3OR16-15 | H2AFJ | HCN2 | NCRNA00167 |
| CYP2S1 | NR2C2 | PDRG1 | SDHC | GSTA3 | PCED1B | COL3A1 | RPL4P5 | PML | PSMA7 | PIWIL4 |
| EMP3 | IDS | 1110020A21RIK | C2CD4A | OAS1G | PDE6G | HELZ2 | IGKV2-28 | AC007879.7 | GPD1 | RIMS3 |
| H2-KE6 | ECHDC3 | HOXD13 | SLC27A1 | CDC42 | GBP6 | CTD-2186M15.3 | IGKV1D-27 | RP5-1091N2.9 | UBXN4 | UGT1A9 |
| RAB26OS | KTI12 | SLC39A6 | ARID1A | ARHGAP26 | IGHV1-24 | G0S2 | RP11-848P1.3 | SMAD3 | SNCA | ZNF813 |
| NDRG3 | DCTPP1 | PLA1A | SLC27A4 | RYBP | IGHV4-4 | TMIE | HSD17B2 | RP4-673D20.3 | FGF18 | CXCL13 |
| 1600014C10RIK | C130046K22RIK | ZADH2 | GSAP | TBL1XR1 | IL12A | RPL6 | SLC6A20 | RPAP1 | ISM1 | MPP5 |
| RPS6KA3 | TRIM13 | SLC33A1 | UBE4A | 4833423E24RIK | SFRP2 | RP11-489O18.1 | TRAV10 | RP11-327J17.9 | CASKIN1 | TMEM56 |
| DTX4 | ZFP329 | BOD1L | | TBC1D8B | IGLC3 | CYP4B1 | RP11-253M7.1 | PLEKHA3P1 | PSD3 | NCAM2 |
| TNFRSF18 | GNS | KLRG2 | | TC2N | CFP | SMCO2 | MEF2C-AS1 | EXD2 | GEMIN5 | MID2 |
| LRRC75A | MICALL1 | MBNL3 | | PGM1 | FYN | IMPDH2 | LEFTY2 | RP11-160O5.1 | CD207 | NIPSNAP3B |
| SMIM3 | ARL2BP | FAM214A | | CASP4 | BATF | THBS2 | OSBPL10 | RP11-667F14.1 | HSPB2 | ASAP2 |
| KLHL5 | UBE2S | SCAP | | CAPZA1 | RFTN1 | PHYHIP | F2RL2 | IGKV1D-33 | FBXO41 | C10ORF122 |
| RANBP17 | PINX1 | D16ERTD472E | | SMEK1 | GIMAP1 | RPL32P1 | SORCS1 | RPS14 | METTL16 | IGFBP4 |
| EGR1 | SLC19A1 | ZFP935 | | ING3 | TRAV21 | ABCB1 | FAM13C | NPRL3 | PRSS12 | NCAPD3 |
| NDUFA1 | PKP2 | DPY19L1 | | ANKRD1 | GALNT6 | AC246787.1 | HOXB3 | ZNF804A | GPR89B | FLJ45244 |
| SREBF1 | ATP8A1 | TARSL2 | | HCAR2 | CXorf65 | ZNF614 | AMOTL2 | MIR4537 | GUCY1B3 | ESPNP |
| ALDH1A7 | ZC3HAV1L | TMBIM4 | | SMNDC1 | EMR1 | TAX1BP3 | CDK18 | EXT1 | PAPOLB | PARP14 |
| MPP1 | MBD1 | TMED10 | | RASD1 | CXCL13 | SSTR2 | OVCH2 | DIO3 | CD9 | PDE4D |
| PIK3IP1 | GM166 | PTPLAD1 | | TOMM5 | LINC00239 | ACSM5 | RP11-274E7.2 | LCNL1 | C1R | PPP1R3G |
| FETUB | ALG1 | PPA2 | | TMEM168 | RELT | CHAD | TXK | LINC00540 | SEMA3E | AIF1L |
| SOX15 | RPL7A | D10JHU81E | | PTPRS | RP5-887A10.1 | PLCD3 | RP11-728F11.4 | GBGT1 | PELI3 | C14ORF23 |
| PDZD11 | GM13826 | IFITM10 | | CTU2 | IGHG2 | RP3-416H24.1 | TWF2 | ARHGAP27 | WFDC1 | ZNF835 |
| FMO3 | OAS1C | CMSS1 | | ST6GALNAC2 | TRAF1 | MATN2 | CTB-60B18.18 | FMO1 | CILP2 | PTCHD3 |
| ODF2 | PDP1 | POMT2 | | HK2 | SIGLEC7 | RBP5 | BMP1 | RP11-7F17.3 | TMED1 | MEST |
| RPL37A | BYSL | LRRC28 | | NUSAP1 | RAB37 | CTC-479C5.12 | RP3-449M8.9 | RMND1 | NBPF11 | ANK3 |
| ATP9A | RPP14 | PLD2 | | MBTPS2 | GPR183 | CXCL1 | IGKV1D-13 | RP11-627K11.1 | KIAA0368 | KLHL24 |
| TMBIM6 | NOP9 | RNF34 | | STAT2 | CASP1P2 | TMLHE | RP11-170M17.2 | RASEF | SUV39H2 | MOS |
| PLXNB1 | ZCWPW1 | RNF126 | | MAX | RP11-428G5.5 | RP4-570O12.3 | RP11-327F22.1 | FXYD2 | PSG1 | DUOX2 |
| GSTM1 | MRPS24 | ACBD5 | | S100A7A | DOK3 | MXRA5 | RP3-455J7.4 | BCL2L10 | AF131215.5 | GIMAP8 |
| FBXO33 | GUCA1A | ANO10 | | AREG | GGTA1P | IL20RB | GALNT7 | OLA1 | CHN2 | PRSS23 |
| SPAG5 | CCDC32 | HEY1 | | PARP10 | PIK3R5 | ZFP36L2 | PDLIM2 | AHR | INHBA | FAM66C |
| KIF5B | EMC1 | BLVRA | | LYPD3 | FMNL3 | CCL4 | NUDCD2 | AC074289.1 | ZNF283 | ROBO3 |
| H2AFJ | SSR3 | 1700034F02RIK | | ZFP36 | CFH | RP11-347C12.3 | RP11-1334A24.5 | AC004870.4 | PRSS37 | SGK494 |
| GPR107 | HP1BP3 | ZFP862-PS | | TSFM | PODNL1 | RP5-1054A22.4 | KRT16P6 | RP11-342A23.2 | ZNF598 | PEX7 |
| RHOBTB3 | TMEM65 | ZFP281 | | RAB3B | ACAP1 | S100Z | ARHGAP24 | TBX6 | PAX9 | SNRPF |
| 1190002F15RIK | GDAP2 | GPX1 | | ARRDC3 | FCGR2C | COL8A1 | XG | MIR4645 | CHSY1 | OR7E37P |
| URB1 | ACAD11 | SYT11 | | GPT2 | CTD-2576D5.4 | LRRN4CL | C4B | MYD88 | SOX10 | EID3 |
| DOCK11 | BC005561 | ZBTB8OS | | HDAC7 | RP11-329N15.3 | RP11-844P9.2 | UCN2 | POLR3A | SPINK1 | PRDX6 |
| SNHG1 | B230219D22RIK | 4930415O20RIK | | RAB21 | IGHV4-55 | CLDN11 | WNT2B | WDR38 | NEDD4L | C17ORF46 |
| UGT1A7C | NUP133 | MXRA8 | | MANEA | GLIS2 | CYP27A1 | KBTBD8 | FAM92B | ANXA7 | ICA1 |
| MICALL2 | ARID5A | NAP1L1 | | ITGA3 | IGHV3-13 | CEACAM4 | PILRA | ADCY5 | SPINT2 | RASSF5 |
| PARP16 | STAU2 | LY6C1 | | PPP3CB | TRAV14DV4 | FOXA1 | SCPEP1 | RHOBTB3 | TGM5 | SFT2D1 |
| SH3KBP1 | BCL11B | DNAJB5 | | 1700017B05RIK | CTD-2341M24.1 | GXYLT2 | NWD1 | CAMK2B | VARS | ZAP70 |
| PAXIP1 | PPWD1 | PAN2 | | SERPINA3F | IGKV3D-11 | MEIS1 | CD97 | BHMT2 | | ZNF84 |
| OGT | EIF3M | JPH1 | | SIRPA | CPVL | IGKV2D-29 | CTD-2267D19.1 | HNRNPK | | ASAP1 |
| GADD45B | 9330151L19RIK | PCGF5 | | CTSC | FXYD7 | LRRK2 | SERBP1 | CILP2 | | SOCS5 |
| SLC44A3 | RASGEF1B | TBC1D19 | | HSPA13 | TTC24 | SERPINB9P1 | IGKV1-13 | RALGAPA2 | | PEX5 |
| FUCA2 | LUC7L2 | EXOC8 | | UGT2B34 | KSR1 | DBNDD2 | LTBP4 | LINC01411 | | DACT3 |
| VWA5A | SLC25A37 | DENND1A | | IDH1 | ANKRD55 | KCNJ15 | PHKB | CDH20 | | MAP6 |
| WDR90 | PLA2G12A | SRA1 | | SMAP1 | TNFRSF8 | IGFBP3 | KCNH4 | RPS7P1 | | PPEF1 |
| GGA2 | MALT1 | PRCP | | CALM3 | S100A6 | LARS | PCDH10 | MEG8 | | SLC2A9 |
| ABCD4 | PRKAG2 | DSEL | | 1110059G10RIK | ADAM8 | PLEKHG5 | ST3GAL5 | RLTPR | | SLFN11 |
| TOMM7 | ETFDH | LOR | | HOMER2 | CCR2 | GNB2L1 | DHTKD1 | RPL15P3 | | CNTD2 |
| ELOVL6 | ALG8 | B4GALNT4 | | HEXDC | PIM2 | TRBV7-4 | KPNB1 | UBE2J1 | | KL |
| LAS1L | MGRN1 | SLC4A4 | | CCDC34 | CADM3 | GALNT11 | TMEM163 | CUL4A | | METTL11A |
| MMS22L | EML6 | STOM | | FAM103A1 | EVA1A | TINAGL1 | AC106786.1 | RP11-752L20.3 | | RARB |
| MINA | RPF2 | TRIM41 | | MRI1 | P2RY13 | LINC01504 | CCND2 | TRIM27 | | ACBD4 |
| LRRC20 | FRAT1 | NCOR2 | | BACE1 | SLFN11 | SDC4 | C10orf99 | SERPINB13 | | EIF4B |
| HERPUD1 | ZFP661 | EFNA3 | | SPINT1 | IGLC7 | RP11-322D14.2 | COG8 | CARD8-AS1 | | SLC38A11 |
| RPS3A1 | ANGPTL2 | COPS3 | | RYK | TMEM106A | MIXL1 | TSPAN15 | ZNF99 | | UNC5C |
| NAT9 | RPL4 | LYRM2 | | MIS18BP1 | CD14 | TLR3 | IMPACT | RPS17 | | DRAM1 |
| CHCHD10 | SERPINB1B | NSMCE1 | | SNX4 | KMO | AF127936.5 | MYO7B | WDR3 | | MFNG |
| MRPS25 | PIK3C2B | AI413582 | | TIMM9 | RGS18 | RASGRP4 | QARS | AC124789.1 | | ARHGEF5 |
| CD81 | CROT | ANO3 | | 9430008C03RIK | CAPN8 | RARA | ITPKA | BX470102.3 | | SLC22A8 |
| CHST2 | FAM168A | GSG2 | | ZDHHC2 | LINC00944 | KLHL23 | AMN | ATP5B | | SUN2 |
| AVPI1 | MVP | UBE2E2 | | ESCO1 | IGKV3-7 | SCARA3 | TDRD10 | RP11-710C12.1 | | ATP5L2 |
| MYL12B | RDH12 | NETO2 | | TRIM25 | BHLHE22 | RP11-764K9.2 | GSTM5 | CGA | | SLC30A9 |
| WARS | HRAS | DOCK5 | | GGT6 | FGR | PADI2 | CTD-3010D24.3 | SRGAP3-AS2 | | NCRNA00176 |
| DBF4 | NKAIN1 | USP14 | | ACNAT1 | RP11-750H9.5 | RARRES1 | RP11-225H22.4 | OGFRL1 | | C12ORF63 |
| TREX1 | EEFSEC | FAM46A | | DLK2 | CASP10 | GLI1 | PRDM6 | GDPD1 | | ZNF185 |
| KIF23 | NAT14 | SNX24 | | IGTP | TLR1 | LZTS1 | TCEAL5 | FABP6 | | ATP1B2 |
| RTN4IP1 | AP2A1 | SEC22B | | TGFBI | DNASE1L3 | SH2D1B | TMEM200A | CTC-457L16.1 | | UBAP2L |
| CARD14 | PDIK1L | REP15 | | KANSL3 | IGSF6 | RPL4P4 | MBOAT1 | ALDH1A3 | | ACTR2 |
| MLH3 | RNF166 | LIPT1 | | EIF3F | IKBKE | LINC01272 | VTI1B | RP11-1084E5.1 | | TMEM151B |
| RRP12 | FLYWCH2 | COA3 | | RASA1 | ST8SIA4 | IL33 | FAM162B | ZNF22 | | SLC3A2 |
| PLEKHG1 | KCNJ15 | RAMP3 | | LEPREL2 | TMEM150B | COL6A3 | MAP7 | METTL20 | | GZMB |
| SPOPL | ZFC3H1 | SPG11 | | SVIP | ZBED2 | KIF17 | PPP2R2A | ASTN2 | | ABCA1 |
| TAF9B | FAM57A | MAP2K5 | | SLMO2 | TRAV23DV6 | C22orf34 | LRRC75A-AS1 | U47924.31 | | CDC25C |
| CCDC136 | TUBB5 | PFKM | | IKZF2 | TRBV12-4 | RP11-148O21.4 | C10orf105 | IGF2-AS | | L1CAM |
| HDHD3 | PCNXL4 | TUSC1 | | VPS13A | TMEM140 | ZDHHC9 | ADCY3 | FRMPD3 | | LEPR |
| NDUFAF7 | UBQLN1 | YTHDC2 | | PGGT1B | RGS19 | U62631.5 | AC093616.4 | GLIS1 | | LOC401052 |
| THSD7A | SALL2 | NCEH1 | | CCNA2 | RASGRP1 | EIF3D | CCNG1 | BIVM | | MEI1 |
| RBM43 | CASP8AP2 | XPOT | | UHRF1BP1L | RAB38 | HNRNPA1P70 | GDF5 | KCNMA1 | | TLR6 |
| CAPRIN2 | LMF1 | VIL1 | | CYB5RL | GATM | IGHV3OR16-12 | GDF11 | GLI3 | | ESYT3 |
| GSDMC4 | CUTC | ACKR4 | | MYL7 | JDP2 | TMEM37 | CYSLTR2 | DET1 | | LOC100270804 |
| ADRB2 | TMEM63C | PQBP1 | | UBE2K | IGKV1D-17 | RP11-212I21.2 | SERPINA9 | SMOC2 | | C20ORF4 |
| CCNI | ARHGAP33 | D630039A03RIK | | CNOT2 | CCR8 | GPR153 | CTD-2575K13.6 | FAM173B | | MAP2K3 |
| ZFPM1 | TIMP4 | TSSC4 | | SEC24A | FUT7 | HDHD2 | RP4-753P9.3 | POMGNT1 | | RPL15 |
| SRSF1 | PDCD4 | NAV2 | | NUP133 | RP11-295P22.2 | VGLL1 | TCN2 | DLG4 | | LRRC3B |
| SCRN3 | MTTP | DGCR6 | | PLXNB1 | ABCD2 | RP3-467K16.2 | CTD-2373N4.3 | CCT8 | | C14ORF180 |
| GJB3 | 1600002K03RIK | JADE1 | | XIAP | PNMA5 | FCGR1B | STMN2 | NXPE4 | | GRIN3B |
| ARHGEF19 | UPK1A | AKNA | | ARL8B | XCR1 | TREM2 | CAMK1 | PCDH7 | | C2ORF49 |
| 2310009B15RIK | KIFC5B | FAM3C | | PRPF40A | RP11-148O21.2 | RP11-367G18.1 | MYOC | C12orf54 | | F10 |
| IRS1 | BCORL1 | PLCXD1 | | DNAJB4 | CCR6 | RPS6 | MMP17 | ADAM28 | | PRIM2 |

**Table S1. Significantly differentially expressed genes (EZH2 Signature High vs Signature Low, p<0.05) continued.**

| | | | | | | | | | |
|---|---|---|---|---|---|---|---|---|---|
| MAFF | SERTAD4 | TMEM41B | STEAP4 | SIGLEC9 | CLEC6A | LMO4 | CASC3 | | USP39 |
| PALD1 | DNTTIP1 | CDC42BPB | LPHN1 | VCAM1 | SIGLEC8 | TRPC2 | AIF1L | | EIF5A2 |
| ABCA7 | OLFML2A | NFKB1 | CYP26B1 | CEACAM6 | PIK3CD-AS1 | NKAIN4 | CCDC170 | | PHGR1 |
| ENO2 | TMEM154 | SKINT7 | ZFP820 | FMNL1 | SPARC | RP11-244H3.1 | RP11-263K19.4 | | VNN2 |
| SNHG5 | SMC2 | CEP44 | MAN2A1 | CTA-384D8.31 | TRPC3 | PSME2P2 | UNC93B1 | | C9ORF156 |
| NRN1 | CCPG1OS | DIAP1 | ATP5G2 | GRAP | CXCL6 | NF2 | ARFGAP3 | | LOC401588 |
| MYO18A | HGS | 2210016F16RIK | DSEL | RAB7B | CCDC89 | FBN1 | CRELD1 | | PLEKHG4B |
| HNRNPA3 | CAMK2D | CBFB | SOAT1 | IGHV3-43 | NACA | GADD45A | SMIM5 | | C11ORF42 |
| NOD1 | ADRBK1 | REXO2 | SLC11A2 | IGHV4-31 | PLAUR | EMR3 | GRAMD1B | | OR51I2 |
| SCEL | NUDT16L1 | TROAP | AP3M1 | TREML1 | RP11-126O1.6 | RP11-433J8.1 | IPP | | MST1P2 |
| EXOC3 | SGSH | WDR54 | ERN2 | IGKV3D-15 | ADAM23 | F10 | KCNF1 | | LOC100125556 |
| CCL9 | KCTD1 | FBXO5 | BMPR1A | FCRL4 | C16orf74 | UBE2O | RRN3 | | RIMBP3C |
| NUDT2 | FIG4 | AKAP2 | SDCBP | HSD17B13 | LTK | RAB5B | RP11-524D16__A.3 | | H3F3B |
| ATP6AP2 | TMEM44 | PRAF2 | TMEM87A | TRAV22 | ALPK1 | CDYL2 | CTSG | | RPL17 |
| CAPNS2 | PREPL | SFN | CDC27 | CARD17 | RPRD1A | VASH1 | CACNA1A | | TMBIM4 |
| APEH | ACBD6 | MRGBP | HIST1H2BC | HLA-H | ID1 | CCT2 | GTF3C2 | | GPC5 |
| PPM1B | ELAC2 | CHPF2 | RPS3A1 | KCNAB2 | FAM172BP | RP11-35N6.1 | HNRNPC | | KLHL2 |
| EPGN | ZFP62 | KDM4C | G6PDX | SCGB3A1 | PRCD | MITF | TMEM120B | | GOSR1 |
| RHOU | SYT7 | NKD2 | AA987161 | SRGN | TLDC2 | CPXM2 | EIF3B | | ISLR2 |
| SH2D4A | NME2 | INPPL1 | CXCL1 | GAS7 | CTGF | TARID | NOS3 | | LOC148413 |
| RPL36A | 0610010K14RIK | SLC8B1 | PPP2R2A | CLEC10A | CRYGN | TBC1D15 | P4HB | | LOC388796 |
| C2CD4B | ACKR3 | TUBA8 | AW549877 | MFAP2 | UPP1 | MTIF2 | TMPRSS7 | | LOC91316 |
| BOLA1 | PCBP3 | NUFIP1 | PRR15 | IFNG-AS1 | CCDC181 | MAVS | RP11-4C20.4 | | PAICS |
| SRSF2 | TUBGCP2 | RPL35 | STXBP6 | MTMR11 | ZNF74 | DNAAF2 | ZBTB42 | | PILRA |
| TRIM16 | 2810442I21RIK | CD177 | APOL9A | CYP2S1 | FN1 | HR | FOLR1 | | RPL26L1 |
| RBMX | PDE7A | RRP15 | DAP | RP1-50J22.4 | STK32B | AC104024.1 | AC005682.6 | | RPS16 |
| AKR1C13 | RAD51AP1 | RAB25 | THOP1 | PRAMENP | RGS13 | RP11-418J17.1 | RPRM | | SLC12A4 |
| GABPB2 | EVC | NAT2 | SEPP1 | ARHGEF2 | RNF166 | SAMD14 | EPHB1 | | SNORA62 |
| ENTPD5 | PAM | FUZ | MIER3 | RAB31 | RHBDL3 | VWCE | CDC42EP3 | | TTC7A |
| KNTC1 | GSTA3 | ZMYM5 | JAG1 | ADAMTS14 | CCNI2 | KRT19 | ARPC5 | | ZNF646 |
| PRODH | FAM171A2 | RAPH1 | GALNT7 | HCG4P11 | RP11-1334A24.6 | LILRA5 | HMGCR | | C4ORF10 |
| ADPRH | NCAPD2 | RPL6 | ATRX | TRAV2 | RN7SL834P | RP11-817I4.2 | SNAP23 | | ZNF688 |
| MFGE8 | ZFP637 | 1700092M07RIK | 1700096K18RIK | RP11-222K16.2 | GRK5 | RP11-54A4.2 | MKX-AS1 | | AGMAT |
| CAR13 | ADRM1 | ITIH2 | NUDT4 | ARHGEF6 | TNF | LINC00565 | PLTP | | CLEC17A |
| TMED3 | ZDHHC18 | DNAJC7 | GIGYF1 | NCCRP1 | MIEF1 | RP11-443P15.2 | RPL3P2 | | DNAJC8 |
| LDHD | TMEM106A | NPHP4 | SPG7 | PTPRE | RASGRF1 | TTC39B | KRT16 | | DNHD1 |
| TTBK2 | RAC1 | ERBB2 | AK3 | GBP3 | RP11-514D23.3 | C21orf91 | PRRC1 | | TMEM223 |
| PITRM1 | BC018242 | APEX1 | COMTD1 | ARRDC5 | CPA6 | TGFA | EDN1 | | GJA10 |
| ANKRD23 | METTL18 | POLR2L | TGM2 | PPP1R16B | NKD2 | HLA-V | RP11-358D17.2 | | LUZP1 |
| SYMPK | BHLHE22 | RHOBTB2 | DRAM1 | PTPLAD2 | CLIP3 | ADAMTSL5 | DHX33 | | C6ORF211 |
| HTRA2 | TGOLN1 | DNPEP | PAFAH1B2 | WNT10A | IGHE | PTCH2 | PLXDC2 | | LRRIQ3 |
| PLCD1 | GBA | NTN4 | PRUNE2 | AXL | RPL7AP6 | NGEF | CBX8 | | CLVS2 |
| MAP3K10 | HDDC3 | KIF20B | ROBO1 | IFI30 | OLFM4 | S1PR3 | CENPN | | GPIHBP1 |
| CD3EAP | PCOLCE | MKI67 | ARHGEF40 | PLXNC1 | HTRA4 | SYT3 | ZNF664 | | IFFO2 |
| TIAL1 | SRSF4 | GGACT | TGFA | COL16A1 | CD160 | MUT | CTB-41I6.1 | | PLAC8L1 |
| STIL | CENPN | PPM1M | TUBA1B | COL8A2 | DNAI2 | CTD-2114J12.1 | FST | | SNORA26 |
| OCEL1 | BB123696 | TTC12 | TAP1 | TRAV41 | RP11-796E2.4 | NEGR1 | COL24A1 | | TTLL13 |
| REEP4 | CDC7 | POU2F3 | COMMD3 | IGKV3D-20 | RP11-336K24.12 | INPP5B | MTPAP | | CCDC40 |
| POM121 | AMDHD1 | ANKRD42 | CAPN15 | PLAU | LINC01150 | C19orf48 | CDC20B | | GFER |
| IL1A | AGPAT2 | RAB7 | CCDC50 | RP11-326C3.2 | RPS4X | PVALB | NEXN | | EHBP1 |
| RPSA | MGST1 | SETMAR | MAPK12 | PHF11 | CEACAM3 | VCAN | TPK1 | | GPR34 |
| GM5431 | PLEKHA6 | D930016D06RIK | TMEM106B | PRKCQ | RP11-236L14.2 | MORN3 | STT3A | | LOC100129726 |
| COL4A2 | GTPBP8 | FXR1 | CEACAM1 | TRAF5 | RP11-317P15.4 | CEACAM7 | CCDC138 | | CDRT15P |
| PFKFB4 | PUS7 | MAX | FDFT1 | TRAV8-1 | CRYBB1 | SPINK2 | CHERP | | IFIT2 |
| KHSRP | CRELD1 | RMND5A | FMO3 | RP11-1166P10.8 | hsa-mir-4538 | RPS3A | ANKRD53 | | RIN3 |
| MAPK1IP1 | MICAL3 | CMBL | OAS3 | RIN3 | CHRDL2 | RPS23 | CFTR | | MIP |
| LAPTM4A | KRT78 | FKBPL | NPC1 | PYCARD | AMPH | RP11-44K6.2 | PTGER2 | | PDE4A |
| LSR | DAGLA | B3GLCT | SSNA1 | DDX60L | PM20D1 | CORO2A | DPF3 | | RNF146 |
| TFB1M | CADPS2 | TOR1AIP1 | TMPRSS11BNL | HLA-DQB1 | TCTEX1D1 | SLC38A3 | LINC01506 | | TMEM192 |
| ACAD10 | LMO2 | MFSD2B | TSC22D1 | AC018816.3 | DACT3 | GUCY2C | TSPAN14 | | TBC1D29 |
| THBS3 | HOMER3 | IL1RL2 | NUBP1 | ALS2CL | CYP2T1P | EOGT | GNAI1 | | ASB6 |
| NFE2L2 | CERS4 | NADK | BCAP29 | CD1A | RPL3 | MARCO | ZNF793-AS1 | | C11ORF34 |
| ENOX2 | RPL7 | ZDHHC21 | PIH1D1 | NME8 | CACNA1I | PPP1CB | PHLDA1 | | CD5 |
| LONP2 | RAB11FIP5 | LY6G6D | ANTXR1 | CTA-212A2.1 | PLSCR4 | KLRG1 | CECR2 | | GJA4 |
| CCND1 | PPAP2A | DYNLL2 | PPP4R1L-PS | C3AR1 | MAGI2-AS3 | PSCA | NID1 | | LOC100134229 |
| MROH2A | SIX5 | CLCN4-2 | KRT1 | TRDC | CRYL1 | RSL24D1 | LINC01374 | | TAC3 |
| MRPS6 | DENND3 | BRIP1 | ACSL4 | ABI3BP | CHST15 | XXYLT1-AS2 | NUPL1 | | KRT86 |
| CCDC171 | ITPR2 | ZFP593 | PICALM | GIMAP5 | FGD6 | TCEB3 | RPL7 | | SEL1L3 |
| PDCD11 | RNPC3 | RAB27A | ATP6V1C2 | IGKV1D-16 | RPL7AP64 | HOTAIR | C1RL-AS1 | | PCNAP1 |
| UBE2C | PCED1A | FRAT2 | SLC25A48 | NLRP3 | IRF1 | SNAI3 | GRASP | | SLC35E4 |
| GM12942 | LTB4R1 | THA1 | RNASE4 | FPR3 | KCTD3 | RP11-701H16.4 | FUBP3 | | WIPF1 |
| ESPL1 | WIZ | ESF1 | ANP32E | PAX5 | SLC16A7 | RP11-588H23.3 | ERLIN2 | | DNAH17 |
| TXNDC12 | 2810403A07RIK | ZFP821 | LDLRAD3 | TRAV12-1 | EHBP1L1 | RP11-551L14.1 | FNBP1 | | EFNB2 |
| SLCO4A1 | 43351 | FAIM | RSRC2 | RAB33A | LINC00987 | RP11-517I3.2 | ENTPD3 | | INMT |
| TINAGL1 | PHF19 | ITGB5 | WWTR1 | MMP9 | IGHV3-25 | HMGXB3 | ACTBL2 | | OR1C1 |
| GADD45G | BZW1 | METTL9 | RAD21 | LOXL3 | RP11-490G8.1 | SLC35F2 | TDRKH | | CLDN10 |
| AMY1 | PLAGL1 | 2810001G20RIK | SOCS6 | TRBV6-6 | RP11-936I5.1 | JAM2 | CCL2 | | CYB5R3 |
| GPCPD1 | TOR1AIP2 | MBTPS1 | PLXNA2 | IGKV2-30 | KIAA2012 | RP11-91K8.4 | FBXO2 | | ZBTB8A |
| SLC37A3 | MFAP3L | GM10069 | CTGF | TMCC2 | TCEAL2 | EEF1A1P6 | YEATS4 | | LSM10 |
| ISCU | RARB | SARM1 | HOOK3 | LMO2 | PLD1 | LINC00942 | DNM1P35 | | SMARCC1 |
| GALNS | METTL21A | ALKBH8 | SLC39A10 | PMP22 | RP11-394O4.5 | DHRS11 | KIF5A | | SNHG7 |
| LGALS3 | SBNO2 | NAT6 | JADE1 | ETS1 | MRPS35 | AGBL5 | MBNL1-AS1 | | ZNF354A |
| ANXA9 | LRRC57 | NSMF | EML2 | PTGIR | DSC2 | CLPX | INSM2 | | LPXN |
| RPLP1 | RAVER1 | CEP97 | PLTP | HMOX1 | IGLV10-54 | FCGBP | MORN4 | | ATAD2 |
| TTC30B | PPCDC | 2610015P09RIK | CDH24 | L3MBTL4 | RP11-370I10.12 | RP11-473M20.16 | IL23R | | FMO3 |
| SVIP | SFXN1 | FADS2 | RPL27A | CLEC12A | RAB3IP | IGKV2-26 | RP11-234A1.1 | | PCBP1 |
| SLC39A10 | PIGF | TYRO3 | RNF128 | RASA3 | SLC35E1 | COX7A1 | GPR17 | | BDNFOS |
| CXCR4 | SCOC | MPC1 | LY6G6E | CACNA2D4 | P2RX2 | AFMID | SYNJ2BP | | MYO1H |
| CCNE2 | TRIM30D | 0610012G03RIK | HOMER1 | IGLV5-45 | PLEKHG1 | CECR6 | IGKV1OR2-3 | | OXER1 |
| RPP38 | VCAN | BCAR1 | UTP20 | MS4A14 | BCL2L13 | DNM3 | NUMA1 | | ZNF221 |
| TXN2 | ZFP467 | PTGES | IRGM2 | MMP2 | LINC00868 | HOXB6 | VPS54 | | COX19 |
| CRABP2 | SNX27 | DAD1 | SAR1B | IGKV1-16 | DSG4 | RAP2B | RP11-431J24.2 | | HINT3 |
| SBSN | GPR108 | CEP170B | STAM | PDLIM1 | KIAA1045 | AC012360.6 | RP11-253E3.1 | | CLSTN1 |
| PLCD3 | AURKAIP1 | SF3A2 | WTIP | TAGLN2 | WNT7A | RP1-266L20.2 | HLA-DMB | | LOC728276 |
| ADAMTS1 | FDXR | GGA3 | LGALS9 | P2RY6 | IL21-AS1 | MMP23B | PLEKHF2 | | SEMA7A |
| PRKAR2B | RASA3 | 2510039O18RIK | HPN | LAYN | HPN | ASPA | PGR | | ESD |
| OLFR1372-PS1 | ACOT13 | CACTIN | 4930506M07RIK | CSF3R | NUAK1 | NSUN2 | POLD2 | | LCN10 |
| 2700089E24RIK | PECAM1 | ISOC2A | HAP1 | RP11-532F6.3 | SH2D3C | GTSF1L | MLST8 | | LOC100133612 |
| JUN | PRC1 | TIMM13 | AHCY | EFEMP2 | GEM | COX6B2 | SEMA6D | | PALM |
| ECHDC1 | MMACHC | STC2 | ALS2CL | P2RY14 | DST | ACTB | MRPS2 | | RPS25 |

#### Table S1. Significantly differentially expressed genes (EZH2 Signature High vs Signature Low, p<0.05) continued.

| | | | | | | | | | |
|---|---|---|---|---|---|---|---|---|---|
| HSPA4L | NUDT18 | ARRDC4 | ZFP750 | TRBV5-4 | PNMAL2 | RPS7 | ZSCAN16 | | PCDHGC3 |
| CD274 | TNFSF15 | CELSR1 | MTDH | TPRG1 | HSH2D | RP11-217B1.2 | TRGV3 | | C21ORF121 |
| CLCA5 | PTK2 | GGT6 | RRM2B | ADCY7 | PARP10 | GP1BA | RP11-480C22.1 | | SAR1B |
| NAGLU | SNX16 | PSMD1 | PML | CTD-2095E4.5 | FAM177B | ZFR2 | NR2F1 | | LOC541473 |
| EIF2A | WHRN | SCARA3 | CERS3 | TRBJ2-7 | RP11-81H14.1 | LONRF3 | ADRB1 | | GBP4 |
| RPS11 | MND1 | CENPK | ZFP800 | ITGA4 | SHANK2 | FENDRR | EEF1A1P19 | | NCF1B |
| CDC34 | RHBDL1 | SLCO3A1 | LY6E | RP11-24F11.2 | CHI3L2 | MYCN | REPIN1 | | SP110 |
| HILPDA | UHRF1BP1L | DTX3 | MED7 | LINC01281 | CPZ | BCAM | PNMA2 | | TBC1D7 |
| RPS6 | R3HDM4 | TMEM50A | COQ2 | PTGS1 | MYOF | C1QTNF7 | SRP54 | | TMEM102 |
| PRKCA | ALDH5A1 | KIF22 | 1810032O08RIK | TRAV27 | LEO1 | MIR205HG | MRPS27 | | ZNF642 |
| RPS5 | RAB6A | EXOC5 | KRCC1 | RP11-960L18.1 | LINC01268 | CLEC4D | RP11-514D23.2 | | RBM42 |
| CBX5 | SNORD47 | RAB24 | NPM3-PS1 | AP000476.1 | DCN | GDF7 | CRYAB | | STAT4 |
| CFL2 | MDC1 | HOOK1 | KLF3 | IGKV1D-8 | BEND4 | ZDHHC20P1 | IRF2BP1 | | TTLL7 |
| PARPBP | TRF | SLC7A5 | TGIF2 | RP6-159A1.4 | CD83 | KIRREL3 | SIPA1 | | NDUFAB1 |
| PPIF | GM14827 | TMEM110 | WWP1 | FCGR1A | RP11-849N15.3 | LANCL2 | TRIP10 | | PCA3 |
| ITPR3 | ITGA9 | 2300009A05RIK | OGFR | IGLV7-46 | C7orf31 | MPHOSPH10 | C18orf32 | | ADAM10 |
| HEPACAM2 | DSG3 | SYNJ2BP | FGF1 | RP11-750B16.1 | RP1-244F24.1 | RBMS3 | ITGA8 | | CARD10 |
| MTR | EHD2 | PIWIL4 | TCEB1 | TRAV6 | NXPH3 | GDI2 | BRS3 | | DNAJB9 |
| FES | PHYHD1 | PEX5 | STX1A | IGKV1-33 | PSORS1C1 | PEX5 | TOX3 | | SNX15 |
| RPS15A | CYLD | TLR4 | ANGPTL2 | MGP | NCOA7 | SLC5A10 | CA8 | | TMEM19 |
| CHST10 | RPL8 | GM10433 | ANKRD6 | IGKV1-12 | LY6E | IGLV2-5 | DZIP1 | | CDKN1B |
| SCNN1A | KHDC1A | ULK3 | LAMA5 | SMTNL1 | CREB3L4 | LINC00665 | GSTM4 | | HIST1H2AG |
| SPRYD4 | HIST1H3A | FNDC3A | YBX3 | SIRPA | BDH2 | MYO3A | S100A2 | | P2RY12 |
| NAIP5 | B930041F14RIK | NEK2 | BC021767 | IGHV3-35 | RP11-536C10.25 | RP11-468E2.4 | SLC47A1 | | C17ORF61 |
| FTL1 | SLC39A14 | SRI | PROCR | ST6GALNAC4 | GSTM2 | RP11-153M7.3 | IL17RE | | LOC729156 |
| LLPH | CAR15 | AMPD2 | ALOX12 | C5orf56 | CTD-2353F22.1 | RP11-11N9.4 | KCNH3 | | N4BP3 |
| A730046J19RIK | CRNKL1 | LRRC14 | SEMA3C | IGHJ3P | HTR2B | MRC1 | KRT17P2 | | NUDT9P1 |
| NFIC | PSMC6 | SETD1B | ABHD11 | IGKV2D-24 | VAV3 | GCK | ZNF253 | | PARP12 |
| EIF2S3Y | SLC39A11 | 1810043G02RIK | INO80C | DOCK11 | MRPL45 | FAM98B | CPSF7 | | TRIML2 |
| PADI4 | RNF13 | CHD6 | RAB6A | 43347 | MXRA8 | EFHD1 | CYLD | | ZMYND11 |
| KIF24 | ZFP219 | NCCRP1 | TAOK1 | IGHV3-38 | RUNX1T1 | ANK2 | ALS2CR12 | | FAM76A |
| RCL1 | ZFAND3 | SNX1 | GOLPH3L | MOB3C | IFNE | ITGA11 | LINC00284 | | C11ORF63 |
| ANGEL2 | KRT80 | RASSF8 | GTF2A1 | HLA-DQB1-AS1 | CD163 | YPEL1 | PPP1R26 | | PRKAR2A |
| H1FX | ANKRD50 | LDB1 | RAB34 | VAMP5 | L3MBTL4-AS1 | FCER1A | RP11-536C10.1 | | RPL24 |
| CAD | U2AF1L4 | TMEM54 | CLIP4 | CD300C | VMO1 | HMGB1P3 | HEATR5B | | TMEM129 |
| RIN3 | ANPEP | DRP2 | PPIG | CD200 | ANO1 | SLC16A5 | GP5 | | TMEM105 |
| HELLS | CENPI | ZFP764 | RNF115 | RP11-383H13.1 | EGFLAM | UCN3 | LAMA2 | | DTYMK |
| SSSCA1 | LY6G6C | PISD | SHMT1 | GCSAM | APOBEC3A | ACOX1 | TJP1 | | KCNC4 |
| PRKAB1 | MBLAC1 | PEO1 | SAMD5 | SYNGR3 | NXPE1 | RERE | NOP9 | | STK10 |
| GFRA4 | RETSAT | PTOV1 | LAMB2 | KIAA1755 | FAM20A | EARS2 | RNASEL | | FAM92A1 |
| MMP28 | PRAM1 | BCKDHA | CFL2 | ANTXRLP1 | TMTC4 | MPZ | METAP1D | | C8ORF77 |
| TK1 | GM7694 | KBTBD8 | WDR90 | IDO2 | L1TD1 | CCDC144NL-AS1 | WSCD2 | | LRDD |
| IQGAP3 | RBMS1 | TAF10 | TAB3 | CMTM4 | SH3BP5 | CCT7 | RPL13AP3 | | ZNF169 |
| RPL5 | RASIP1 | PIH1D1 | 1110038B12RIK | VNN2 | RPL15 | SERPINH1 | DNAJC16 | | KIAA0748 |
| LRFN4 | SPRR2E | SERPINB10 | FLNA | IGHV1OR15-9 | FILIP1L | C22orf15 | AC093850.2 | | NFXL1 |
| TJAP1 | ABI3 | RPL21 | NDFIP1 | EEF2 | IL13RA2 | AL357515.1 | PDLIM4 | | ALKBH4 |
| SPN | IWS1 | TMEM45B | AI661453 | M1AP | RP4-671O14.5 | COL11A1 | AC064834.3 | | CPA5 |
| AI607873 | AP4B1 | RPS27A | SIVA1 | LIMD2 | LAMA4 | AC011515.2 | NBPF8 | | OXT |
| SNAP29 | ARHGAP40 | ABL1 | ANK | GNG2 | SPNS2 | RPS2P7 | IGKV6D-21 | | KIAA1377 |
| ACAD9 | NEURL1A | RAPGEF5 | GNS | CCDC88B | ITGB4 | VAMP1 | TTC39C-AS1 | | DLL1 |
| PPT1 | RNF31 | TAGLN2 | MOSPD1 | IL22RA1 | ZNF267 | EDN2 | RP11-554A11.5 | | ATP1A3 |
| ICAM5 | ART4 | CTSZ | INF2 | ISLR | ENTPD1 | HSF5 | PGM5P4-AS1 | | TNFAIP6 |
| FAM65C | CCDC130 | KIF1C | CLDN1 | HAMP | MEOX1 | SLC16A1-AS1 | RP11-180I4.4 | | C21ORF57 |
| SPG20 | REM2 | BCL9L | MID2 | TDRD7 | HOXD13 | ENG | SYNGR2 | | FUT5 |
| C1QTNF6 | CXCL11 | STAT6 | PTGS1 | HOXB2 | WBP11 | LINC00032 | RP11-713P17.3 | | KIAA1009 |
| DAB2IP | SERPINA3G | PRR15 | TRIM14 | IGHV3-41 | DIRAS3 | RP11-118B22.4 | RP11-467H10.2 | | MSRB3 |
| ANKRD37 | CTNNB1 | RC3H2 | CHN2 | MEG3 | LGALS9C | SAPCD2 | ZNF101 | | SLC9A5 |
| SH3D19 | GATSL2 | HIST1H4D | RRAS2 | BLACAT1 | GNG8 | RAMP3 | LMO3 | | TMEM31 |
| 2310001H17RIK | PSME4 | SENP6 | 2610002M06RIK | FSD1 | RP11-459I19.1 | KLF5 | HOXA7 | | ZMIZ1 |
| SLC10A2 | ATP1A1 | GMNN | CAMSAP2 | IGLV2-18 | ZSCAN2 | CTD-2003C8.2 | SLC43A1 | | ZNF594 |
| VAV3 | CCL5 | RFFL | TOX3 | NPFFR1 | BEND5 | FUT2 | GPR78 | | CLDN6 |
| CXCL17 | RPH3AL | LRFN3 | RPS6KA2 | ANKRD36BP2 | AVPI1 | RP11-340F14.6 | AP001626.1 | | PLEKHA6 |
| SPRED3 | MPP2 | KDM1A | EIF4EBP2 | TRAV20 | RP11-16P20.4 | RNF223 | C1QL1 | | NCKAP5 |
| PINLYP | EXOG | UNC93B1 | RAB5A | CCRL2 | RHOV | RP11-247C2.2 | EIF3A | | HIST1H3J |
| ZFP523 | BRD3 | TLDC1 | ATG5 | IGHV3-19 | XPNPEP2 | SPSB1 | RILPL2 | | PTGES |
| FAM46C | GSTZ1 | PRKAA1 | CLCA4 | IGKV6-21 | PDE3B | MUC20 | N4BP2L1 | | RTN4R |
| GEMIN5 | PPP4C | GSDMCL-PS | PDGFA | ARPC1B | CTD-3252C9.4 | RP11-243M5.2 | OXSM | | SSR3 |
| CSTA1 | SNX33 | FANCG | D8ERTD82E | TMEM51 | RHOJ | RP13-580F15.2 | CYFIP1 | | DGKH |
| PLAUR | TRIB2 | GLUL | MOB3B | RP3-477O4.14 | ZNF80 | IGHJ1 | RP1-29C18.8 | | DISP2 |
| CAV1 | ZZZ3 | PYURF | ATF1 | BTN2A3P | IGLL1 | DDR2 | SMCO4 | | TNRC6C |
| PCMT1 | BRI3BP | EFNA1 | API5 | KLRK1 | KIRREL | PLB1 | ARHGEF25 | | PCDH15 |
| CSK | SRGAP3 | ADD2 | CAV1 | MIR4435-1HG | AC002331.1 | RP11-132F7.2 | EEF1GP5 | | ACSBG1 |
| HAGHL | 5430435G22RIK | PI4KA | PWP2 | TRBV5-5 | SDR16C5 | POLR1A | AQP9 | | ALS2CR4 |
| ZC3HAV1 | POMK | KMT2B | DSC2 | P2RX7 | BTF3 | RPL13AP5 | RP11-449D8.1 | | C16ORF45 |
| CLPP | MIR24-2 | COQ7 | STX7 | AC244250.2 | SNX7 | ARL4D | AC069363.1 | | C3ORF33 |
| UTP14B | PARK7 | TMCO6 | OLFML2A | MSN | EMILIN1 | KRT7 | ENO2 | | COX7C |
| FBXO11 | ATXN2 | EPG5 | SLC29A1 | PIK3CG | ARSD | VWA9 | MAPKAPK5 | | HEG1 |
| MZT1 | GNB2 | CENPH | HOXA10 | BST1 | RP11-51F16.1 | UBE2E2 | INCA1 | | LAMB2L |
| RPS16 | RPL35A | BEND5 | SWT1 | RHBDF2 | TRIM47 | IQSEC3 | RASA4 | | LGR6 |
| TFAP2A | ERCC4 | ESRP1 | RB1CC1 | IGKV2D-40 | ID3 | RP11-493L12.4 | PRSS51 | | SPTBN5 |
| E130008D07RIK | DBN1 | YIPF6 | RAB9 | UCP2 | TSPAN2 | RPS24 | ZNF423 | | THBS2 |
| SPRR2D | PGM2 | GMEB2 | RMND5B | TRBV15 | RP11-498E2.9 | ST3GAL2 | RP11-442O1.4 | | URM1 |
| EPHX1 | B230208H11RIK | TOPBP1 | FBXO11 | TRBV24-1 | ADAMTS12 | SORCS2 | COLCA2 | | ZMYM1 |
| CSNK2A2 | PAPSS2 | FBLIM1 | UNC5A | AKR1B1 | CD248 | SOWAHD | ATP1B2 | | ZNF547 |
| ZNFX1 | MMGT2 | CFDP1 | THAP2 | IGHV3-20 | MEIS2 | RPS18P9 | PRELID2 | | ZNF248 |
| PM20D2 | DQX1 | FLNB | CTDSPL2 | PSME2 | REEP2 | GMIP | RP11-280K24.4 | | DNAJC21 |
| RRS1 | DAXX | NAA10 | MTMR1 | MS4A7 | RP11-6I2.3 | GFPT1 | CCL11 | | BMP8A |
| COA4 | LENG9 | ANKS1B | NOXO1 | GYPC | SLC7A1 | SPON1 | ARSH | | KIF4B |
| MAST2 | ST8SIA6 | PRRC2A | VGLL3 | MECOM | KRT87P | HRH1 | SLC38A5 | | RPL23AP64 |
| RBM15 | TRIM56 | SLC16A1 | PTPRJ | ZNFX1 | NXNL2 | RP11-563J2.3 | FCN1 | | COX7A2L |
| KRT32 | EFCAB4A | DHRS7 | 2810008D09RIK | S100A3 | TCEAL7 | TCF4 | PDCD11 | | FMO6P |
| PCBP4 | GALNT14 | CHRNB1 | UNG | TRBV27 | SERPINI2 | GOLGA2 | SLC5A5 | | KATNAL2 |
| P2RX2 | GCSH | KLHL36 | HMGN3 | C9orf3 | RP11-108M9.4 | RP11-549B18.1 | KRT18 | | SETD6 |
| CUL3 | SLCO2A1 | DHX33 | GEMIN5 | IGKV1D-43 | RP11-395B7.4 | ZNF711 | ZFPM2 | | LOC730668 |
| SRPK1 | GRIN1 | RMI1 | TTF2 | RP11-357H14.17 | GSDMC | RASGRF2 | SEC16A | | MLF1 |
| AKT3 | NUCB2 | NOP58 | ZFR | FHAD1 | VSNL1 | RPL13AP7 | NAA60 | | NLRC3 |
| LUC7L3 | VIMP | 5430417L22RIK | NAPG | SNCAIP | PDE9A | SLC6A16 | RIIAD1 | | NRGN |
| GLT8D1 | SS18 | ACTG1 | SRI | PLEKHA4 | CTSE | KDF1 | CTD-2325P2.4 | | C4ORF21 |
| RPS26 | MEX3B | FABP2 | CWC15 | CFI | NLRC4 | RPL12 | GPAM | | PDZD8 |
| NUDT14 | SMYD4 | PLD3 | NDUFA12 | FGL2 | CCL28 | MIR222HG | TRIM24 | | PRPH |

**Table S2: Master regulator analysis of EMC 3D organoids treated with DZNep vs DMSO (FDR < 0.05, Top 50)**

| Regulon | Size | NES | p.value | FDR |
|---|---|---|---|---|
| STAT1 | 34 | 8.88 | 6.42E-19 | 1.61E-15 |
| IRF9 | 40 | 7.6 | 3.07E-14 | 3.86E-11 |
| SP110 | 41 | 5.9 | 3.67E-09 | 3.07E-06 |
| STAT2 | 10 | 5.16 | 2.46E-07 | 1.54E-04 |
| PARP14 | 29 | 3.66 | 2.49E-04 | 1.25E-01 |
| TRIM22 | 146 | 3.29 | 1.02E-03 | 3.19E-01 |
| SOD2 | 84 | 3.05 | 2.29E-03 | 5.26E-01 |
| ZNFX1 | 23 | 3.05 | 2.30E-03 | 5.26E-01 |
| PURB | 17 | 2.91 | 3.59E-03 | 5.79E-01 |
| MRPL28 | 81 | 2.88 | 4.00E-03 | 5.79E-01 |
| NMI | 53 | 2.87 | 4.06E-03 | 5.79E-01 |
| PCBD1 | 200 | 2.84 | 4.52E-03 | 5.79E-01 |
| VPS25 | 298 | 2.78 | 5.51E-03 | 5.79E-01 |
| SCGB1A1 | 61 | 2.74 | 6.17E-03 | 5.79E-01 |
| NMRAL1 | 195 | 2.7 | 6.99E-03 | 5.79E-01 |
| HNRNPAB | 161 | 2.67 | 7.59E-03 | 5.79E-01 |
| NDUFA13 | 163 | 2.67 | 7.62E-03 | 5.79E-01 |
| TRAFD1 | 20 | 2.62 | 8.87E-03 | 6.03E-01 |
| MAP2K1 | 201 | 2.46 | 1.39E-02 | 7.26E-01 |
| IFI16 | 123 | 2.45 | 1.45E-02 | 7.26E-01 |
| ZNF705D | 26 | 2.4 | 1.65E-02 | 7.26E-01 |
| PRDX3 | 264 | 2.37 | 1.76E-02 | 7.26E-01 |
| TCEB2 | 175 | 2.37 | 1.77E-02 | 7.26E-01 |
| ZNF622 | 18 | 2.34 | 1.94E-02 | 7.26E-01 |
| MLX | 134 | 2.29 | 2.21E-02 | 7.31E-01 |
| SNF8 | 113 | 2.28 | 2.24E-02 | 7.31E-01 |
| POLR2I | 163 | 2.19 | 2.85E-02 | 8.06E-01 |
| MORC4 | 185 | 2.18 | 2.91E-02 | 8.06E-01 |
| SAP18 | 54 | 2.18 | 2.92E-02 | 8.06E-01 |
| ZNF557 | 16 | 2.17 | 3.03E-02 | 8.07E-01 |
| POLR3K | 34 | 2.15 | 3.15E-02 | 8.07E-01 |
| PFDN1 | 90 | 2.15 | 3.18E-02 | 8.07E-01 |
| ZNF793 | 24 | 2.12 | 3.39E-02 | 8.07E-01 |
| VGLL1 | 27 | 2.11 | 3.46E-02 | 8.07E-01 |
| PSMC3 | 167 | 2.11 | 3.49E-02 | 8.07E-01 |
| PPM1A | 104 | 2.1 | 3.62E-02 | 8.07E-01 |
| ELOF1 | 118 | 2.07 | 3.82E-02 | 8.07E-01 |
| PRDX2 | 164 | 2.06 | 3.94E-02 | 8.07E-01 |
| TLR4 | 49 | 2 | 4.54E-02 | 8.07E-01 |
| HIVEP2 | 12 | 1.98 | 4.76E-02 | 8.07E-01 |
| STAT3 | 71 | 1.97 | 4.94E-02 | 8.07E-01 |
| ZNF662 | 14 | 1.96 | 4.94E-02 | 8.07E-01 |
| ZNF81 | 24 | 1.96 | 4.97E-02 | 8.07E-01 |
| C14orf156 | 127 | 1.95 | 5.15E-02 | 8.07E-01 |
| PDCD4 | 31 | 1.94 | 5.23E-02 | 8.07E-01 |
| ZRSR2 | 11 | 1.93 | 5.33E-02 | 8.07E-01 |
| SMARCD2 | 241 | 1.93 | 5.34E-02 | 8.07E-01 |
| SCYL1 | 131 | 1.92 | 5.51E-02 | 8.07E-01 |
| PCGF1 | 181 | 1.89 | 5.83E-02 | 8.07E-01 |
| DDX5 | 7 | 1.88 | 5.95E-02 | 8.07E-01 |

**Table S3: EZH2 Repression Signature**

| Gene | Weights |
|---|---|
| IFIT1 | 189.5737206 |
| MCAM | 126.9552146 |
| LGALS3BP | 104.5763895 |
| XAF1 | 103.2764477 |
| IRF7 | 102.3259138 |
| CXCL10 | 94.91667878 |
| PPBP | 93.07112204 |
| ISG15 | 87.77826856 |
| HR | -81.50172117 |
| KRT10 | 69.51001869 |
| BST2 | 56.71217579 |
| RSAD2 | 55.69539853 |
| IFI44 | 51.66033176 |
| OAS3 | 43.00782536 |
| KRT1 | 41.06430867 |
| LGALS9 | 39.61557658 |
| DHX58 | 36.77590278 |
| RTP4 | 36.66686165 |
| SCGB1A1 | 36.45179926 |
| MX2 | 35.51494032 |
| GBP7 | 32.48968587 |
| DDX60 | 30.35789776 |
| IFIT3 | 26.45578194 |
| SP100 | 25.93567214 |
| CMPK2 | 22.59376575 |
| GBP3 | 21.46126774 |
| OAS2 | 17.90711659 |
| ZBP1 | 16.06445844 |
| NLRC5 | 15.55581122 |

#### Table S4: Details of transcriptomic datasets in study

| Dataset | Samples (Total) | Samples (Included) | Subtype (Included) | Reference [PMID] |
|---|---|---|---|---|
| **Mouse** | | | | |
| EMC Organoids - DZNep vs DMSO | 6 | 6 | Prostate Adenocarcinoma | Unpublished |
| EMC Organoids - Tam vs Ethanol | 4 | 4 | Prostate Adenocarcinoma | Unpublished |
| **Human** | | | | |
| LNCaP - EPZ vs DMSO | 6 | 6 | Prostate Adenocarcinoma | Kim *et al.* Cell Rep 2018 |
| TCGA - PRAD | 449 | 408 | Prostate Adenocarcinoma | Provisional |
| Trento/Cornell/Broad | 47 | 33 | Prostate Adenocarcinoma (mCRPC) | Beltran *et al.* Nat Med 2016 |
| SU2C/PCF Dream Team | 212 | 212 | Prostate Adenocarcinoma (mCPRC) | Abida *et al*. PNAS 2019 |
| NCI - Sowalsky | 17 | 17 | Prostate Adenocarcinoma | Unpublished |

#### Table S5: List of antibodies used in experiments

| Antibody | Catalog | Source | Application | Concentration |
|---|---|---|---|---|
| EZH2 | 5246S | Cell Signaling Technology | IHC | 1:200 |
| PD-L1 | 13684S | Cell Signaling Technology | IHC | 1:400 |
| dsRNA | J2 | Scicons | FC, IF | 1:100 |
| CD3 | 100210 | BioLegend | IF | 5 µg/mL |
| CD4 | 100446 | BioLegend | IF | 5 µg/mL |
| CD8α | D4W2Z | Cell Signaling Technology | IF | 1:200 |
| Arginase-1 | D4E3M | Cell Signaling Technology | IF | 1:50 |
| iNOS | D6B6S | Cell Signaling Technology | IHC | 1:200 |
| Tri-Methyl-Histone H3 (Lys27) | 12158S | Cell Signaling Technology | FC, IHC | 1:100 |
| Cleaved Caspase 3 | 9602S | Cell Signaling Technology | FC, IHC | 1:100 |
| Ki-67 | 12075 | Cell Signaling Technology | FC | 1:100 |
| Ki-67 | 12075 | Cell Signaling Technology | IHC | 1:200 |
| PD-L1 (mouse) | 124308 | BioLegend | FC | 0.25 µg/$10^6$ cells |
| PD-L1 (human) | 374512 | BioLegend | FC | 5 µL/$10^6$ cells |
| CD45 | 103122 | BioLegend | FC | 0.25 µg/$10^6$ cells |
| CD45 | 103138 | BioLegend | FC | 1:100 |
| CD3 | 100306 | BioLegend | FC | 1:100 |
| mCD8 | 100750 | BioLegend | FC | 1:100 |
| hCD8 | 301010 | BioLegend | FC | 1:100 |
| CD4 | 100548 | BioLegend | FC | 1:100 |
| PD-1 | 135228 | BioLegend | FC | 1:100 |
| CD11b | 101263 | BioLegend | FC | 1:100 |
| Ly6C | 128044 | BioLegend | FC | 1:100 |
| Foxp3 | 20-0191-U100 | Tonbo Biosciences | FC | 1:100 |
| Ly6G | 80-5931-U100 | Tonbo Biosciences | FC | 1:100 |
| 4-1BB | 106106 | BioLegend | FC | 1:100 |
| I-A/I-E | 107632 | BioLegend | FC | 1:100 |
| anti-PD-1 | BE0273 | BioXCell | *in vivo* | 20 µg/treatment |
| anti-PD-1 | BE0273 | BioXCell | MLR | 10 µg/mL |
| IgG | BE0089 | BioXCell | *in vivo* | 20 µg/treatment |
| IgG | BE0089 | BioXCell | MLR | 10 µg/mL |

#### Table S6: List of oligo and primer sequences used in experiments

| Primer | Application | Sequence |
|---|---|---|
| 60S Ribosomal Protein L32 - forward | qRT-PCR | TTCCTGGTCCACAACGTCAAG |
| 60S Ribosomal Protein L32 - reverse | qRT-PCR | TGTGAGCGATCTCGGCAC |
| PD-L1 - forward (mouse) | qRT-PCR | CAGCAACTTCAGGGGGAGAG |
| PD-L1 - reverse (mouse) | qRT-PCR | TTTGCGGTATGGGGCATTGA |
| PD-L1 - forward (human) | qRT-PCR | TGGCATTTGCTGAACGCATTT |
| PD-L1 - reverse (human) | qRT-PCR | AGTGCAGCCAGGTCTAATTGT |
| PSA-Cre(ERT2) - forward | genotyping | ATCCGAAAAGAAAACGTTGA |
| PSA-Cre(ERT2) - reverse | genotyping | ATCCAGGTTACGGATATAGT |
| MYC - forward | genotyping | AAACATGATGACTACCAAGCTTGG |
| MYC - reverse | genotyping | ATGATAGCATCTTGTTCTTAGTCTTTTTCTTAATAGGG |
| EZH2 - forward | genotyping | CATGTGCAGCTTTCTGTTCA |
| EZH2 - reverse | genotyping | CACAGCCTTTCTGCTCACTG |
| Pb-Cre - forward | genotyping | TACAACTGCCAACTGGGATG |
| Pb-Cre - reverse | genotyping | AGGCAAATTTTGGTGTACGG |
| Pten - forward | genotyping | CAAGCACTCTGCGAACTGAG |
| Pten - reverse | genotyping | AAGTTTTTGAAGGCAAGATGC |
| Rb1 - forward | genotyping | GGCGTGTGCCATCAATG |
| Rb1 - reverse | genotyping | CTCAAGAGCTCAGACTCATGG |
